# Neural dynamics of phoneme sequences: Position-invariant code for content and order

**DOI:** 10.1101/2020.04.04.025684

**Authors:** Laura Gwilliams, Jean-Remi King, Alec Marantz, David Poeppel

## Abstract

Speech consists of a continuously-varying acoustic signal. Yet human listeners experience it as a sequence of discrete units (phonemes and words). To identify how the brain continuously represents and updates speech units, we recorded two-hour magnetoencephalograms from 21 subjects listening to short narratives. Our encoding and decoding analyses show that the human brain uses a dynamic coding scheme to represent approximately three successive phonetic features at each time sample. These dynamic phonetic representations, not directly accessible from the acoustic signal, are active earlier when phonemes are more predictable, and are sustained when lexical identity is uncertain. The results reveal how phoneme sequences in natural speech are neurally represented and shed new light on how the human brain combines them to probe the mental lexicon.

**One sentence summary:** The human brain keeps track of the relative order of speech sound sequences by jointly encoding content and elapsed processing time

---

Speech comprehension involves mapping variable continuous acoustic signals onto discrete linguistic representations [1]. Although the human experience is typically one of effortless understanding, uncovering the computational infrastructure underpinning speech processing remains a major challenge for neuroscience [2] and artificial intelligence systems [3, 4, 5] alike.

Existing cognitive models primarily explain the recognition of words in isolation [6, 7, 8]. Predictions of these models have gained empirical support in terms of neural encoding of phonetic features [9, 10, 11, 12], and interactions between phonetic and (sub)lexical units of representation [13, 14, 15, 16]. What is not well understood, however, is how *sequences* of acoustic-phonetic signals (e.g. the phonemes *k-a-t*) are assembled during comprehension of naturalistic continuous speech, in order to retrieve lexical items (e.g. *cat*).

Correctly parsing auditory input into phoneme sequences is computationally challenging for a number of reasons. First, there are no reliable cues for when each speech unit begins and ends (e.g. word or morpheme boundaries). Second, adjacent phonemes can acoustically blend into one another due to co-articulation [1]. Third, the same set of phonemes can form completely different words (e.g. *pets* versus *pest*), which makes the order of phonemes – not just their identity – critical for word recognition.

The neurophysiology of speech-evoked activity also poses practical challenges. Previous studies have shown that speech sounds elicit a complex hierarchy of brain responses, ranging from early auditory cortex to parietal and ultimately prefrontal areas [9, 17, 5]. Critically, the neural responses to an individual phoneme can be long-lasting and surpass the duration of the phoneme itself [18, 19, 11, 16]. This phenomenon implies that a given phoneme is still encoded in neural activity while subsequent phonemes stimulate the cochlea. Such a tangle of representations raises complicated questions regarding how invariance and perceptual constancy is achieved in spoken language comprehension.

To identify how the brain continuously represents, updates, and disentangles sequences of phonemes, we recorded 21 subjects with magneto-encephalography (MEG) while they attentively listened to 2 hours of spoken narratives. We then use a combination of encoding and decoding analyses time-locked to each phoneme onset to track how the brain represents past and present phonetic features.

Our results reveal how the language system (i) simultaneously processes acoustic-phonetic information of overlapping inputs; (ii) keeps track of the relative order of those inputs; and (iii) maintains information sufficiently long enough to interface with (sub)lexical representations. Our findings shed new light on how the brain orchestrates the overlapping neural processes elicited by continuous speech, without confusing either their content or ordering.

## 1. Results

Neural responses were recorded with magnetoencephalography (MEG) while 21 participants listened to four short stories recorded using synthesised voices from Mac OS text-to-speech. Each subject completed two one-hour recording sessions, yielding brain responses to 50,518 phonemes, 13,798 words and 1,108 sentences per subject. We annotated each phoneme in the story for its timing, location in the word, as well as phonetic and statistical properties (Figure 1A).

**Figure 1:**
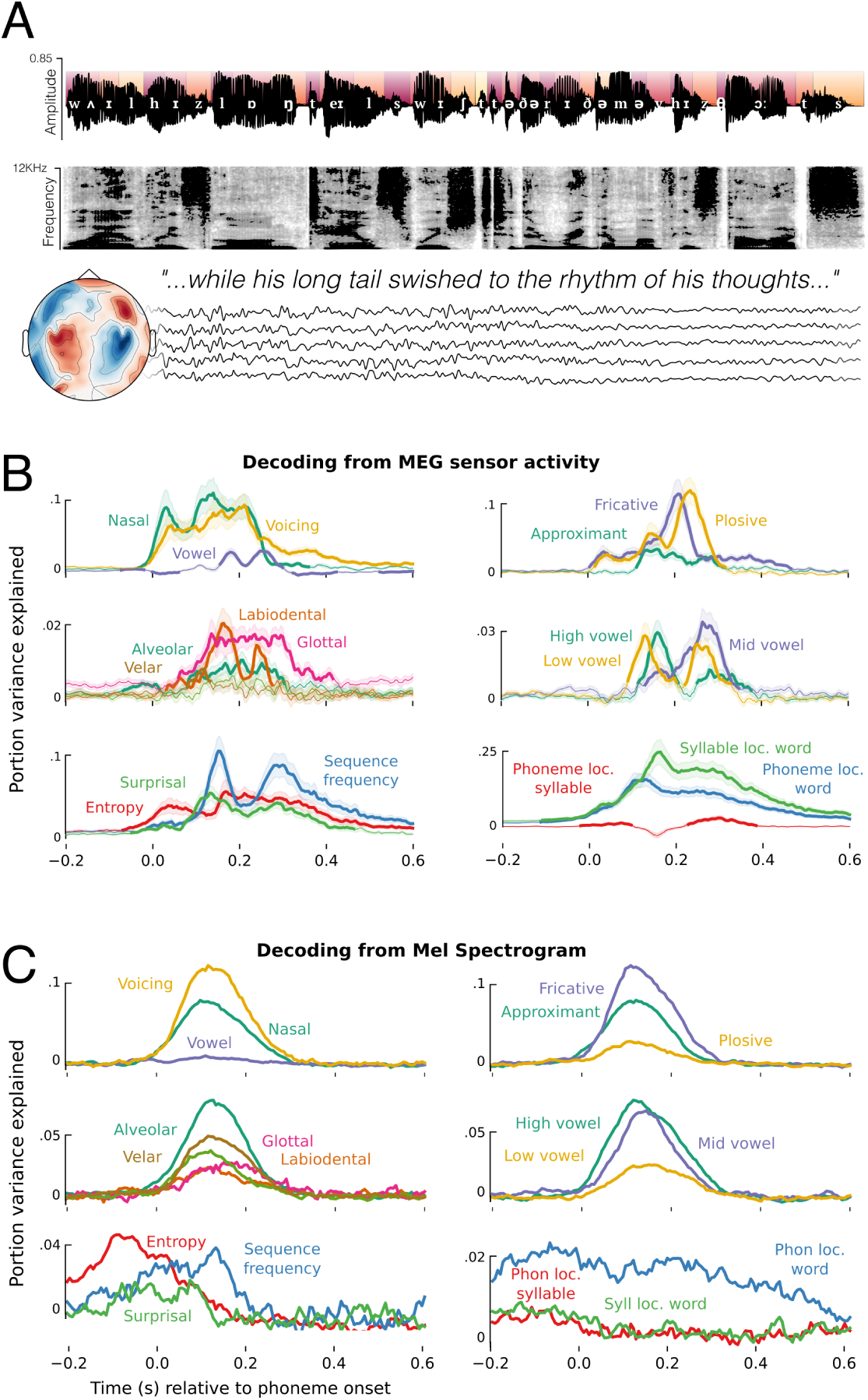
Experimental design and phoneme-level decoding. A: Example sentence, with the phonological annotation superimposed on the acoustic waveform. Colours of the segments at the top half of the waveform indicate the phoneme’s distance from the beginning of a word (darker red closer to word onset). The spectrogram of the same sentence appears below the waveform. Five example sensor residual time-courses are shown below, indicating that all recordings of the continuous stories were recorded with concurrent MEG. B: Timecourses of decoding phoneme-level features from the MEG signal using back-to-back regression, time-locked to phoneme onset. C: Timecourses of decoding the same features from the mel spectrogram. Traces represents the average decoding performance over subjects, shading represents standard error of the mean. The y-axis represents portion of explainable variance given all features in the model.

### 1.1. Phonetic feature encoding in acoustic and neural signals

Are phonetic features linearly represented in the brain activity? To address this issue, we first trained a temporal receptive field (TRF) model to predict MEG data from the non-linguistic variability in the acoustics, namely the amplitude envelope and the pitch contour of the speech signal [20]. We used the predictions of this model to regress out low-level auditory responses from the MEG signal. All subsequent analyses were applied to the residuals of this model, and so can be interpreted with the potential confounds of loudness and pitch being removed.

To track the neural representations of linguistic features, we then fit a back-to-back regression (B2B) [21] to (1) decode 31 linguistic features from the residual MEG signal and (2) evaluate their actual contribution to the corresponding decoding performance. This was applied at each time sample independently. These phonetic features included 14 acoustic-phonetic features (binary one-hot encoding of *place, manner* and *voicing*), 3 information theoretic measures (surprisal, entropy, and log sequence frequency), 8 unit boundary features (onset/offset of words, syllables, morphemes) and 6 positional features (phoneme and syllable location in syllable, word, and sentence). All features were included in the model simultaneously. B2B was chosen to control for the co-variance between features in the multivariate analysis while optimizing the linear combination of MEG channels to detect the encoding of information even in low signal-to-noise circumstances [21]. We applied the same model separately to the MEG signals (n=208 channels) and to the acoustic signal (n=208 frequency bands of the acoustic mel spectrogram, which approximates the auditory signal after passing through the cochlea [22]).

The output of the B2B is a set of beta coefficients that map a linear combination of ground-truth features to the decoder’s predictions of a feature. For explicit comparison between the results using more classic decoding metrics (AUC) and B2B regression coefficients, see Supplementary Figure 6. For reference, the average significant decoding accuracy is modest, at around 51-52% *but* given the number of phonemes presented to the subjects, it is highly significant: all *p* < 10^−4^. Here the effect sizes we report are the portion of unique variance explained by each feature, relative to a noise ceiling of total explainable variance.

As an important theoretical point is that we are decoding features of each phoneme rather than decoding phoneme categories per se. However, when decoding phoneme categories instead of features, we observe very similar results (see supplementary Figure 19).

Figure 1B summarises the result of the neural analysis. Confirming previous studies [9, 11, 16], phonetic features were decodable on average between 50-300 ms from the MEG signal and accounted for 46.2% of variance explained by the full suite of 31 features. Performance averaged across features was statistically greater than chance, as confirmed with a temporal permutation cluster test, based upon a one-sample t-test (df=20) applied at each time-sample (*p* < .001; critical 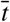 averaged over time = 3.61). Information theoretic measures such as surprisal, entropy, and log sequence frequency accounted for 14.5% of explainable variance, which was also statistically better than chance on average (−10-540 ms; *p* < .001; 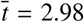). Positional properties such as location in the word explained 31% of variance, and showed the greatest individual effect sizes (−120-600 ms; *p* < .001; 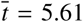). Here, phoneme order is coded the same for all phonemes regardless of feature. Finding a significant effect of phoneme order therefore suggests that order is represented independently from phonemic content, and is the same for all phonemes regardless of the features they contain. The remaining variance was accounted for by boundary onset/offset features (0-410 ms; *p* < .001; 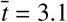).

Are phonetic feature representations in the neural signal similar to those of the auditory input? Figure 1C shows the result of the same B2B analysis, but now applied to the audio mel spectrogram. Phonetic features accounted for 52.1% of the explainable variance between 0-280 ms (*p* < .001; 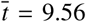). We also observed correlates of statistical structure (−180-140 ms; *p* < .01; 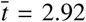) and phoneme position (−200-580 ms; *p* < .001; 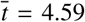). Phonetic features that were better decoded from the acoustic signal were also better decoded from the neural signal (Spearman correlation of average performance *r* = .59; *p* = .032). However, there was no significant correlation between the decoding performance of statistical structure and phoneme position across the auditory and neural analyses (Spearman *r* = .13; *p* = .41), and these features accounted for significantly more variance in the MEG as compared to the auditory analysis (*t* = 2.82; *p* = .012). Overall, this analysis suggests that phonetic and statistical features of speech have correlates in the acoustic signal, but that the decoding of higher-order structure from neural responses cannot be explained by the acoustic input alone.

### 1.2. Multiple phonemes are processed simultaneously

The average duration of phonetic decodability (310 ms) contrasts with the average duration of the phonemes in our stories (78 ms, SD = 34 ms). This discrepancy suggests that the brain processes multiple phonemes concurrently. To evaluate how many phonemes can be simultaneously decoded at each time sample, we decoded, from the same MEG responses, the place, manner, and voicing of the current and three preceding phonemes. We find that a history of three phonemes can be robustly decoded from an instantaneous neural response (see Supplementary Figure 11).

To assess whether this result implies that the brain can retrieve the content and order of successive phonemes, we simulated the neural responses to 4-phoneme anagrams using the decoding model coefficients to reconstruct synthesised MEG responses (see Methods). Under ideal conditions (no added noise) neural responses robustly encoded the history of four preceding phonemes (Figure 2B). This result confirms the high degree of overlapping content during phoneme processing. Although decodability need not imply that the brain actually *uses* this information [23], this result indicates that the brain has access to *at least* a 3-phoneme sequence at any given processing time.

**Figure 2:**
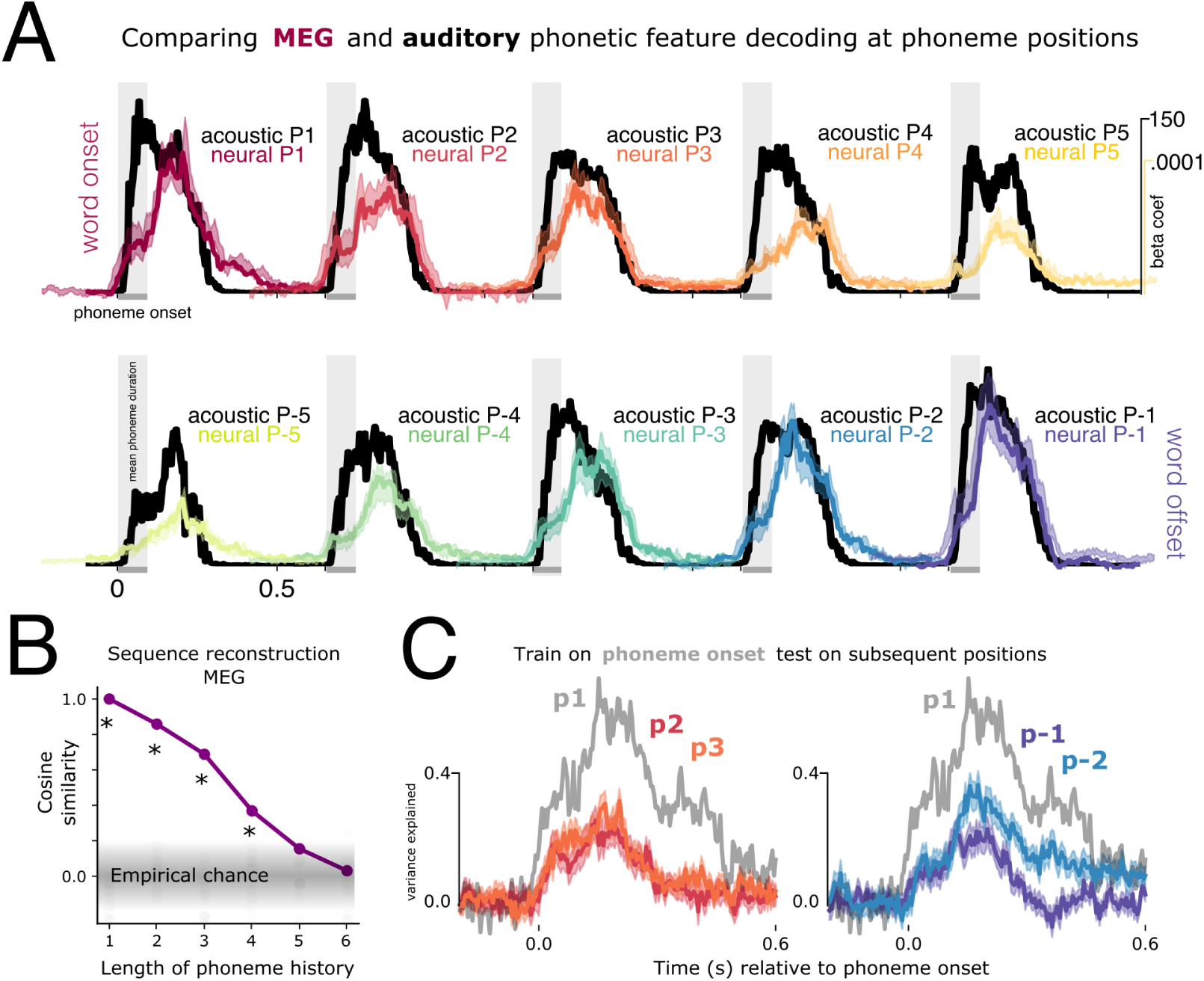
Decoding as a function of phoneme position. A: Timecourses of decoding phoneme-level features from the mel spectrogram (black) and MEG sensors (coloured), time-locked to phoneme onset. Performance is averaged over the 14 phonetic feature dimensions. Shading in the neural data corresponds to the standard error of the mean across subjects. Results are plotted separately for 10 different phoneme positions, where P1:P5 indicates distance from word onset (red colour scale) and P-1:P-5 distance from word offset (purple colour scale). E.g. P-1 is the last phoneme of the word, P-2 is the second to last phoneme of the word. Y-axis shows the raw beta coefficients before noise-ceiling correction. All plots share the same y-axis scales, which are different for neural and acoustic analyses (top right). B: Result of simulating and reconstructing phoneme sequences from the MEG model coefficients. X-axis represents how many previous phonemes can be reconstructed; y-axis represents the cosine similarity between the true phoneme sequence and the reconstruction. C: Testing generalisation across phoneme positions. Grey trace shows average performance when training on phonemes at word onset and testing on phonemes at word onset. Coloured traces show average performance when training on phonemes at word onset, and testing at second (P2), third (P3), last (P-1) and second to last (P-2) phoneme positions. Y-axis is the noise-normalised variance explained within the generalisation analysis.

### 1.3. Representations are shared across phoneme positions: invariance

How does the brain represent these multiple phonetic features without confusing their identity? One computational solution is position-specific encoding, which posits that the representation of each phoneme is different depending on where it occurs in a word. This coding scheme avoids representational overlap by having a specific coding scheme for first phoneme position (P1), second phoneme position (P2), etc.

To test the hypothesis that the brain uses this coding scheme, we trained a decoder on the responses to phonemes in first position and evaluated the model’s generalisation to subsequent phoneme positions (Figure 2C). We could read out the features of P2, P3, P-1 and P-2 from 20-270 ms (*p* < .001; 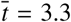), with comparable performance (max variance for P2 = 26%, SEM = 4%; P3 = 32%, SEM = 3%; P-1 = 23%, SEM = 3%, P-2 = 37%, SEM = 4%). This result supports the existence of a position-*invariant* representation of phonetic features and therefore contradicts a purely position-specific encoding scheme.

Interestingly, training and testing on the same phoneme position (P1) yielded the strongest decodability (max = 71%, SEM = 5%), which was significantly stronger than when generalising across positions (e.g. train P1 test P1 vs. train on P1 test on P2: 110:310 ms, p = .006). It is unclear whether this gain in performance is indicative of position-specific encoding *in addition to* invariant encoding, or whether it reflects bolstered similarity between train and test signals due to matching other distributional features. Future studies could seek to match extraneous sources of variance across phoneme positions to test this explicitly.

### 1.4. Phonetic representations rapidly evolve over time

The sustained and position-invariant representation of successive phonemes is at odds with subjects’ ability to recognise words and thus leaves our original question unanswered: how does the brain prevent interference between successive phonemic content?

To address this issue, we used temporal generalisation (TG) analysis [24]. This method consists of learning the most informative spatial pattern for a phonetic feature at a given moment, and assessing the extent to which it remains stable over time. TG results in a training × testing matrix that helps reveal how a neural representation evolves over time.

To evaluate the extent of representational overlap across phoneme positions, we aligned the TG matrices relative to the average latency between two adjacent phonemes. For example, relative to word onset, phoneme P1 is plotted at t=0, P2 at t=80, P3 at t=160, and so on (Figure 3). Note that the number of trials at test time decreases as we test phoneme positions further from word boundaries, due to differences in word length. P1 and P-1 are defined for 7335 trials per subject per repetition, P2 and P-2 for 7332 trials, P3 and P-3 for 5302 trials, P4 and P-4 for 3009 trials, P5 and P-5 for 1744 trials. Models are trained on all phonemes regardless of position (25259 per subject per repetition). This encourages the model to learn the topographic pattern which is common across phoneme positions.

**Figure 3:**
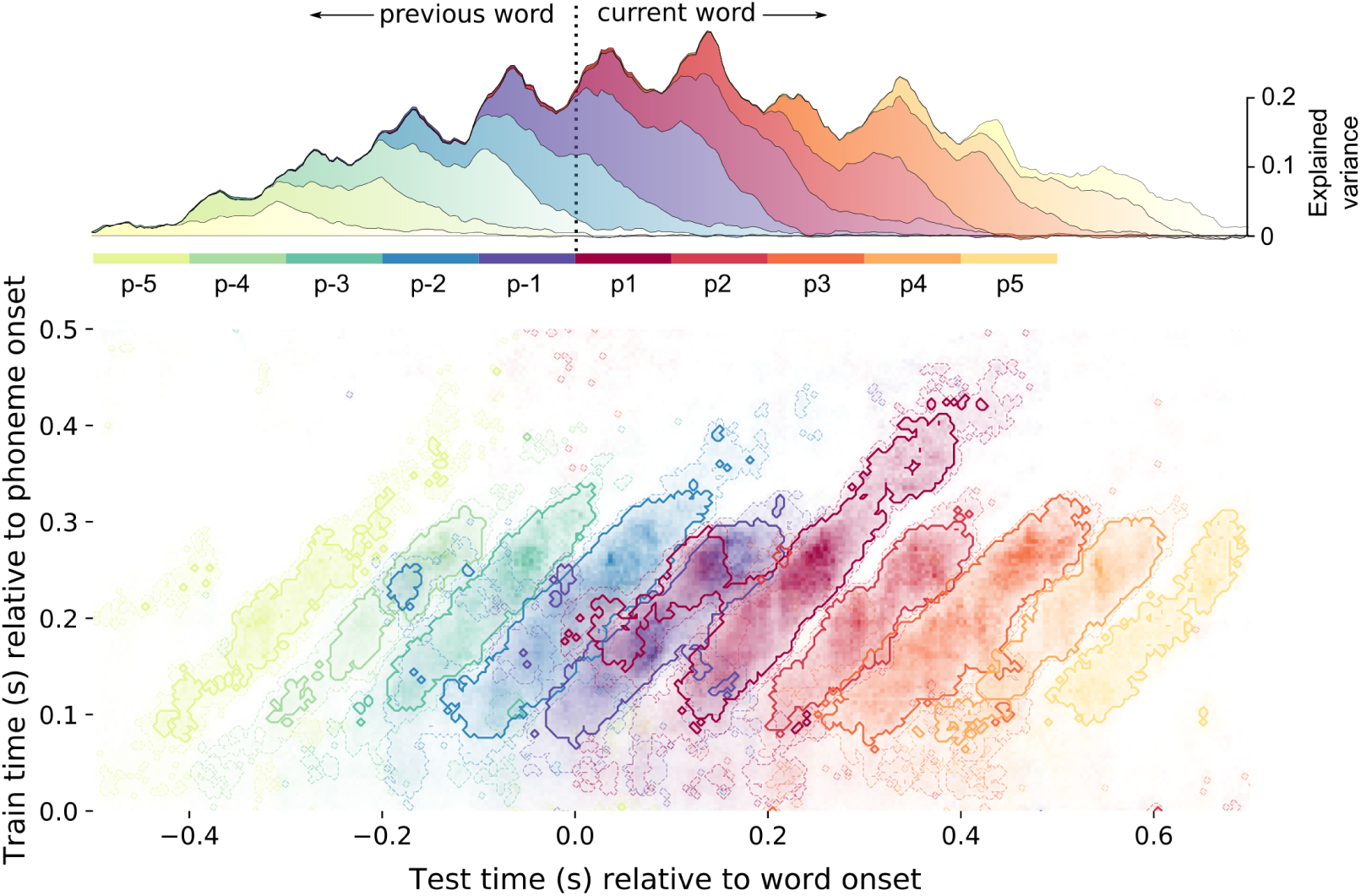
Phonetic feature processing across the sequence. Temporal generalisation (TG) results superimposed for 10 phoneme positions. Results represent decoding performance averaged over the 14 phonetic features. From the first phoneme in a word (P1, dark red) moving forwards, and the last phoneme in a word (P-1, dark blue) moving backwards. The result for each phoneme position is shifted by the average duration from one phoneme to the next. The y-axis corresponds to the time that the decoder was trained, relative to phoneme onset. The x-axis corresponds to the time that the decoder was tested, relative to (superimposed) word onset. Contours represent a t-value threshold of 4 (darker lines) and 3.5 (lighter lines). Note that fewer trials are entered into the analysis at later phoneme positions because words contain different numbers of phonemes. This is what leads to the reduction of decoding strength at p5 and p-5, for example. The upper panel shows cumulative decoding performance of just the diagonal plane, representing when phonetic information is available, regardless of the topography which encodes it.

While each phonetic feature could be detected for around 300 ms (train time == test time, ‘diagonal axis’, Figure 3 lower panel), the corresponding MEG patterns trained at each time sample were only informative for about 80 ms (marginal over train time, ‘horizontal axis’, Figure 3 lower panel). This discrepancy demonstrates that the neural representations of phonetic features quickly change over time. We confirmed the statistical reliability of this evolution using an independent samples t-test, comparing diagonal and horizontal decoding performance axes (df = 200, *p* < .001; 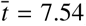). Together, these results suggest that the brain uses a dynamic coding scheme to simultaneously represent successive phonemes.

### 1.5. Dynamic coding disentangles successive phonetic representations

To further test whether this dynamic coding scheme minimises the overlap between neighbouring representations, we quantified the extent to which successive phonemes were simultaneously represented in the same neural assemblies - i.e. by the same MEG decoders. To this end, we extracted all time samples across train and test time where decoding performance exceeded a *p* < .05 threshold, Bonferroni-corrected across the 201 time-samples of a single processing time. This is equivalent to a statistical *t* threshold > 4, an effect size threshold variance explained > 5%, and decoding AUC > .507, as represented by the the darker contour on Figure 3. We then compared the number of significant time-points shared within and across phoneme positions. On average, 7.3% of time-points overlapped across phoneme positions (SD = 9%). The majority of overlap was caused by the first phoneme position, which shared 20% of its significant time-samples with the preceding phonemes, and reciprocally, the last phoneme which shared 27.9% of its time-samples with the subsequent (as can be seen in the left-sided ‘appendage’ that overlaps with the last phoneme of the previous word). When removing the first and last phonemes from the analysis, this overlap drops to just 3.1% (SD = 3.2%).

Overall, these results show that while multiple phonemes are processed in parallel, any given pattern of neural activity primarily represents one phoneme at a time, granting each phoneme an individuated representation.

### 1.6. Dynamic representations are absent from audio spectrograms

Is this dynamic coding scheme also present in the auditory signal, and thus trivially explains the brain dynamics? To address this issue, we applied the TG analysis to the audio mel spectrogram. We found no statistical difference between the accuracy time-course of a single decoder as compared to independent decoders at each time sample (*p* = .51; 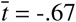). The ‘square’ shape of the TG matrix suggests that acoustic signals contain stationary cues for acoustic-phonetic features (see Supplementary Figure 7). We also applied the overlap analysis to the auditory signal, using the same statistical threshold to identify relevant time-samples. On average, 92.5% of time-samples overlapped across positions (SD = 12.3%). This remained similar when removing the first and last phonemes from the analysis (mean = 91%, SD = 13.3%). Finally, we also confirmed that this overlap was qualitatively similar in different signal-to-noise ratio simulations (see Supplementary Figure 9).

These results demonstrate that the dynamics we observe in the neural signal are not present in the acoustic signal. There was significantly more representational overlap in the auditory encoding as compared to the neural encoding (*t* = −21.3, *p* < .001), suggesting that the dynamic coding scheme we observe in neural responses is not a straightforward reflection of the acoustic input. Rather, it is the consequence of an active process the brain applies to the relatively static auditory representations.

### 1.7. Dynamic coding automatically recovers the order of successive phonemes

The dynamic coding scheme we show above may implicitly encode the order of each representation in the sequence [25]: Given that the representation of each phoneme evolves over time, each phoneme is continuously ‘time-stamped’. To formally test this possibility, we cropped all phoneme-aligned responses between 100-400 ms and used ridge regression to decode the latency since phoneme onset (see Methods). The reconstructed latency significantly correlated with the true latency (Pearson r = 0.87, p < .001; see Figure 4A), suggesting that the dynamic spatial topography also encodes the time elapsed since phoneme onset. This time-stamp thus distinguishes sequences composed of the same phonemes such as ‘pets’ versus ‘pest’.

**Figure 4:**
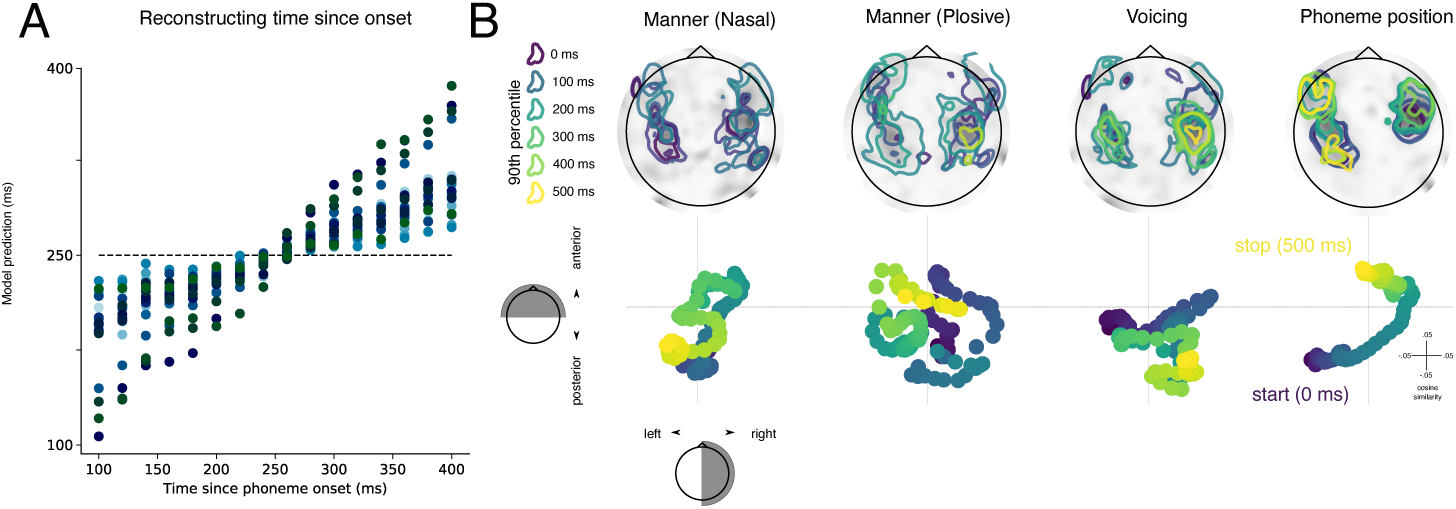
Spatial evolution of processing trajectory. A: Reconstructing time since phoneme onset from 15 equally spaced samples from 100-400 ms. Horizontal dashed line indicates mean of time latency (i.e. what would be reconstructed in absence of evidence to the contrary). Each coloured dot corresponds to one subject. B: Decoding model coefficients plotted as topographies (above) and trajectories (below) over time. Coefficients are shown for the decoding of three phonetic features and phoneme position. Phoneme onset is shown in the darkest purple (0ms) and colours shift towards yellow as a function of time. Contours on the topographies highlight 90% percentile coefficient magnitude at 100 ms increments. Trajectories below represent the cosine distance between the coefficient topography and a binary posterior/anterior mask (y-axis) and a binary left/right mask (x-axis). Overall the results show that the topographies evolve over time, with phonetic features being concentrated around auditory cortex, and phoneme position taking an anterior path from auditory to frontal cortices.

### 1.8. MEG topographies associated with phonetic representations

To clarify the spatial underpinning of these dynamic representations, we analysed the beta coefficients of the decoding model when withholding regularisation (see Methods). Figure 4B displays the temporal evolution of the MEG topographies associated with the most robustly encoded phonetic features. We quantified the extent of spatial evolution over time by projecting the coefficients onto orthogonal axes of the sensor array (front/back, left/right) and computing the extent of temporal structure (see Methods for details).

Overall, we find that informative activity remained local within auditory regions, in contrast to the representation of phoneme position which followed a clear posterior-anterior gradient. Given that the brain has simultaneous access to phoneme content and elapsed processing time, the coupling of these trajectories permits simultaneous read-out of both the *identity* and the *relative order* of a speech sound in a sequence from any given spatial pattern of activity. These neural patterns encoding phonetic features exhibited significantly more structured movement over time than a null trajectory (*p* < .001).

### 1.9. Representations evolve at the rate of phoneme duration

The representational trajectories (Figure 3) complete a full evolution cycle every ∼80 ms (i.e. average phoneme duration). Next, we sought to test whether (i) the cycle always unfolds at a fixed rate, which entails that different numbers of phonemes are maintained depending on their length or (ii) the rate is faster for shorter phonemes and slower for longer phonemes, which means that the same number of phonemes are always maintained, regardless of their length.

We grouped trials into quartiles and analysed brain responses to the shortest and longest phonemes (∼4500 trials in each bin; mean duration 45 and 135 ms, respectively). Phoneme duration predicted the duration of temporal generalisation across training time: longer phonemes generalised for an average 56 ms longer than shorter phonemes (*p* = .005; 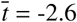) (Figure 5A).

**Figure 5:**
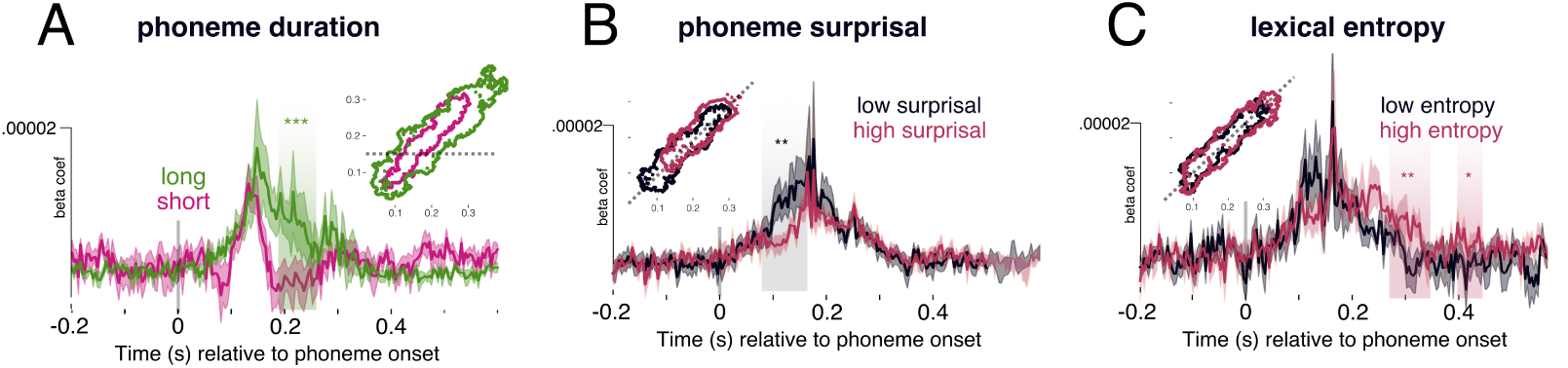
Manipulating axes of sequence dynamics. We find that different properties of the speech input modulate ‘finger’ latency, width and angle. A: TG analysis median split into short (average 45 ms) and long (average 135 ms) phonemes. Contour inlay represents the borders of the significant clusters at *p* < .001. Waveforms represent a horizontal slice at 140 ms (shown as a dashed line in the contour plot). B: Analysis on all non-onset phonemes split into median surprisal, along the diagonal plane (slice shown in contour plot). Highlighted areas show significant temporal clusters between low and high surprisal. C: Analysis on all non-onset phonemes split into median cohort entropy, also along the diagonal plane. Highlighted areas show significant temporal clusters between low and high entropy. Shading on the waveform of all plots represents standard error of the mean across subjects. * = *p* < .05; ** = *p* < .01; *** = *p* < .001.

In addition, we observed a small but robust deflection in the angle of the trajectory ‘finger’. If a processing trajectory moves faster than the training data, it will have an angle of less than 45 degrees. Slower trajectories will have an angle greater than 45 degrees. Using principal component analysis to find the axis of greatest variance, we found that shortest phonemes had an average angle of 42.3 degrees, whereas the longest phonemes had an average angle of 47.1. This difference was statistically significant when applying an independent samples t-test between short and long phonemes across subjects (df = 20, *t* = 2.56, *p* = .013).

This pattern suggests that the brain adapts its speed of processing as a function of the rate of speech input, to ensure that approximately the same number of phonemes are encoded at a given test time while minimising representational overlap. This potentially supports the conjecture that the brain preferentially derives higher order structure by a computing sliding triphone sequences from the speech input.

### 1.10. Phonetic representations vary with linguistic processing

To what extent do these neural representations of phonemes relate to linguistic processing? To address this question, we next tested whether these phonetic representations systematically vary with (i) word boundaries, (ii) surprisal, and (iii) lexical identification.

#### 1.10.1. Phonetic processing is modulated at word boundaries

We evaluated decoding performance at word boundaries: word onset (position P1) and word offset (position P-1) separately for each family of phonetic features (place of articulation, manner, and voicing). Phonetic features were decodable earlier at word onset then offset, yielding a significant difference during the first 250 ms (place: *p* = .03, 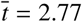, 84-112 ms; *p* < .001, 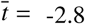, 156-240 ms; manner *p* < .001, 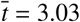, 72-196 ms; *p* = .004, 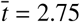, 220-300ms). When averaging decoding performance across phonetic features, the lag between neural and acoustic maximum accuracy was 136 ms (SD = 13 ms) at word onset, which reduced to 4 ms (SD = 13 ms) at word offset (see Figure 2A). This average onset/offset latency difference was statistically significant (*t* = −3.08; *p* = .002). Furthermore, place and voicing features were sustained in the neural signal significantly longer for phonemes at the beginning of words as compared to the end (place: *p* = .009, 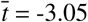, 302-418 ms; voicing: *p* < .001, 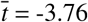, 328-428 ms). This was also true when averaging over all features (*p* < .001, 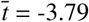, 328-396 ms) (see Figure 3).

#### 1.10.2. Predictable phonemes are processed earlier

Is this latency shift purely lexical in nature, or could statistical properties of speech sounds at word boundaries explain the result? In general, and in our speech stimuli, phonemes at the beginning of words are less predictable than phonemes at the end of words. We tested whether expected phonemes could be decoded earlier from the neural signal than unexpected ones by grouping trials as a function of surprisal. To control for co-linearity between word-boundaries and surprisal, we selected all phonemes that were *not* at word onset, and tested decoding accuracy relative to quartile phoneme surprisal (see Methods for details on how this variable was computed). Each analysis bin contained ∼4500 trials. If the shift in latency is purely a word-boundary effect, we should observe no latency difference as a function of non-onset phoneme surprisal.

We observed a systematic latency shift as a function of surprisal (Figure 5B): more predictable phonemes were decoded earlier than less predictable ones, leading to a significant difference between low and high surprisal from 120-132 ms (*p* = .007). Surprisal did not significantly modulate peak decoding accuracy (all uncorrected *p*-values > .2).

This result suggests that the brain initiates phonetic processing earlier when the phoneme identity is more certain. This result at least partially, and potentially fully, explains the latency shift at word boundaries.

#### 1.10.3. Phonetic features are maintained until lexical identification

Our results suggest that the temporal dynamics of phonetic processing are modulated by certainty about the phoneme unit being said. Does this extend to certainty about word identity, such that phonetic representations are maintained until higher order structure can be formed?

We quantified word uncertainty using lexical cohort entropy [26]. If many words are compatible with a given phoneme sequence, then lexical entropy is high. On the contrary, if only one word is likely given a specific phoneme sequence (which is more often the case towards word offset), then lexical entropy is low. We evaluated whether decoding performance between 300-420 ms (the window that showed the word onset/offset effect) varied with lexical entropy (Figure 5C). We grouped trials based on lexical entropy, and ran the analysis on all phonemes that did *not* occur at word onset (∼4500 trials per bin). In the window of interest, higher entropy phonemes were decoded with significantly higher performance 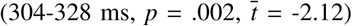. This suggests that phonetic information is maintained for longer in cases of higher lexical uncertainty. When we expand the analysis window to the entire 0-500 ms epoch with a lower cluster-forming threshold, we find that lexical entropy significantly modulates phonetic decoding from 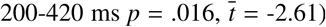. This is a reasonable time-frame within which lexical entropy could exhibit an effect, and matches the time-frame of entropy reduction in our stimulus materials remarkably well. The average entropy of the low-entropy bin was 1.19 bits, which on average occurs 333 ms into a word (SD=158 ms). The average entropy of the high-entropy bin was 4.3 bits, which on average occurs 108 ms into a word (SD=51 ms). This therefore means that the average time between a phoneme in the high entropy bin and low entropy bin was 225 ms, which matches the duration of the entropy modulation effect in our data (220 ms). This in turn suggests that phonetic detail is maintained until lexical ambiguity is sufficiently reduced.

In a separate analysis, we tested whether word length played a role in the maintenance of phonetic detail of the onset phoneme. We compared decoding performance of word-onset phonemes grouped into median word length (shorter mean length = 2.54 phonemes; 4058 trials; longer mean length = 5.2 phonemes; 2841 trials). No significant differences between groups were found (all clusters *p* > .2, duration < 2 ms).

Overall, our results confirm that the neural representations of phonemes systematically vary with lexical features.

## 2. Discussion

We analysed MEG responses to continuous speech as a function of phonetic features and position in the phoneme sequence. Our results show that the brain maintains phonetic representations of the three most recently heard phonemes. These representations are invariant to position, that is, the representation of /b/ is the same whether that phoneme occurs early or later in a word. They follow a common ‘processing trajectory’ over auditory and frontal cortices bilaterally, as the representations evolve over time. This predictable temporal evolution encodes the time elapsed since phoneme onset, and distinguishes the most recent phoneme representation from the second and third most recent. Importantly, we find that the speed of this processing trajectory is dynamic. It evolves at the rate of phoneme duration (slower for longer phonemes, faster for shorter phonemes), consequently preventing representational overlap between neighbouring speech sounds. Further, it begins earlier for predictable phonemes, and continues longer when lexical uncertainty is higher. Taken together, our results shed new light on how the human brain continuously transcribes the stream of auditory speech sounds it receives into a sequence of speech units.

The striking contrast between the stationarity of the audio spectrogram *versus* the dynamics of the corresponding neural representations highlights that the neural representations of phonemes are more than a mere reflection of the acoustic signals [27]. Our simulations further demonstrate that these dynamic representations permit the auditory system access to the history of multiple (at least three - our lower bound estimate) phonemes simultaneously, without blending them within a common activity pattern. This coding scheme grants two computational advantages. First, it serves to avoid interference between phonetic units, by ensuring an orthogonal format in processing space. This first property provides a clear answer to how speech sequences are maintained without merging their contents into unusable representations. Second, relative order is implicitly coded in the pattern of activity that encodes phonetic detail at variable latencies relative to phoneme onset. In other words, the relative ‘location’ along the processing trajectory provides the brain with ordinal sequence information. The encoding of phoneme content regardless of order, and phoneme order regardless of content, is what allows the brain to keep track of speech sound sequences, i.e. to know that you were asked to *teach* and not *cheat*, or that you are eating *melons* and not *lemons*.

Building on this observation, we found that the representational trajectory is consistent across phoneme positions, leading to significant generalisation from one phoneme position to another, with comparable magnitude. This finding rules out a number of competing hypotheses. First, it is difficult to reconcile these results with an explicit sequence representation. For example, if the brain represents a sequence of all elapsed phonemes, the representation of phoneme *X* at word onset would generalise poorly to third position *ABX* and even worse to sixth position *ABCDEX*. Second, under the same logic, this result rules out the idea that phonemes have a purely context-dependent encoding scheme, such as being represented along with their co-articulatory neighbours [28]. In that case, phoneme *X* would have a different representation in the context *AXB* and *VXY*. Finally, generalisability is inconsistent with position-specific encoding accounts, such as edge-based schemes [29, 30], which would posit that *X* is encoded differently in *ABX* and *XBC*. While it is possible that multiple representational systems co-exist, our results support that at least one of those encoding schemes is context-independent, which encodes content regardless of lexical edges.

When testing the spatial evolution of information flow, we found that phonetic features are encoded within auditory regions for around 100-400 ms, and engage in complex interactions with frontal areas starting about 200 ms. The trajectory was different across features and did not trace a clear spatial path, suggesting that information may be locally changing within the auditory cortices rather than following a strict anatomical transposition from low to high level areas [12, 5, 25]. In contrast, the explicit encoding of phoneme position *did* follow a clear spatial trajectory, moving toward the anterior parts of the auditory regions. Unfortunately the signal strength of our single trial estimates, and the spatial resolution of MEG, limits the specificity of the spatial claims we can make based on these data. Future work will thus be critical to clarify the exact spatio-temporal profile of this processing trajectory.

Given that each phonetic feature appears to follow a common representational trajectory, how can we describe the transformational space? One possibility links to articulatory phonology [31]. Although we showed the input acoustic space to be (surprisingly) static, the articulatory gestures which produce those acoustics are inherently dynamic [32]. It is plausible, therefore, that speech sounds are processed via articulatory commands, which are believed to jointly encode both the sound being produced and the temporal delay relative to articulatory onset. This idea of joint content-temporal coding resonates with recent findings of sequence encoding – finding evidence for dedicated temporal codes in rat hippocampus [33] or in the human visual system [25]. Future work will need to further delineate the temporal code used for phonological processing, and how the spatial location of these responses changes as a function of processing time.

Several elements suggest that the present neural representations of phonemes directly relate to high-level linguistic processing. First, the representational trajectory gets systematically delayed as a function of phonological uncertainty (surprisal) and systematically sustained as a function of lexical uncertainty (cohort entropy). This suggests that the language system continuously adapts its processes based on information across multiple levels of linguistic description simultaneously. Such latency shifts fit with models of predictive coding [34, 35] and analysis-by-synthesis [36]: when predictability for a phoneme is strong, processes can be initiated earlier (perhaps in some cases before the sensory input) than when the phoneme identity is unknown. Although previous work has shown that processing of the speech signal is sensitive to phoneme probability within a word [37, 13, 38, 15, 39] (see [26] for a review), this is the first study quantifying the *consequences* this has for encoding the content of those phonemes. Interestingly, we did not observe an effect of predictability on overall decoding performance, suggesting that processing delays may serve as a compensatory mechanism to allow more information to be accumulated in order to reach the same strength of encoding [40]. Future work should test whether this local (within-word) predictability metric has similar consequences to global (across-word) metrics.

Second, phonetic features are maintained longer during lexical ambiguity. While previous work has shown active maintenance of phonetic information [16], to our knowledge, this is the first evidence that maintenance is dynamic and scales with certainty over higher-order units. This result not only highlights the adaptability of speech processing but also demonstrates the remarkable bi-directional interaction between hierarchical levels of processing. Our results suggest that acoustic-phonetic information is maintained until the (sub)lexical identity meets a statistically-defined threshold, providing clear processing advantages in the face of phonological ambiguity and lexical revision [16]. Future work would benefit from orthogonalizing lexical entropy and word-internal disambiguation points (i.e. when entropy reduces to zero) to more precisely quantify the temporal relationship with information maintenance.

Our study has several shortcomings that can be improved upon in future work. First, we chose a passive listening paradigm, in order to optimise the number of trials we could obtain within a single recording session. However, this prohibits us from associating decoding performance with behaviour and task performance. Second, using the non-invasive recording technique of MEG allows us to conduct long recording sessions in healthy individuals. However, the spatial resolution and the signal-to-noise ratio of MEG remain insufficient to clearly delineate the spatial underpinning of the dynamic encoding scheme. This limited signal-to-noise ratio also explains the relatively low decoding performances - peaking only 1-2% greater than chance in most cases. Future work may benefit from recording a greater number of repetitions of fewer speech sounds to allow for signal aggregation across trials.

Our results reveal that the brain implements an extremely elegant computational solution to the processing of rapid, overlapping phoneme sequences: the phonetic content of the unfolding speech signal is jointly encoded with elapsed processing time, thus representing content and order without relying on a position-specific coding scheme. The temporal dynamics of these computations are strikingly flexible, and vary as a function of certainty across both phonemic and lexical unit identity. These findings provide a critical piece of the puzzle for understanding how the human brain continuously parses and represents speech input to access the mental lexicon.

## 3. Method

### 3.1. Participants

Twenty-one native English participants were recruited from the NYU Abu Dhabi community (13 female; age: M=24.8, SD=6.4). All provided their informed consent and were compensated for their time. Participants reported having normal hearing and no history of neurological disorders. Each subject participated in the experiment twice. Time between sessions ranged from 1 day to 2 months. The experiment was approved by the IRB ethics committee at New York University Abu Dhabi.

### 3.2. Stimulus development

Four fictional stories were selected from the Open American National Corpus [41]: Cable spool boy (about two bothers playing in the woods); LW1 (sci-fi story about an alien spaceship trying to find its way home); Black willow (about an author struggling with writer’s block); Easy money (about two old friends using magic to make money).

Stimuli were annotated for phoneme boundaries and labels using the ‘gentle aligner’ from the Python module *lowerquality*. Prior testing provided better results for *lowerquality* than the Penn Forced Aligner [42].

Each of the stories were synthesized using the Mac OSX text-to-speech application. Three synthetic voices were used (Ava, Samantha, Allison). Voices changed every 5-20 sentences. The speech rate of the voices ranged from 145-205 words per minute, which also changed every 5-20 sentences. The silence between sentences randomly varied between 0-1000 ms.

### 3.3. Procedure

Stimuli were presented binaurally to participants though tube earphones (Aero Technologies), at a mean level of 70 dB SPL. The stories ranged from 8-25 minutes, with a total running time of ∼1 hour. Before the experiment proper, every participant was exposed to 20 seconds of each speaker explaining the structure of the experiment. This was designed to help the participants attune to the synthetic voices.

The order of stories was fully crossed using a Latin-square design. Participants heard the stories in the same order during both the first and second sessions.

Participants answered a two-choice question on the story content every ∼3 minutes. For example, one of the questions was ‘what was the location of the bank that they robbed’? The purpose of the questions was to keep participants attentive as well as to have a formal measure of engagement. Participants responded with a button press. All participants performed this task at ceiling, with an accuracy of 98%.

### 3.4. MEG acquisition

Marker coils were placed at five positions to localize each participant’s skull relative to the sensors. These marker measurements were recorded just before and after the experiment in order to track the degree of movement during the recording.

MEG data were recorded continuously, using a 208 channel axial gradiometer system (Kanazawa Institute of Technology, Kanazawa, Japan), with a sampling rate of 1000 Hz and an online low-pass filter of 200 Hz and a high pass filter of 0.01 Hz.

### 3.5. Preprocessing MEG

The raw MEG data were noise reduced using the Continuously Adjusted Least Squares Method (CALM: (Adachi et al., 2001)), with MEG160 software (Yokohawa Electric Corporation and Eagle Technology Corporation, Tokyo, Japan).

We used a temporal receptive field (TRF) model to regress from the raw MEG data responses that were sensitive to fluctuations in the pitch and envelope of the acoustic speech signal. We used the *ReceptiveField* function from MNE-Python [43], using ridge regression as the estimator. We tested ten lambda regularisation parameters, log-spaced between 1^−6^ and 1^+6^, and picked the model with the highest predictive performance averaged across sensors. MEG sensor activity at each ms was modelled using the preceding 200 ms of envelope and pitch estimates. Both the acoustic and MEG signals were processed to have zero mean across frequency bands and sensors respectively, and scaled to have unit variance before fitting the model. MEG acoustic-based predictions were then transformed back into original MEG units before regressing out of the true MEG signals. This process, including fitting hyper-parameters, was applied for each story recording and for each subject separately, across 3 folds. Because all analyses are fit on these residual data, we can interpret our results knowing that they cannot be accounted for by low-level acoustic attributes such as the acoustic amplitude.

The data were bandpass-filtered between 0.1 and 50 Hz using MNE-Python’s default parameters with firwin design [43] and downsampled from 1000 Hz to 250 Hz. Epochs were segmented from 200 ms pre-phoneme onset to 600 ms post-phoneme onset. No baseline correction was applied. Our results are not dependent upon the filter applied.

### 3.6. Preprocessing auditory signals

We computed a time-frequency decomposition of the auditory signals by deriving a mel spectrogram representation using the Python module *librosa* (version 0.8.0). We applied a 2048 sample Hamming window to the auditory waveform, with a 128 sample overlap between successive frames. We derived a power estimate at each of 208 frequency bands (analogous to the 208 MEG channels) using non-linearly spaced triangular filters from 1-11250 Hz. These data were then also downsampled to 250 Hz, and segmented from 200-600 ms in order to match the dimensionality and size of the MEG epochs. No baseline correction was applied.

We also tried using 50 linearly spaced frequency bands, and this did not change the interpretation of our acoustic analyses.

### 3.7. Modelled features

We investigated whether single-trial sensor-level responses varied as a function of fourteen binary phonetic features, as derived from the multi-value feature system reported in [44]. Note that this feature system is sparse relative to the full set of distinctive features that can be identified in English; however, it serves as a reasonable approximation of the phonemic inventory for our purposes.

#### Voicing

This refers to whether the vocal chords vibrate during production. For example, this is the difference between *b* versus *p* and *z* versus *s*.

#### Manner of articulation

Manner refers to the way by which air is allowed to pass through the articulators during production. Here we tested five manner features: fricative, nasal, plosive, approximant, and vowel.

#### Place of articulation

Place refers to where the articulators (teeth, tongue, lips) are positioned during production. For vowels, this consists of: central vowel, low vowel, mid vowel, high vowel. For consonants, this consists of: coronal, glottal, labial and velar.

#### Nuisance variables

In the same model, we also accounted for variance explained by ‘nuisance variables’ – i.e. structural and statistical co-variates of the phonemes. Though we were not interested in interpreting the results of these features, we included them in the model to be sure that they did not account for our main analysis on the phonetic features. These features included: primary stress, secondary stress, frequency of the phoneme sequence heard so far, suffix onset, prefix onset, root onset, syllable location in the word, and syllable onset. These features were extracted from the English Lexicon Project [45].

#### Subset variables

Throughout the analysis, we subset trials based on their relationship to: word onset, word offset, surprisal, entropy, distance from onset, distance from offset.

Surprisal is given as:

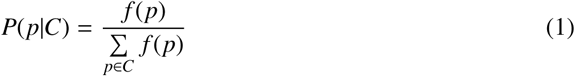

and cohort entropy is given as:

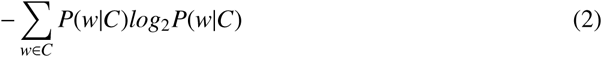

where *C* is the set of all words consistent with the heard sequence of phonemes thus far, and *f* (*w*) is the frequency of the word *w* and *f*(*p*) is the frequency of the phoneme *p*. Measures of spoken word frequency were extracted from the English Lexicon Project [45].

### 3.8. Decoding

Decoding analyses were performed separately on the acoustic signal and on the neural signal. For the acoustic decoding, the input features were the power estimates at each of the 208 frequency bands from 1-1125 Hz. For the neural decoding, the input features were the magnitude of activity at each of the 208 MEG sensors. This approach allows us to decode from multiple, potentially overlapping, neural representations, without relying on gross modulations in activation strength [46].

Because some of the features in our analysis are correlated with one another, we need to jointly evaluate the accuracy of each decoding model relative to its performance in predicting all modelled features, not just the target feature of interest. This is because, if evaluating each feature independently, we will not be able to dissociate the decoding of feature *f* from the decoding of the correlated feature 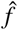. The necessity to use decoding over encoding models here, though (which, do not suffer so harshly from the problem of co-variance in the stimulus space) is one of signal-to-noise: we expect any signal related to linguistic processes to be contained in low-amplitude responses that are distributed over multiple sensors. Our chances of uncovering reliable responses to these features is boosted by using multi-variate models [46].

To overcome the issue of co-variance, while still capitalising on the advantages of decoding approaches, we implement a back-to-back ridge regression model [47]. This involves a two stage process. First, a ridge regression model was fit on a random (shuffled) half of the epoched data. The model was applied independently at each of 201 time-points in the epoch. We did not use a sliding window. The mapping was learnt between the multivariate input (either activity across sensors or power over frequency bands) and the univariate stimulus feature (one of the 31 features described above). All decoders were fit on data normalised by the mean and standard deviation in the training set:

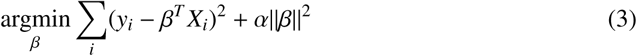

where *y*_*i*_ ∈ {±1} is the feature to be decoded at trial *i* and *X*_*i*_ is the multivariate acoustic or neural measure. The l2 regularisation parameter *α* was also fit, testing 20 log-spaced values from 1^−5^ to 1^5^. This was implemented using the *RidgeCV* function in *scikit-learn* [48].

Then, we use the other half of the acoustic or neural responses to generate a prediction for each of the 31 features corresponding to the test set. However, because the predictions are correlated, we need to jointly evaluate the accuracy of decoding each feature, to take into account the variance explained by correlated non-target features. To do this, we fit another ridge regression model, this time learning the beta coefficients that map the matrix of *true* feature values to vector of *predicted* values for the target feature of interest:

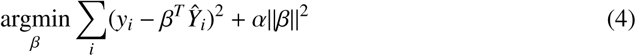

where *y*_*i*_ ∈ {±1} is the ground truth of a particular stimulus feature at trial *i* and *Ŷ*_*i*_ is the prediction for all stimulus features. A new regularisation parameter *α* was learnt for this stage. By including all stimulus features in the model, this accounts for the correlation between the feature of interest and the other features. From this, we use the beta-coefficients that map the true stimulus feature to the predicted stimulus feature. Beta coefficients serve as a metric of decoding performance: if a stimulus feature is not encoded in neural responses (the null hypothesis) then there will be no meaningful mapping between the true feature *y* and the model prediction *ŷ*. Thus, the beta coefficient will be zero – equivalent to chance performance. If, however, a feature *is* encoded in neural activity (the alternative hypothesis), we should uncover a significant relationship between *y* and *ŷ*, thus yielding an above-zero beta coefficient.

The train/test split was performed 100 times, and the beta-coefficients were averaged across iterations. This circumvents the issue of unstable coefficients when modelling correlated variables. These steps were applied to each subject independently.

### 3.9. Temporal generalisation decoding

Temporal generalization (TG) consists of testing whether a temporal decoder fit on a training set at time *t* can decode a testing set at time *t*′ [24]. This means that rather than evaluating decoding accuracy just at the time sample that the model was trained on, we evaluate its accuracy across all possible train/testing time combinations.

TG can be summarised with a square training time × testing time decoding matrix. To quantify the stability of neural representations, we measured the duration of above-chance generalisation of each temporal decoder. To quantify the dynamics of neural representations, we compared the mean duration of above-chance generalization across temporal decoders to the duration of above-chance temporal decoding (i.e. the diagonal of the matrix versus its rows). These two metrics were assessed within each subject and tested with second-level statistics across subjects.

### 3.10. Comparing decoding performance between trial subsets

To compare decoding performance for different subsets of trials that are of theoretical interest (e.g. between high/low surprisal, or beginning/end of word), we add a modification to our train/test cross-validation loop. The data are trained on the entire training set (i.e. the same number of trials as the ‘typical analysis’), and test set is grouped into the different levels of interest. We evaluate model performance separately on each split of the test data, which yields a time-course or generalisation matrix for each group of trials that we evaluate on.

### 3.11. Group statistics

In order to evaluate whether decoding performance is better than chance, we perform second-order permutation-based statistics. This involves testing whether the distribution of beta coefficients across subjects significantly differs from chance (zero) across time using a one-sample permutation cluster test with default parameters specified in the MNE-Python package [43]. The permutation test first computes the observed metric of interest, namely the summed t-value in a temporal cluster. The metric is then re-computed 10,000 times, each time randomly permuting the sign of the beta coefficients before finding temporal clusters. Significance is assessed by comparing the proportion of times that the summed t-value in any identified temporal cluster in the null distribution exceeds the observed sum t-value in the cluster of interest.

### 3.12. Spatial coefficient analysis

To test the spatial evolution of phonetic representations over time, we estimated beta coefficients by fitting linear regression (i.e. ridge regression with no regularisation, so that the coefficients were interpretable). We did this both in a decoding approach (predicting a single stimulus property from multivariate sensor activity) and in an encoding approach (predicting a single sensor response from multivariate stimulus properties). Because our goal was to estimate the model weights rather than to predict out of sample data, we fit the model on all data (i.e. no cross validation). We confirmed that the beta coefficients from the decoding model and encoding model yielded the same result and continued with the decoding coefficients for conceptual simplicity.

We averaged the coefficients across subjects and took the root mean square. Then, to compute the trajectories, we created two orthogonal sensor masks. First, a y-coordinate mask whereby the posterior sensors were coded as ‘0’ and anterior sensors were coded as ‘1’. Second, a x-coordinate mask, whereby the left sensors were coded as ‘0’ and the right sensors as ‘1’. These masks were normalised to have a norm equal to 1. The coefficients were projected onto these masks at each time sample, to yield a cosine distance over time, within the x-y coordinate plane.

We computed a metric of trajectory structure which was a weighted combination of range of movement (*m*, maximum cosine distance minus minimum), smoothness (*s*, mean absolute step size at each time sample) and variance (*v*, standard deviation across time samples), thus:

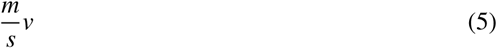

We compute this metric for the x co-ordinates and y co-ordinates separately, and then average the result. To evaluate whether a measure was statistically significant, we generated a surrogate ‘null’ dataset. For each subject, we ran the decoding analysis with shuffled feature labels. This provided an estimate of chance decoding, which we could use to generate null processing trajectories to compare to the trajectories in our empirical data. If the trajectory emerges from chance, movements should all be centred around zero (see Supplemental Figure 21). Whereas if the trajectory contains meaningful structure, it should significantly differ from the distribution of random trajectories (see Figure Supplemental 20). Significance was evaluated from the proportion of instances where the observed trajectory structure metric exceeded the null distribution.

### 3.13. Simulation of phoneme sequence reconstruction

To quantify how many phonemes the brain processes at once, and how it retains the order of those phonemes, we simulated responses to 4-phoneme anagrams. We estimated the spatiotemporal coefficients from the decoding model without regularisation (the same coefficients used for the trajectory analysis) for voicing and plosive manner of articulation. Simulated responses were generated by multiplying the coefficients of each feature by a noise factor, shifting the phoneme response by 100 ms for each phoneme position and summing the result. This resulted in twenty-four unique sequences without repetition (e.g., bpsz, pbzs, zspb …).

We simulated 10 responses to each unique phoneme sequence. We epoched the data around each phoneme onset and used ridge regression to reconstruct the phoneme sequence. We used cosine similarity to quantify the accuracy of the reconstruction. To evaluate statistical significance, we compared observed cosine similarity to the cosine similarity when randomly shuffling phoneme labels 10,000 times, and computed the proportion of times that the observed similarity exceeded the null distribution. We could accurately reconstruct the history of five previous phonemes significantly better than chance (1: cosine similarity 1.0, p < .001; 2: cosine similarity 0.91, p < .001; 3: cosine similarity 0.79, p < .001; 4: cosine similarity 0.38, p < .001; 5: cosine similarity 0.19, p = .012; 6: cosine similarity 0.1, p = .6).

### 3.14. Reconstructing time since phoneme onset

We assessed whether responses encoded elapsed time since phoneme onset by applying ridge regression to phoneme-locked responses. We cropped the time dimension from 100-400 ms which encapsulates the time window of strongest phonetic decoding, and downsampled to 53.3 Hz. For each subject, we collapsed the trial (∼50,000 phonemes) and time sample (16 time-points equally spaced between 100-400 ms) dimensions. This yielded around 800,000 neural responses per subject, each labelled for its relative time since phoneme onset.

Using a 5-fold cross validation loop with shuffled order, we fit ridge regression using default parameters in *scikit-learn*. For each trial, the input features to the model were the 208-channel sensor responses. The target features were latency since phoneme onset. Both were scaled to have zero mean and unit variance.

For each test set, we computed the model’s prediction of onset latency, which is a continuous measure between 100 and 400 (shown in Figure 4). In absence of evidence to the contrary, the model will predict the mean latency of the test set, in this case 250 ms. The within-fold correlation across subjects between the true and predicted latency since onset was highly significant (r = 0.83, p < .001).

## 4. Acknowledgements

We thank Graham Flick for help with data collection. We are very grateful to William Idsardi, Arianna Zuanazzi, Joan Orpella, Pablo Ripolles-Vidal and Omri Raccah for their feedback on an earlier version of the manuscript.

## Funding

This project received funding from the Abu Dhabi Institute G1001 (AM); European Union’s Horizon 2020 research and innovation program under grant agreement No 660086, the Bettencourt-Schueller Foundation, the Fondation Roger de Spoel-berch, the Philippe Foundation (JRK) and The William Orr Dingwall Dissertation Fellowship (LG).

## Author contributions

LG: conceptualisation; methodology; software; validation; formal analysis; investigation; data curation; writing - original draft preparation and review and editing; visualisation. JRK: conceptualisation; methodology; software; supervision. AM: conceptualisation; writing - review and editing; supervision; funding acquisition. DP: conceptualisation; writing - review and editing; supervision; funding acquisition.

## Competing interests

The authors declare no competing interests.

## Data and materials availability

Preprocessed data will be made available after publication.

## 5. Supplementary Results

### 5.1. Validation of back-to-back regression

**Figure 6:**
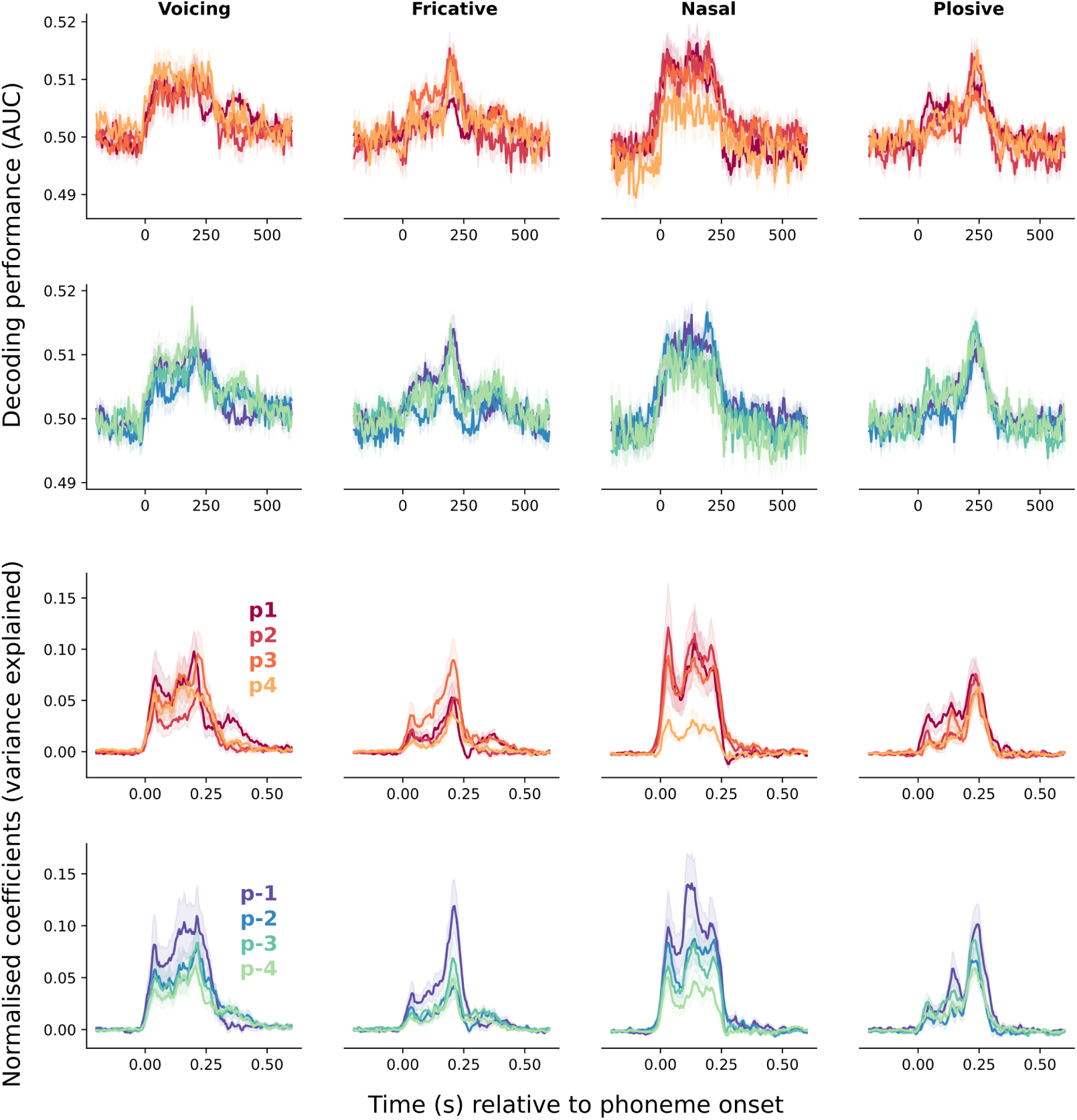
Comparing logistic regression to back-to-back regression. Above: Results of decoding four phonetic features using ‘classic’ logistic regression decoding analysis with AUC as the performance metric. Each line corresponds to a different phoneme location in the word. Below: Same analysis when using back-to-back regression.

### 5.2. Sequence analysis on mel spectrogram

**Figure 7:**
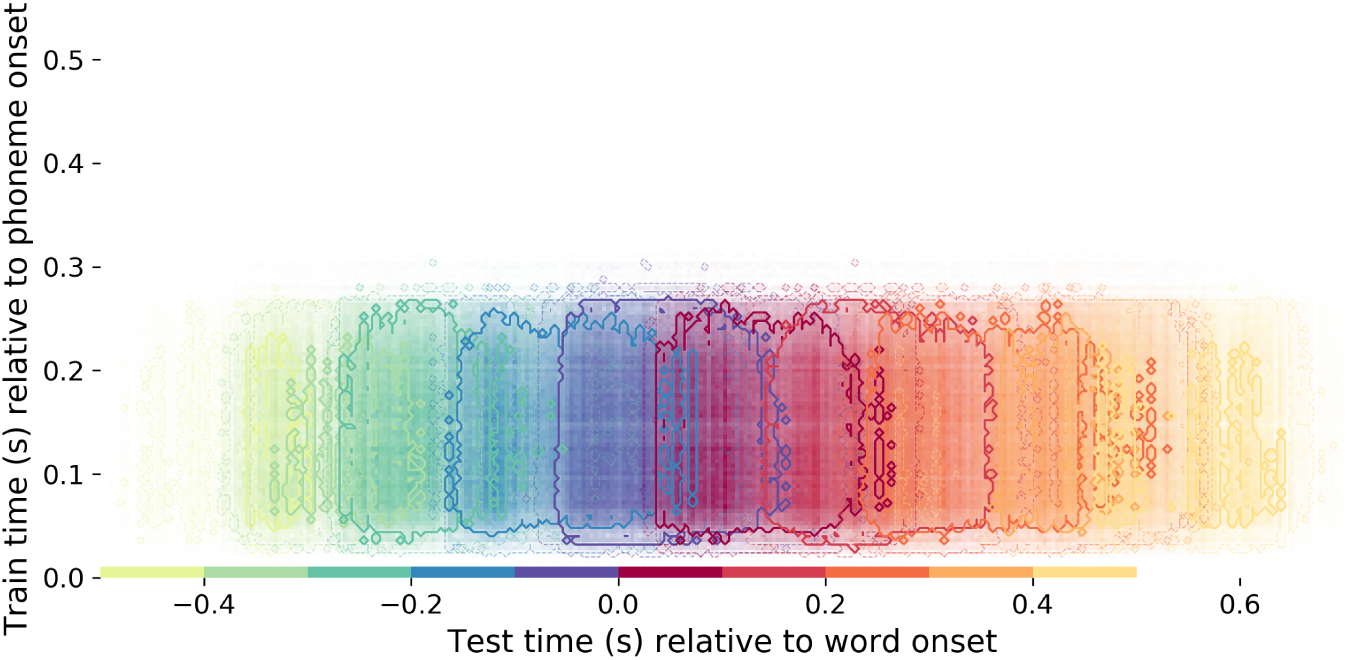
Temporal generalisation analysis of each phoneme position applied to the mel spectrogram.

**Figure 8:**
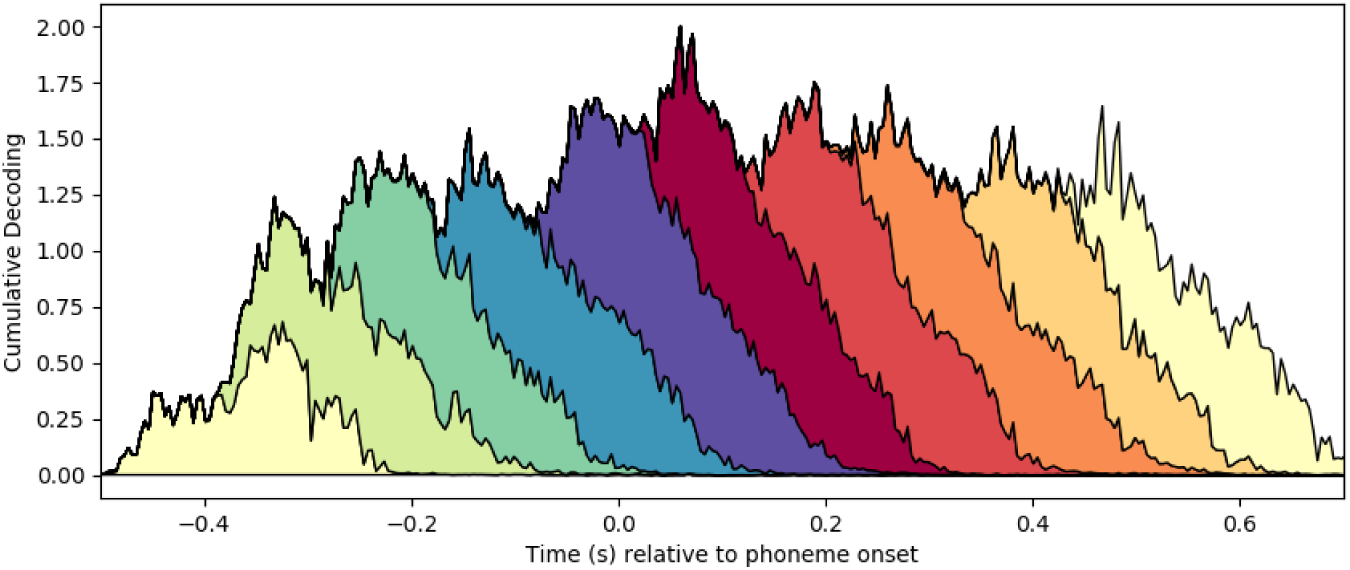
Cumulative decoding performance across the sequence, applied to mel spectrogram.

### 5.3. Signal to noise ratio manipulation on acoustic generalisation analysis

To test whether the sustained generalisation we observe in the auditory analysis is due to the relatively high SNR as compared to the MEG data, we re-ran the generalisation analysis when adding different levels of noise to the mel spectrogram. We observe a sustained decoding pattern at all SNR levels.

**Figure 9:**
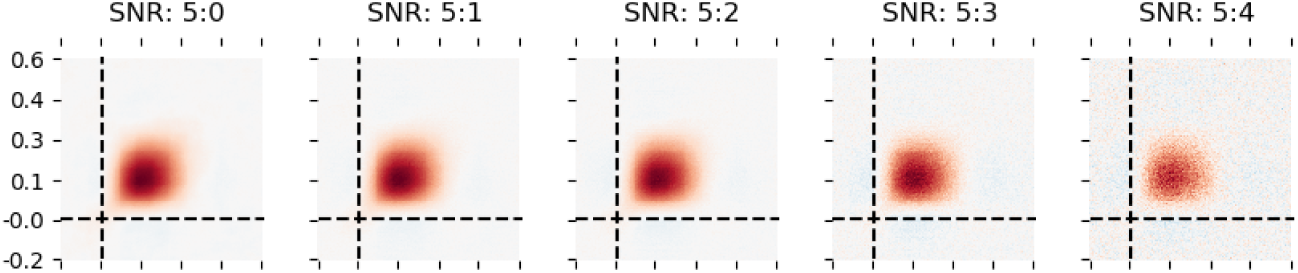
SNR manipulation on auditory decoding.

### 5.4. Strength of MEG signal and its relation to decoding performance

One potential confound in decoding performance is the overall signal strength (e.g. magnitude of the MEG response). It is possible that stimulus features which lead to larger responses will, in turn, aid better decodability of the other features encoded in that response. To test whether this was the case, we computed the root mean square (RMS) of the sensor data and attempted to decode this from the z-scored MEG responses, from 200 ms to 1000 after to each phoneme onset. For this, we used a Ridge regression decoder with Spearman R as the performance metric. First, we found that overall signal strength did not show the temporal dynamics that were elicited by any of our stimulus features of interest (see below figure). Second, we fit a mixed effects regression model between the single-trial RMS and decoding accuracy, for phonation, manner, and place of articulation. We modelled random slopes per subject and repetition. There was no significant relationship for any of the three features p’s ¿ .3. Overall this suggests that the features we show to interact with phonetic encoding strength (e.g. surprisal and entropy) cannot be explained by the global strength of the signal. We believe the lack of effect may be due to our stimuli: continuous speech does not elicit clear evoked responses, and so single-trial variability in signal strength is negligible.

**Figure 10:**
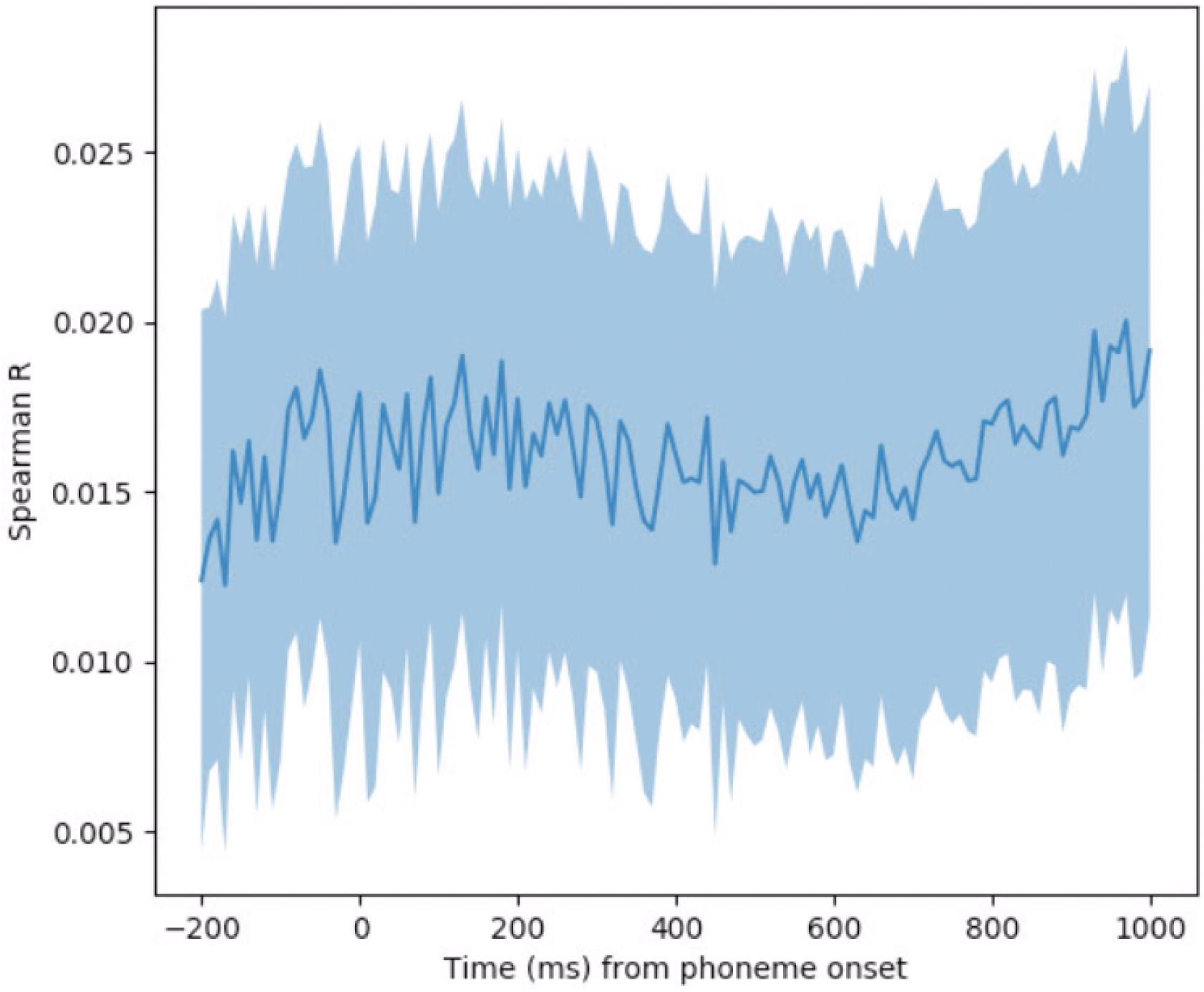
Decoding MEG signal strength.

### 5.5. History and future decoding

Our results suggest that at the brain processes at least three phonemes concurrently. To add further support to this claim, we decoded the content of three preceding phonemes and three subsequent phonemes from a given moment in test time. As shown in Figure 11, we can indeed decode the phonetic content of three phonemes from the same neural response. This was clearer for voicing and manner than place of articulation, in line with the general trend that place of articulation is not as robustly encoded as the other properties [9, 11]. For the history decoding, decoding performance appears to peak time-locked to the onset of subsequent phonemes, perhaps suggestive of a re-activation procedure. This is in line with previous results from our lab (e.g. [16]).

**Figure 11:**
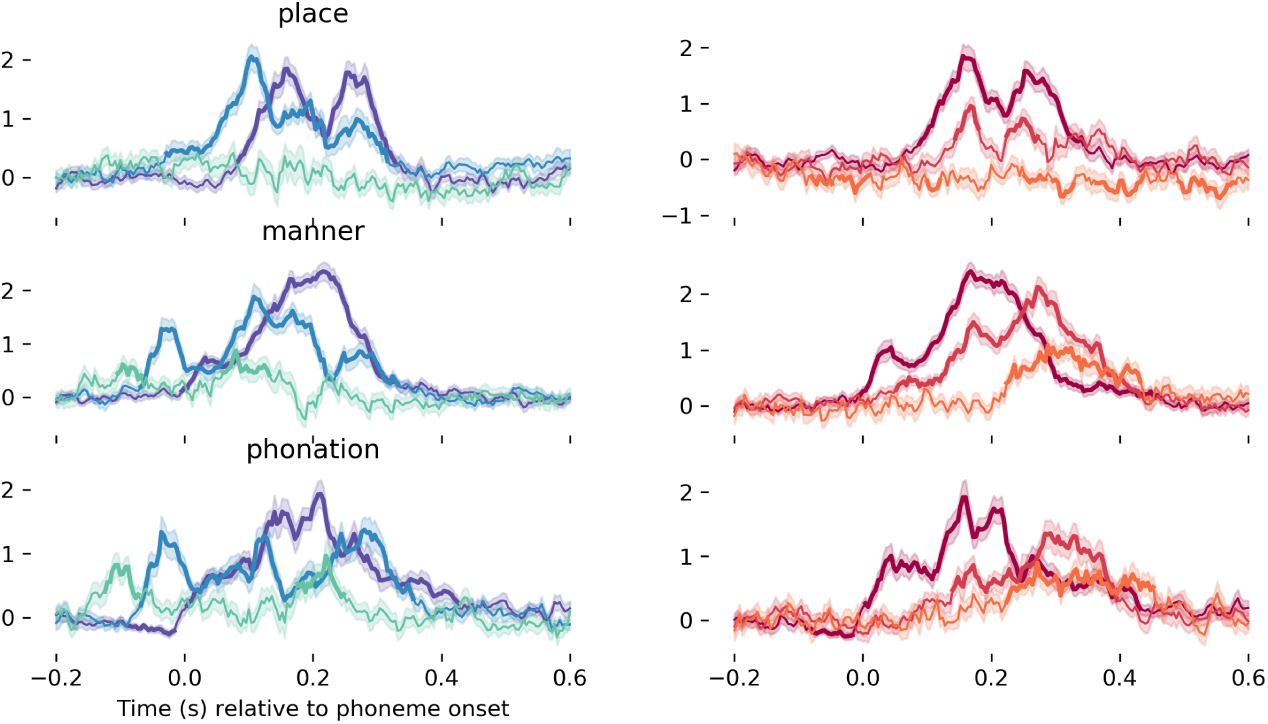
Decoding phonetic history and future from a single neural timecourse.

### 5.6. Analysis on equalised trial counts

Due to the natural variation in word length, there were different numbers of trials included in the decoding analysis at phoneme positions. Maximum numbers of trials at the word boundaries (because all words have an onset and an offset) and gradually decreasing numbers of trials as the position moves further from the boundary. Estimates of decodability are noisier for phoneme positions with fewer trials. To allow direct comparisons of decoding strength across positions, we re-ran the analysis matching the number of trials to the fourth phoneme position (about 1500 trials).

Once the number of trials was equalised across positions, there were no significant differences in the strength of phoneme decoding. We replicate all the main results on this subset of trials.

**Figure 12:**
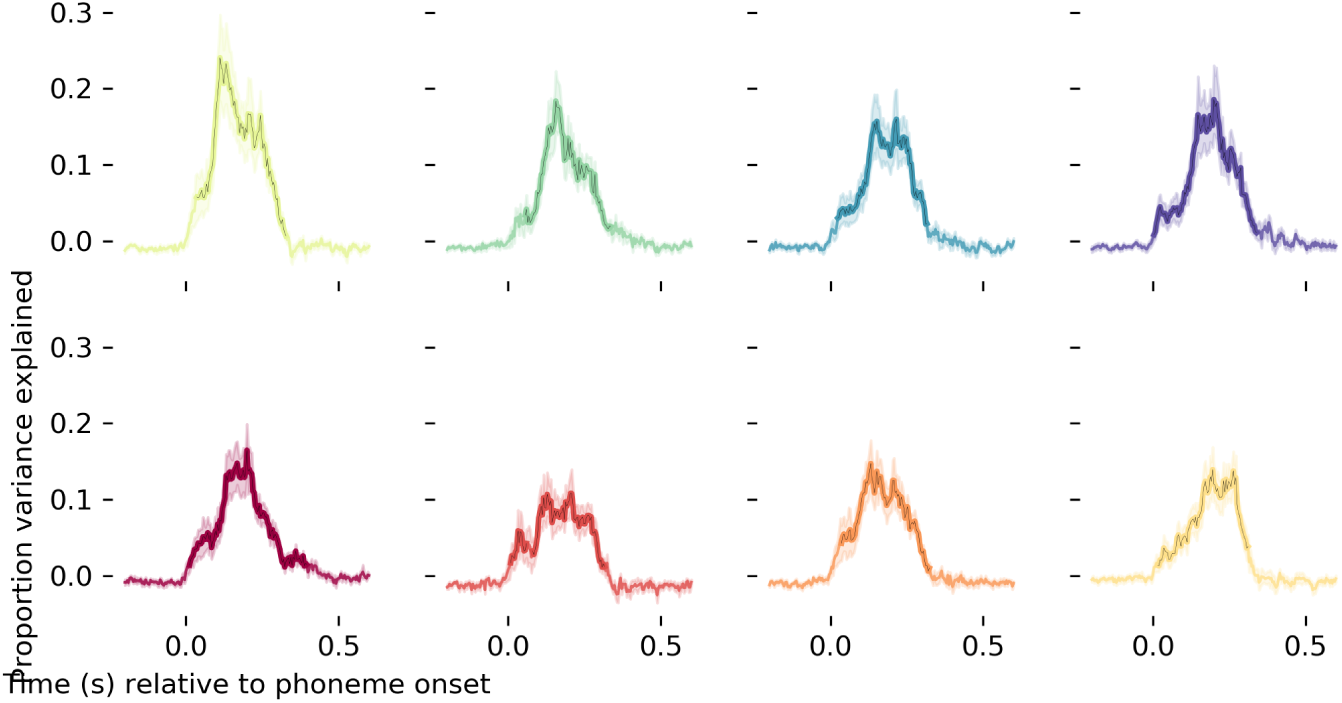
Diagonal decoding on equalised phoneme counts.

**Figure 13:**
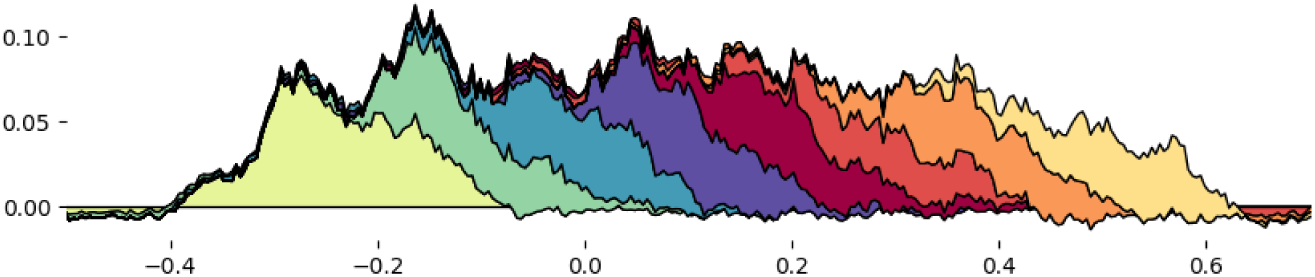
Cumulative diagonal decoding.

**Figure 14:**
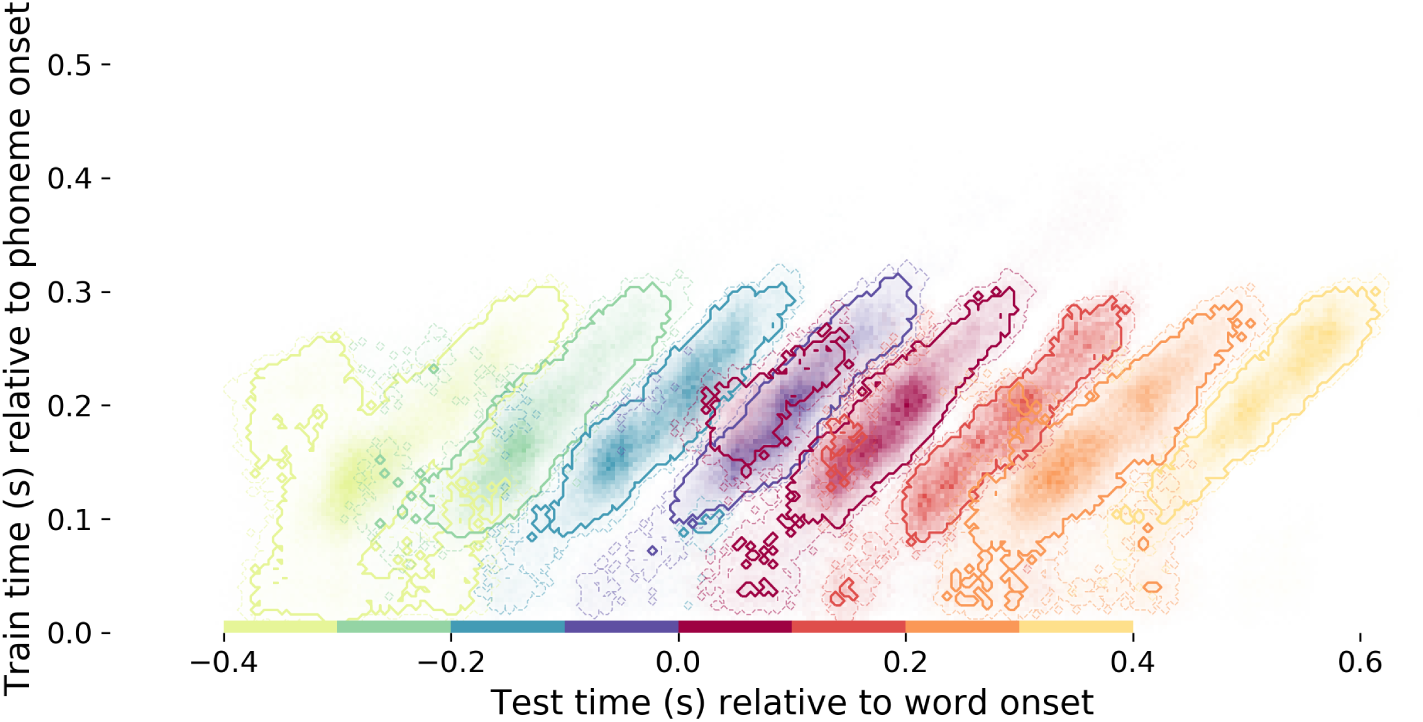
Temporal generalisation decoding on equalised phoneme counts.

### 5.7. Sequence representation for different phonetic features

A lot of the analyses we applied were aggregated over the individual phonetic features. However, even though the individual feature sequence maps are noisier, they show the same diagonal patterns as the full aggregated data.

There are, however, some interesting dynamics that may be worth exploring in future work. For example, the voicing feature is appears more sustained across positions. And the ‘appendage’ of the first phoneme appears most pronounced for manner of articulation.

**Figure 15:**
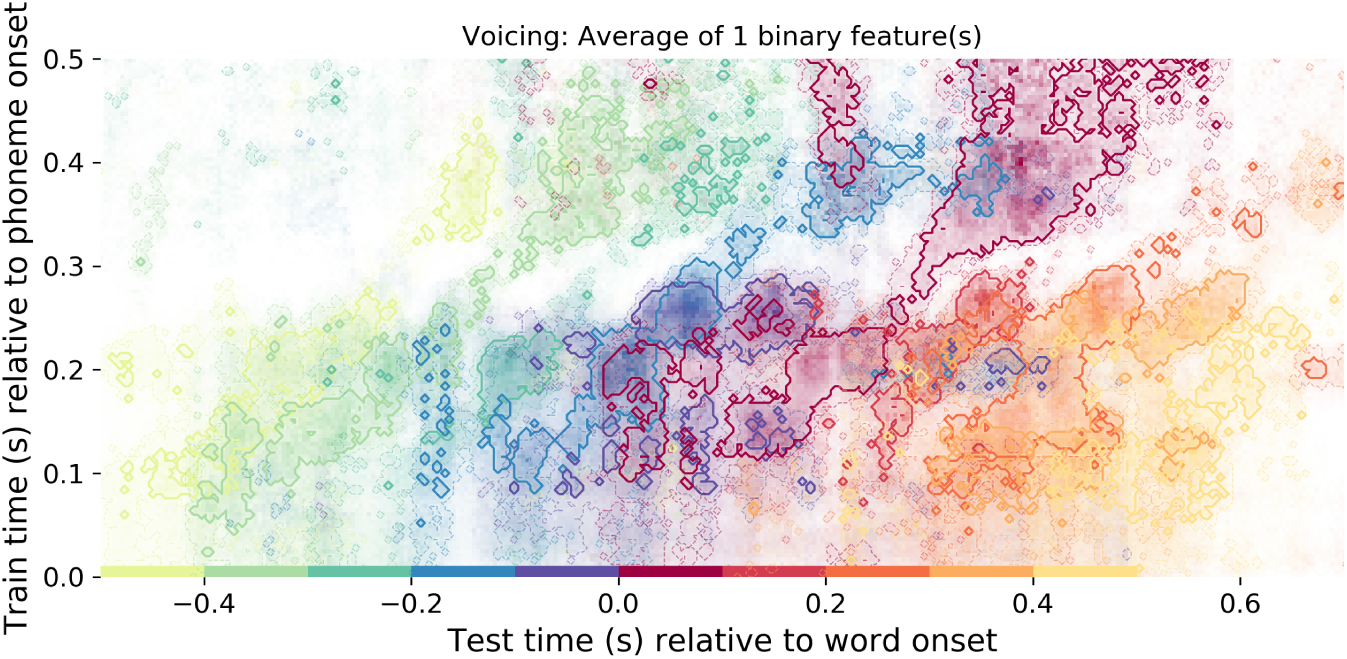
Sequence decoding for voicing.

**Figure 16:**
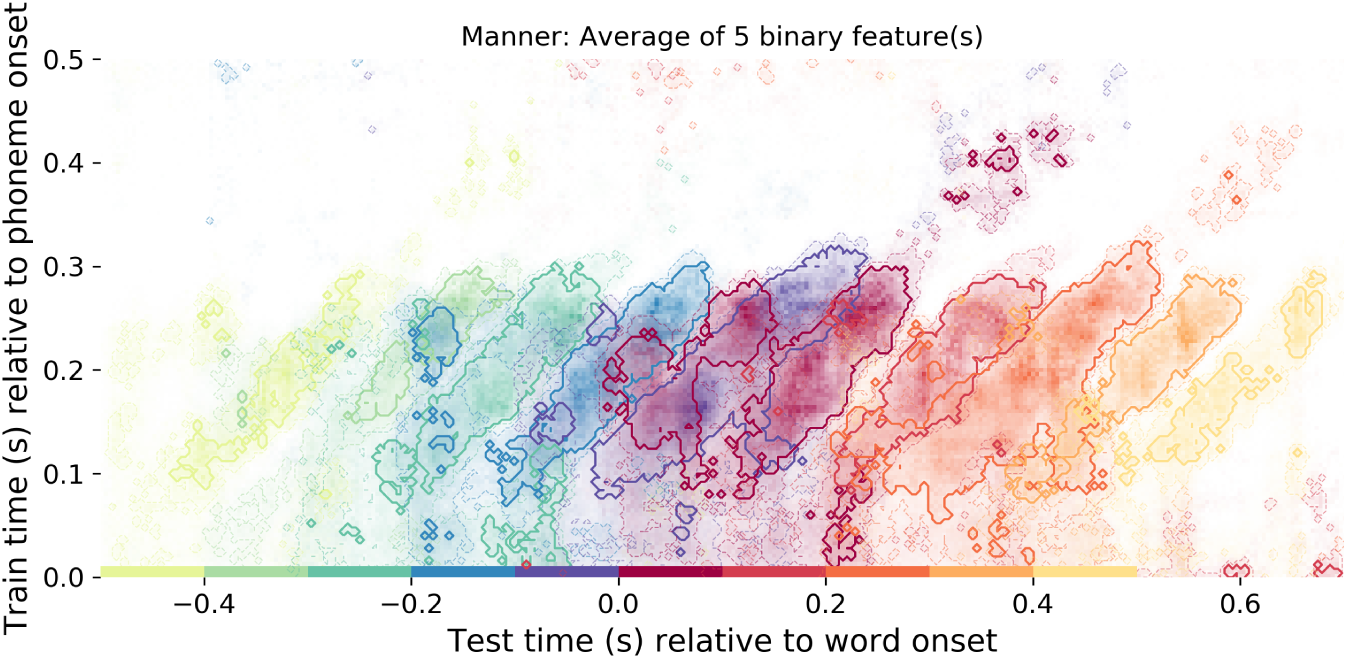
Sequence decoding for manner.

**Figure 17:**
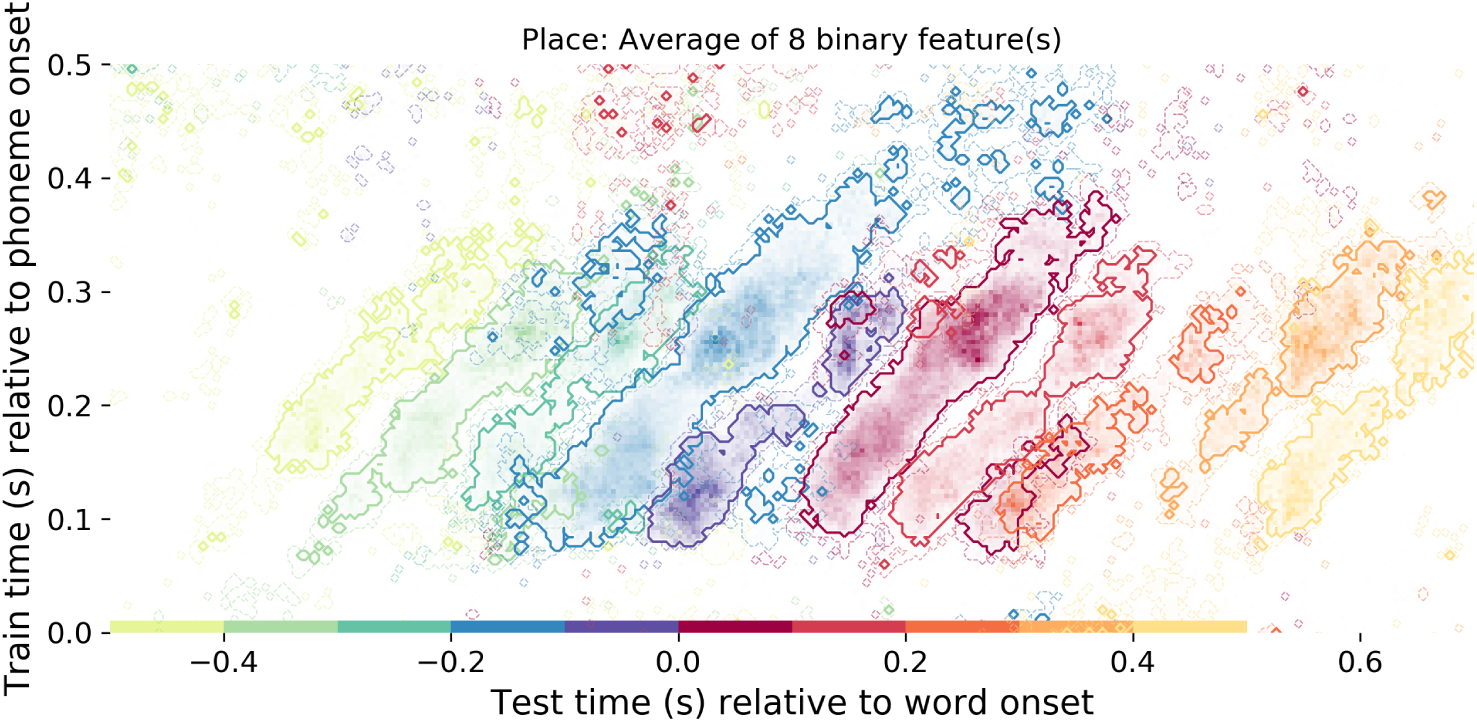
Sequence decoding for place of articulation.

### 5.8. Dynamics at sentence onset and sentence offset

We re-ran the neural generalisation analysis on two subsets of phonemes. First, testing on all phonemes at the onset of words and the onset of sentences (and training on all others). Second, testing on all phonemes at the offset of words and at the offset of sentences (and training on all others). The goal was to test whether (i) the ‘appendage’ observed for the onset phoneme was caused by predictability of the phoneme in context, in which case we should not observe it at the beginning of sentences because there is no context upon which to generate reliable predictions; (ii) whether we still observe ‘diagonal dynamics’ when the phoneme is followed by silence. The model was fit on responses to about 20,000 phonemes and tested on about 500 phonemes. The results show average decoding performance across phonetic features.

Interesting we observe that the ‘appendage’ is still present for onset phonemes at the beginning of sentences. This suggests that the maintenance of phonetic features may be some that always happens at the beginning of words, regardless of how predictable the word is in context. Future work should test the reason for this.

Furthermore we find that the diagonal pattern is also present for phonemes at the end of sentences which are followed by silence. It seems that phonetic detail is actually maintained during the silent period, another fascinating result which should be followed up with subsequent research.

**Figure 18:**
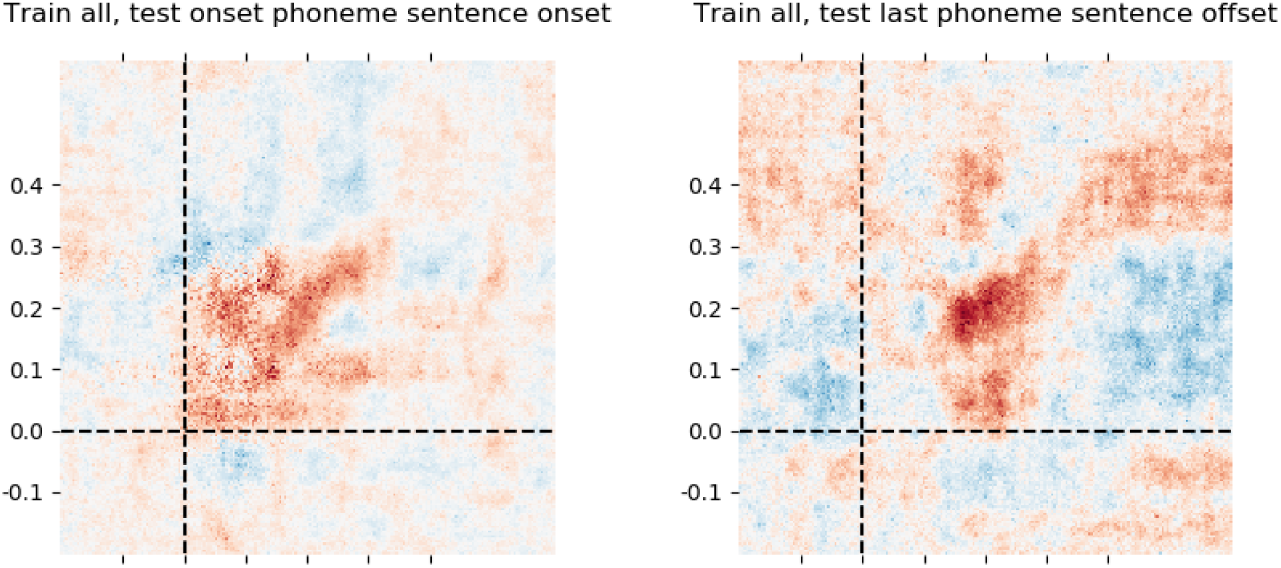
Generalisation analysis applied to the first phoneme of a sentence and the last phoneme of a sentence.

### 5.9. Testing granularity of representation

Throughout the manuscript we focus on phonetic features as the way to represent linguistically relevant information about speech sounds. One possibility is that, during the timecourse of processing, representations evolve from more sensory to more abstract - e.g. from acoustics to phonetic features to phoneme categories.

We ran the analysis on the raw (not acoustically residual data), trying to decode (i) spectral features of the phonemes (ii) the same 14 phonetic features that this manuscript is concerned with (iii) one-hot encoding of consonants (e.g. /b/, /p/) (iv) one-hot encoding of vowels (e.g. /ae/, /oo/) and (vi) all the other higher order features that we have analysed in the study. We tested these different formats of representing speech sounds for their relative timecourse of decodability. Unfortunately, the representations are too correlated to make any strong claims in one direction or another, and there was no clear temporal separation between the different formats. Future controlled studies may be able to associate different moments in the processing trajectory with different representational formats, once they have been sufficiently de-correlated in the design

**Figure 19:**
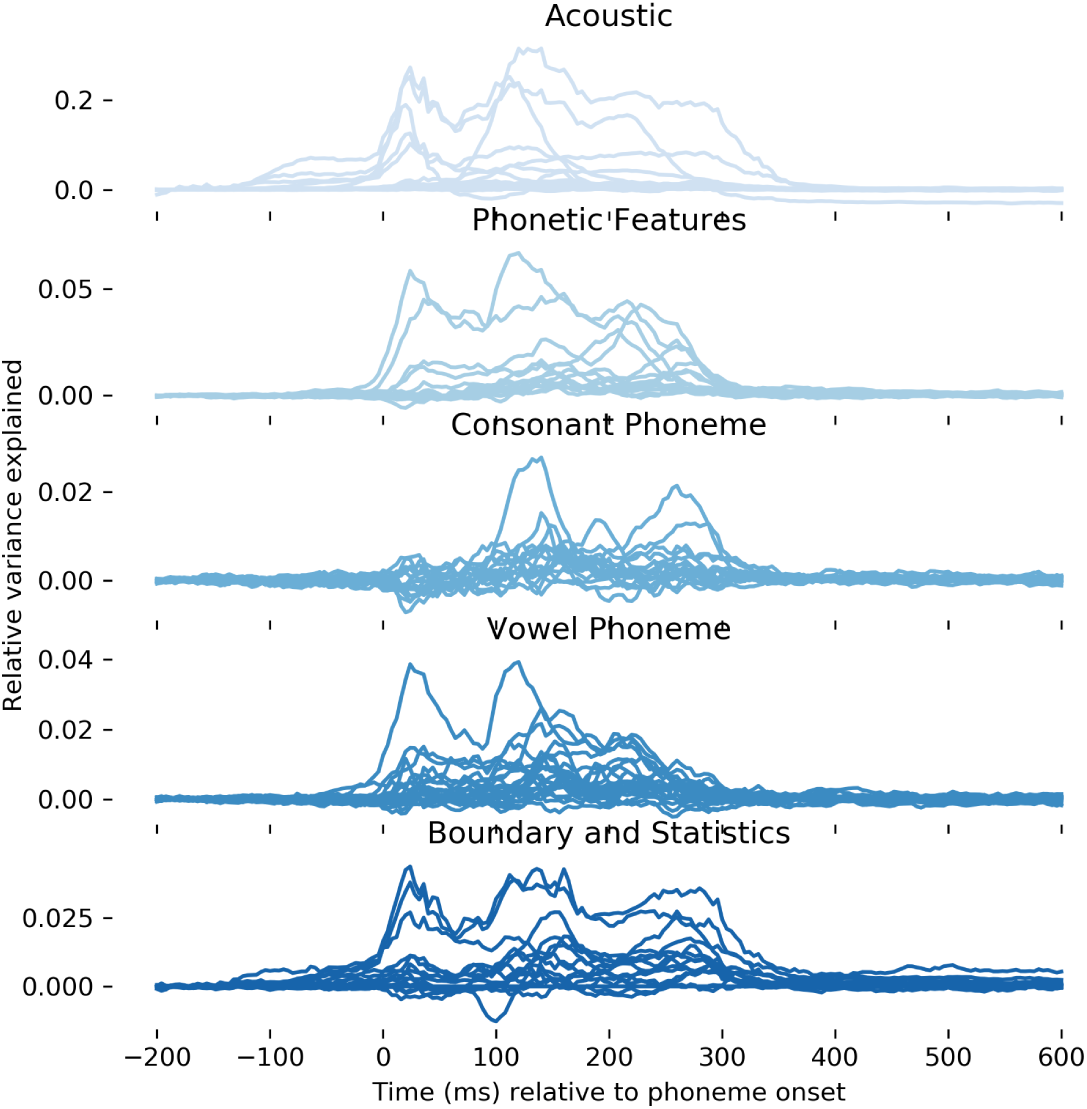
Decoding different speech sound representational formats.

### 5.10. Beta coefficients of decoding model

Here we show the coefficients of a linear decoding model averaged across subjects, for each of the 31 features. Each subject and each repetition of the story contains 25259 individual phoneme responses.

To evaluate statistical significance we also ran the decoding analysis on shuffled versions of each feature, for each subject and repetition. We computed a metric of trajectory structure which was a weighted combination of range of movement (*m*, maximum cosine distance minus minimum), smoothness (*s*, mean absolute step size at each time sample) and variance (*v*, standard deviation across time samples), thus:

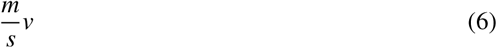

We compute this metric for the x co-ordinates and y co-ordinates separately, and then average the result. To test statistical reliability we ran an independent t-test between the coefficients from the true features and the coefficients from the shuffled features.

Some notable results are the robust position-based trajectories such as phoneme location in the word (True Mean = 1.72, SD = 0.85, Null Mean = 1.06, SD = 0.5, *t* = 4.25, *p* < .001) and phonetic feature trajectories such as voicing (True Mean = 1.42, SD = 0.59, Null Mean = 1.04, SD = 0.44, *t* = 3.24, *p* < .001), nasality (True Mean = 1.34, SD = 0.57, Null Mean = 1.07, SD = 0.4, *t* = 2.56, *p*= 0.01) and fricative (True Mean = 1.32, SD = 0.55, Null Mean = 1.09, SD = 0.38, *t* = 2.26, *p* = 0.03).

Our results for all of the 31 features are as follows: Word length: True Mean = 1.09, SD = 0.63, Null Mean = 1.19, SD = 0.5, *t* = −0.78, *p* = 0.44. Root Onset: True Mean = 1.7, SD = 0.62, Null Mean = 1.18, SD = 0.49, *t* = 4.27, *p* = 0.0. Phon. offset distance: True Mean = 1.28, SD = 0.73, Null Mean = 1.04, SD = 0.44, *t* = 1.86, *p* = 0.07. Entropy: True Mean = 1.58, SD = 0.62, Null Mean = 1.05, SD = 0.45, *t* = 4.43, *p* = 0.0. Phon. Loc Sentence: True Mean = 0.86, SD = 0.97, Null Mean = 1.11, SD = 0.42, *t* = −1.51, *p* = 0.13. Phon. Loc Syllable: True Mean = 1.39, SD = 0.56, Null Mean = 1.03, SD = 0.36, *t* = 3.44, *p* = 0.0. Phon. Loc Word: True Mean = 1.72, SD = 0.85, Null Mean = 1.06, SD = 0.5, *t* = 4.25, *p* = 0.0. Prefix Onset: True Mean = 1.21, SD = 0.54, Null Mean = 1.18, SD = 0.47, *t* = 0.34, *p* = 0.74. Primary Stress: True Mean = 1.52, SD = 0.49, Null Mean = 1.13, SD = 0.5, *t* = 3.48, *p* = 0.0. Secondary Stress: True Mean = 1.1, SD = 0.5, Null Mean = 0.95, SD = 0.37, *t* = 1.56, *p* = 0.12. Sequence Frequency: True Mean = 1.79, SD = 0.83, Null Mean = 1.09, SD = 0.39, *t* = 4.85, *p* = 0.0. Suffix Onset: True Mean = 1.38, SD = 0.51, Null Mean = 0.98, SD = 0.42, *t* = 3.85, *p* = 0.0. Surprisal: True Mean = 1.71, SD = 0.71, Null Mean = 0.97, SD = 0.32, *t* = 6.06, *p* = 0.0. Syll. Loc Word: True Mean = 1.77, SD = 0.92, Null Mean = 1.09, SD = 0.47, *t* = 4.23, *p* = 0.0. Syllable Onset: True Mean = 1.52, SD = 0.43, Null Mean = 1.1, SD = 0.34, *t* = 4.86, *p* = 0.0. Word Offset: True Mean = 1.44, SD = 0.62, Null Mean = 1.24, SD = 0.59, *t* = 1.53, *p* = 0.13. Word Onset: True Mean = 1.68, SD = 0.57, Null Mean = 1.12, SD = 0.49, *t* = 4.83, *p* = 0.0. Approximant: True Mean = 1.27, SD = 0.49, Null Mean = 1.14, SD = 0.38, *t* = 1.36, *p* = 0.18. Fricative: True Mean = 1.32, SD = 0.55, Null Mean = 1.09, SD = 0.38, *t* = 2.26, *p* = 0.03. Nasal: True Mean = 1.34, SD = 0.57, Null Mean = 1.07, SD = 0.4, *t* = 2.56, *p* = 0.01. Plosive: True Mean = 1.33, SD = 0.39, Null Mean = 1.16, SD = 0.44, *t* = 1.87, *p* = 0.06. Vowel: True Mean = 1.4, SD = 0.36, Null Mean = 1.11, SD = 0.41, *t* = 3.38, *p* = 0.0. Voicing: True Mean = 1.42, SD = 0.59, Null Mean = 1.04, SD = 0.44, *t* = 3.24, *p* = 0.0. Glottal: True Mean = 1.06, SD = 0.41, Null Mean = 1.12, SD = 0.5, *t* = −0.61, *p* = 0.54. Coronal: True Mean = 1.36, SD = 0.43, Null Mean = 1.15, SD = 0.48, *t* = 2.13, *p* = 0.04. Dental: True Mean = 1.25, SD = 0.49, Null Mean = 1.12, SD = 0.42, *t* = 1.34, *p* = 0.18. High Vowel: True Mean = 1.31, SD = 0.52, Null Mean = 1.06, SD = 0.41, *t* = 2.42, *p* = 0.02. Labial: True Mean = 1.25, SD = 0.51, Null Mean = 1.14, SD = 0.49, *t* = 1.01, *p* = 0.31. Low Vowel: True Mean = 1.18, SD = 0.53, Null Mean = 1.06, SD = 0.41, *t* = 1.18, *p* = 0.24. Middle Vowel: True Mean = 1.19, SD = 0.42, Null Mean = 1.16, SD = 0.54, *t* = 0.32, *p* = 0.75. Velar: True Mean = 1.13, SD = 0.41, Null Mean = 1.09, SD = 0.5, *t* = 0.45, *p* = 0.65.

#### 5.10.1. Trajectory for each feature

**Figure 20:**
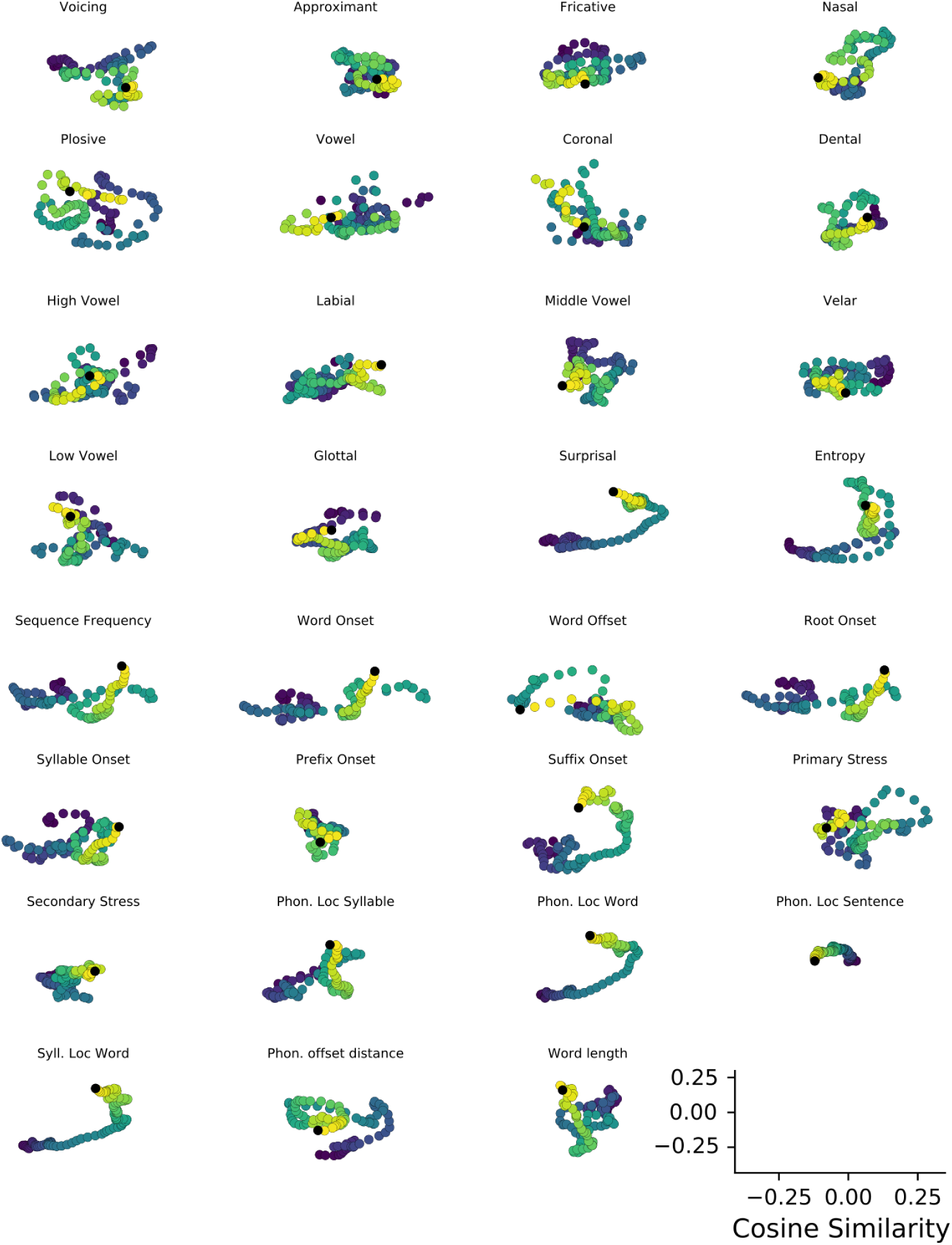
Trajectory for each feature.

**Figure 21:**
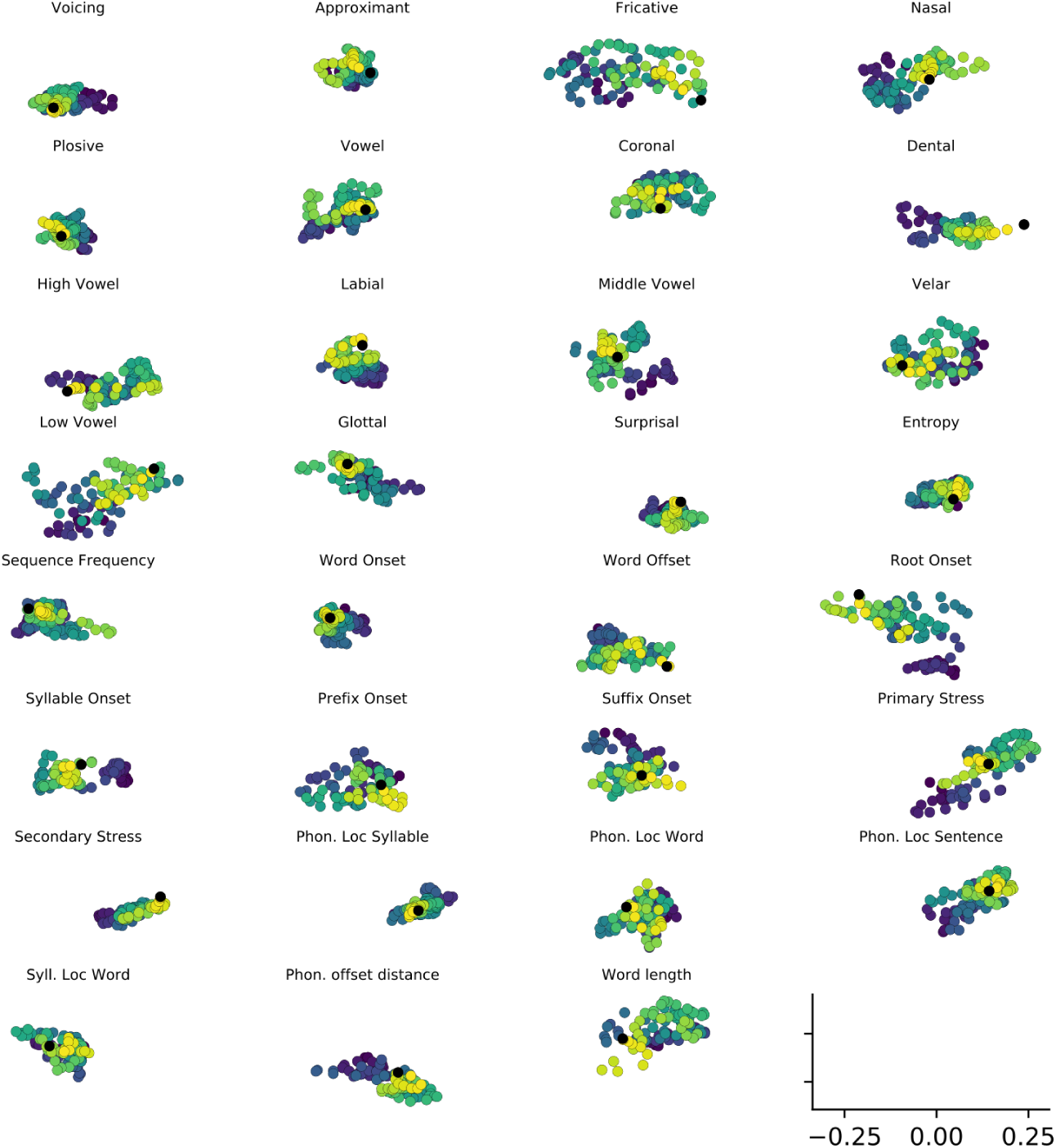
Trajectory when shuffling trial correspondence in decoding (i.e. null trajectory for comparison).

#### 5.10.2. Voicing

**Figure 22:**
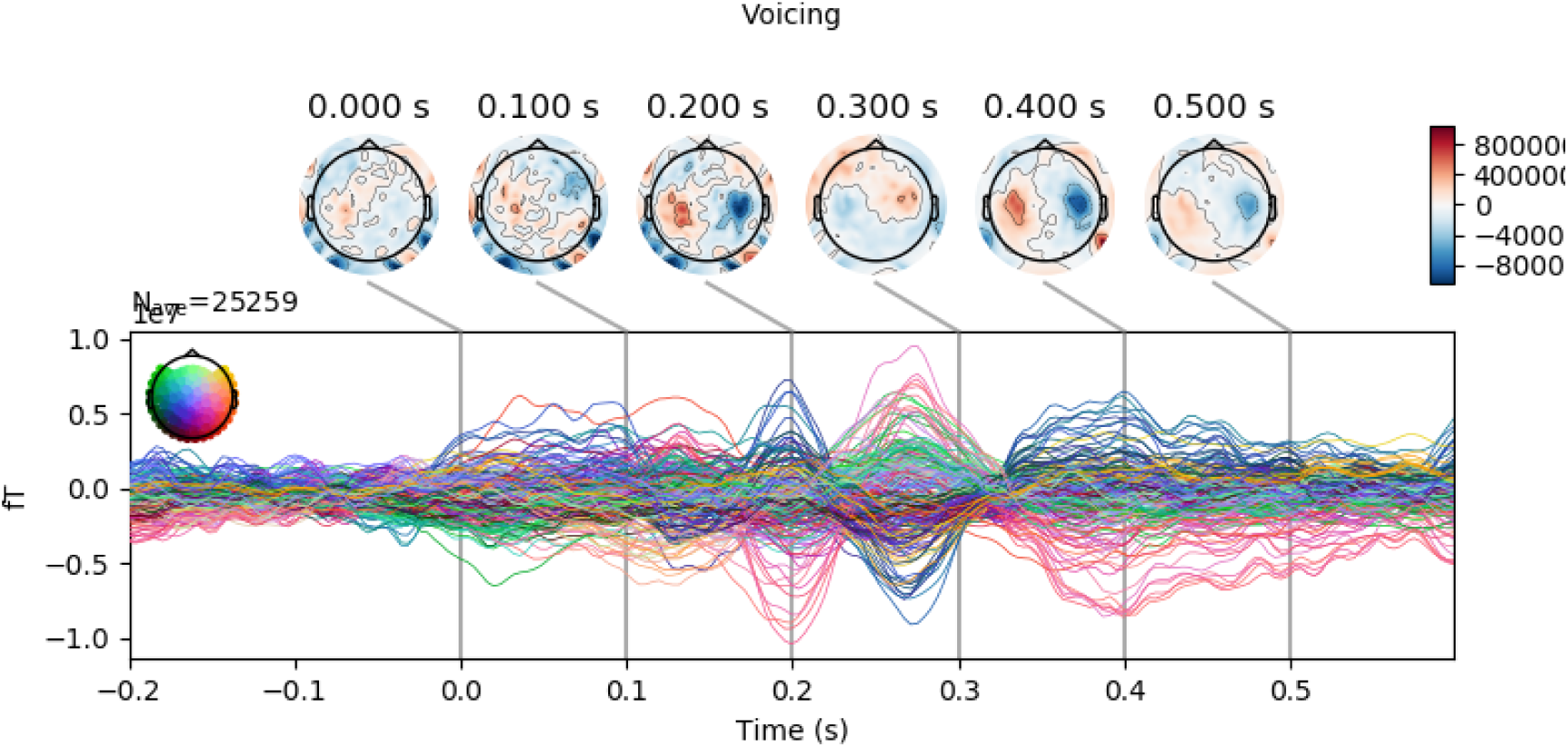
Coefficients for Voicing.

#### 5.10.3. Manner

**Figure 23:**
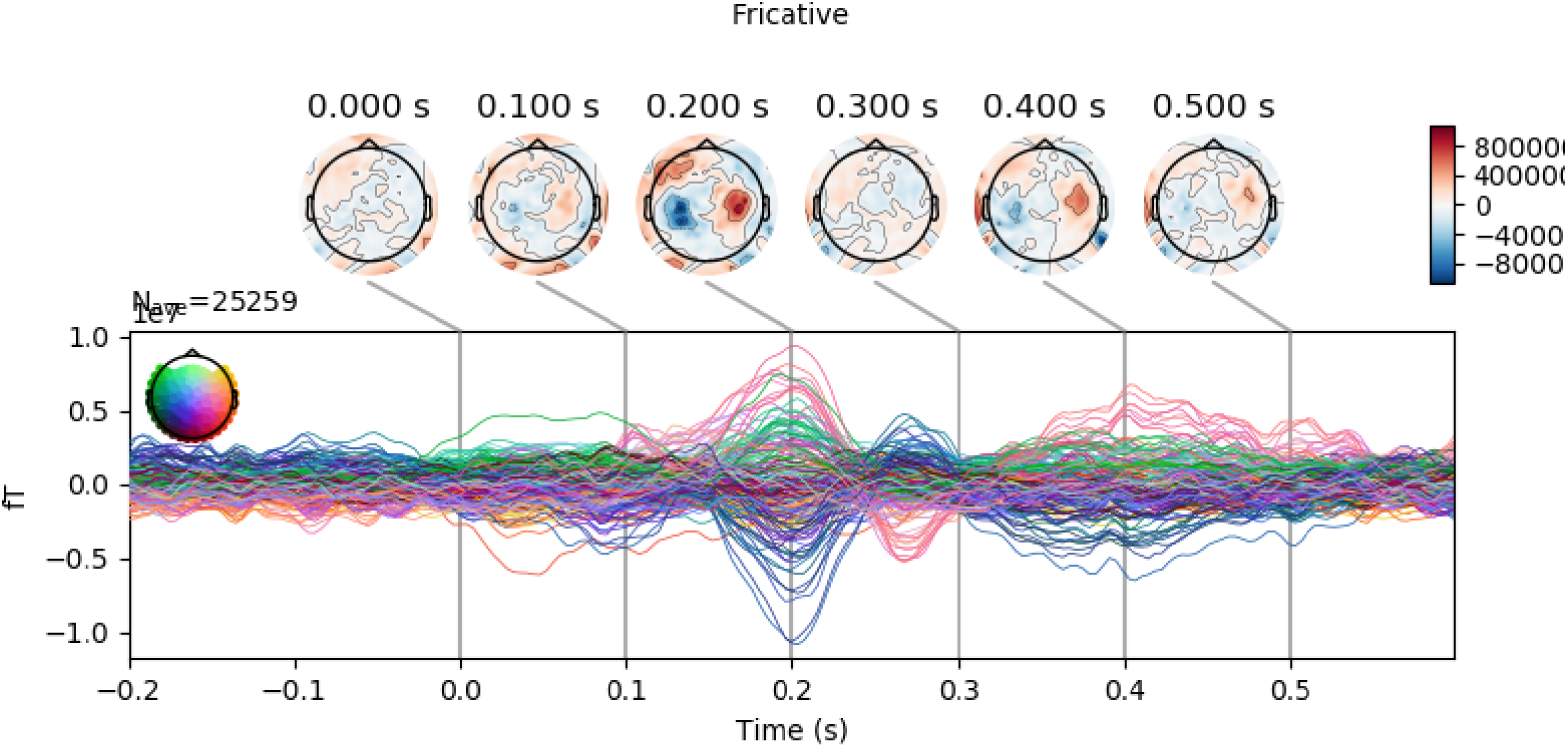
Coefficients for Frication.

**Figure 24:**
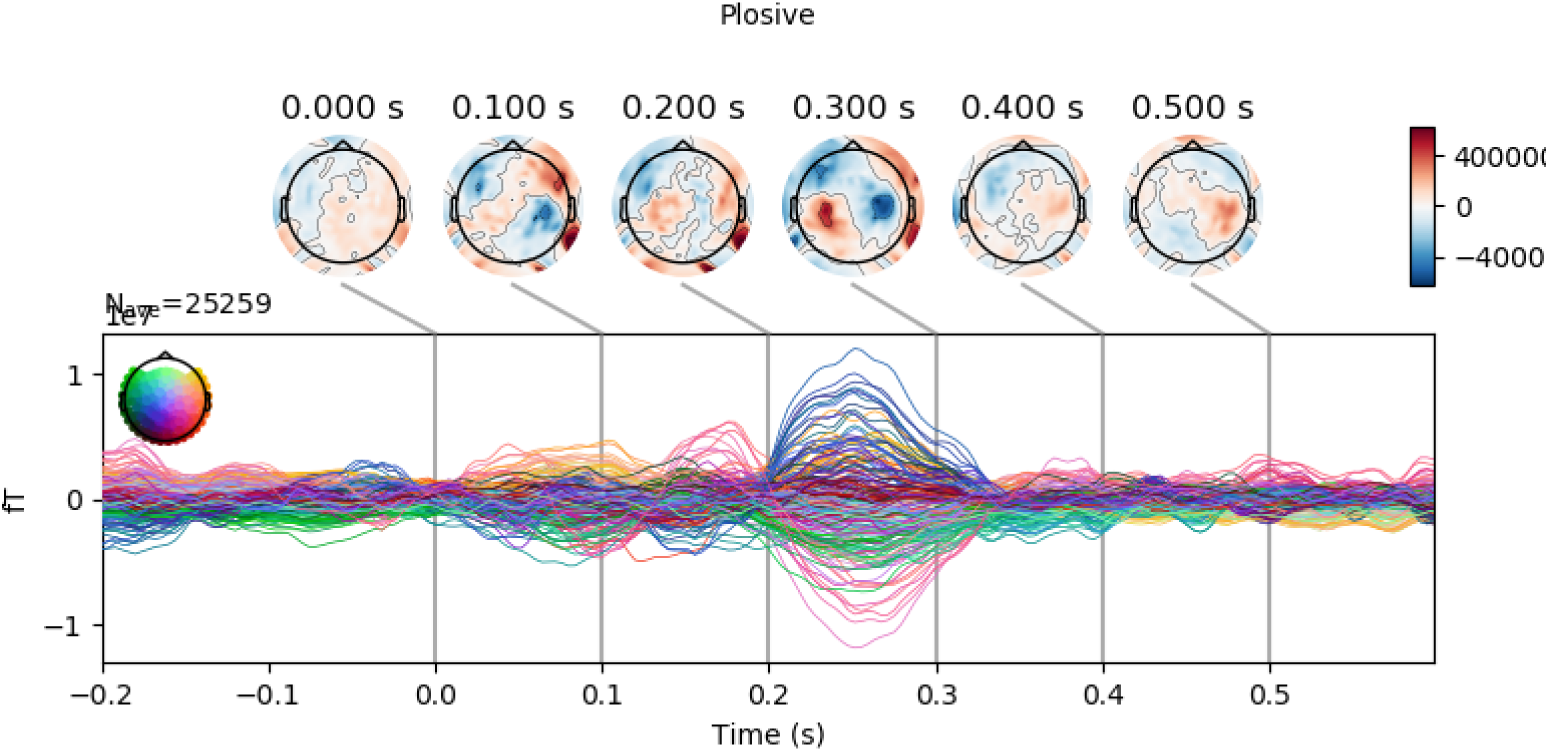
Coefficients for Plosive.

**Figure 25:**
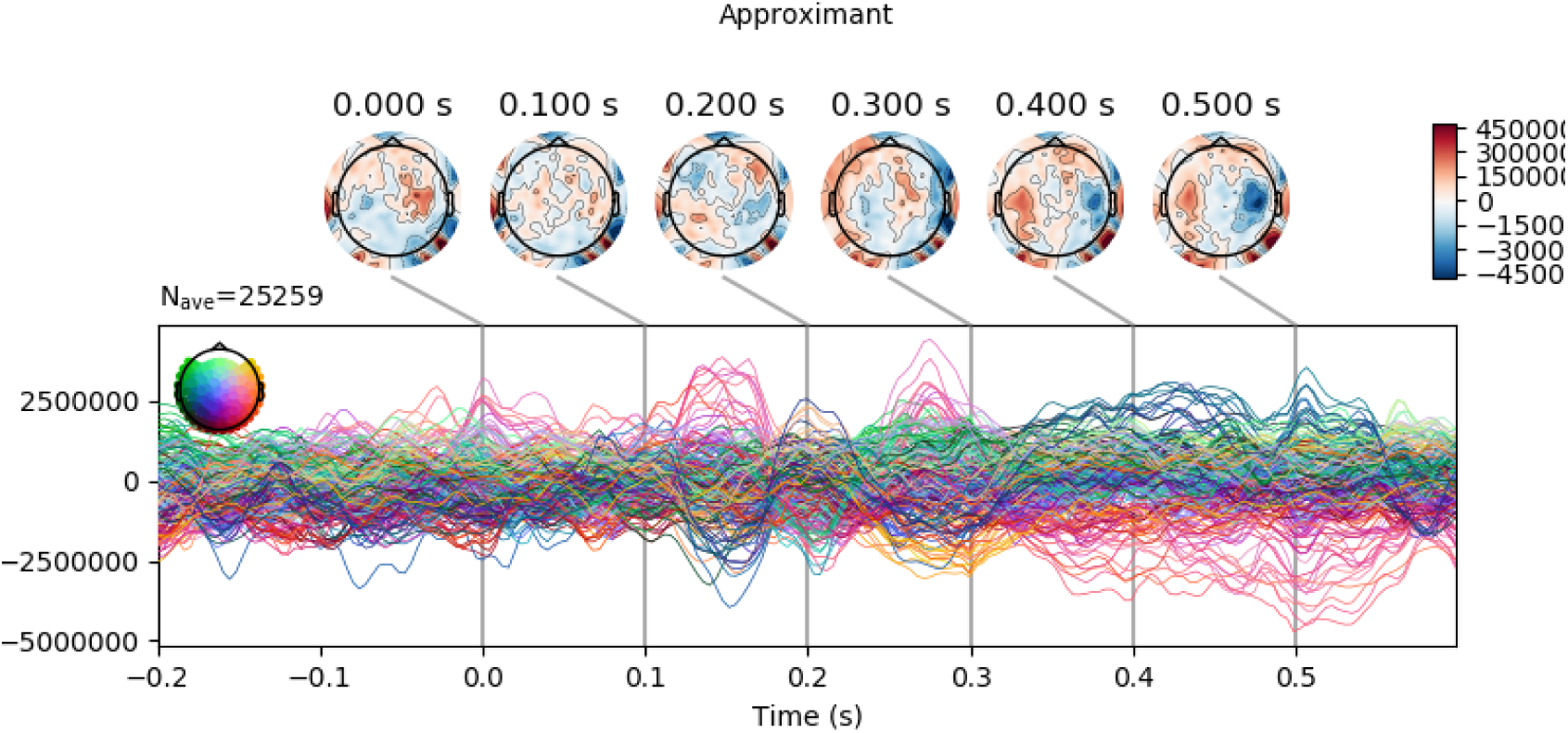
Coefficients for Approximant.

**Figure 26:**
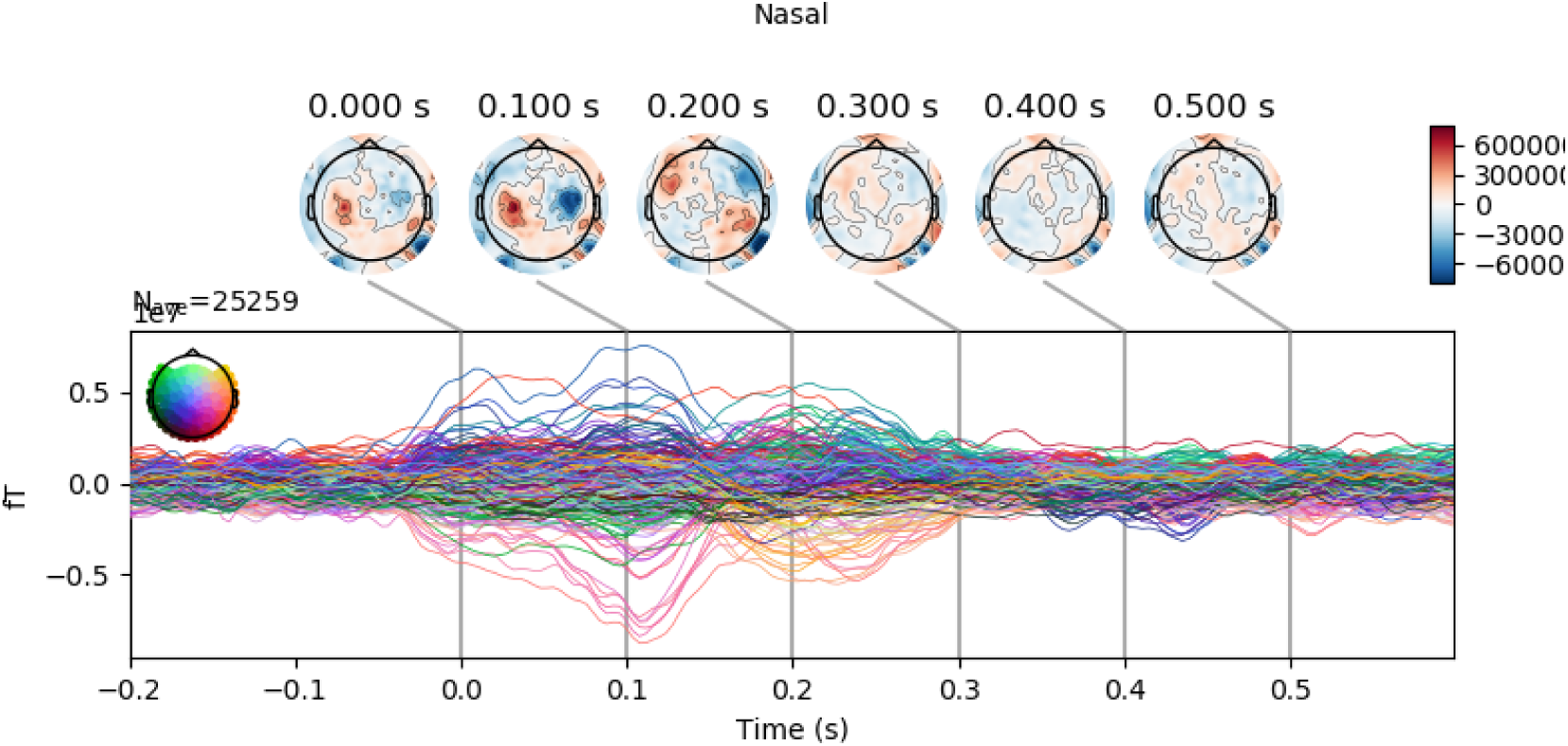
Coefficients for Nasal.

**Figure 27:**
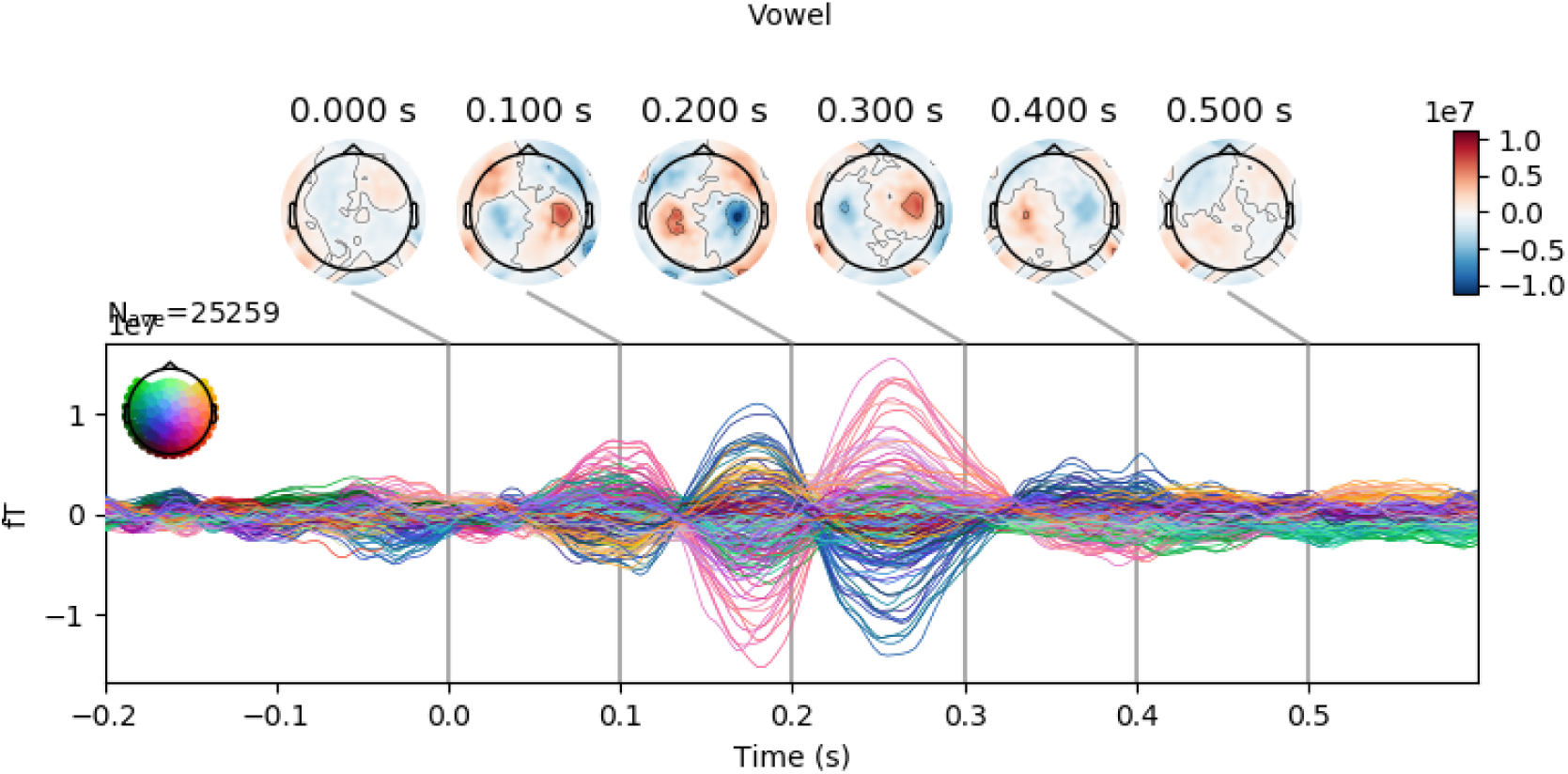
Coefficients for Vowel.

#### 5.10.4. Place of articulation: Consonants

**Figure 28:**
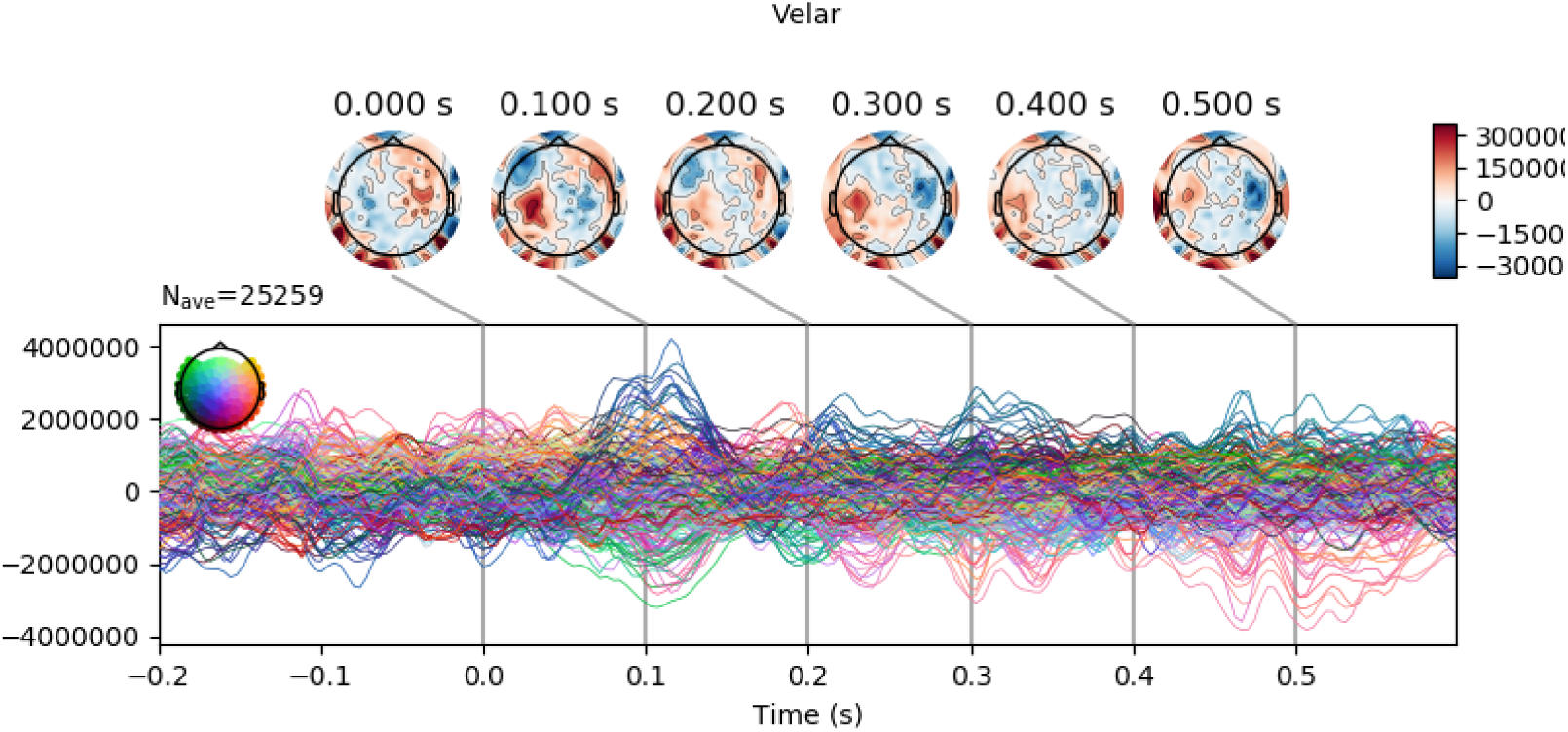
Coefficients for Velar.

**Figure 29:**
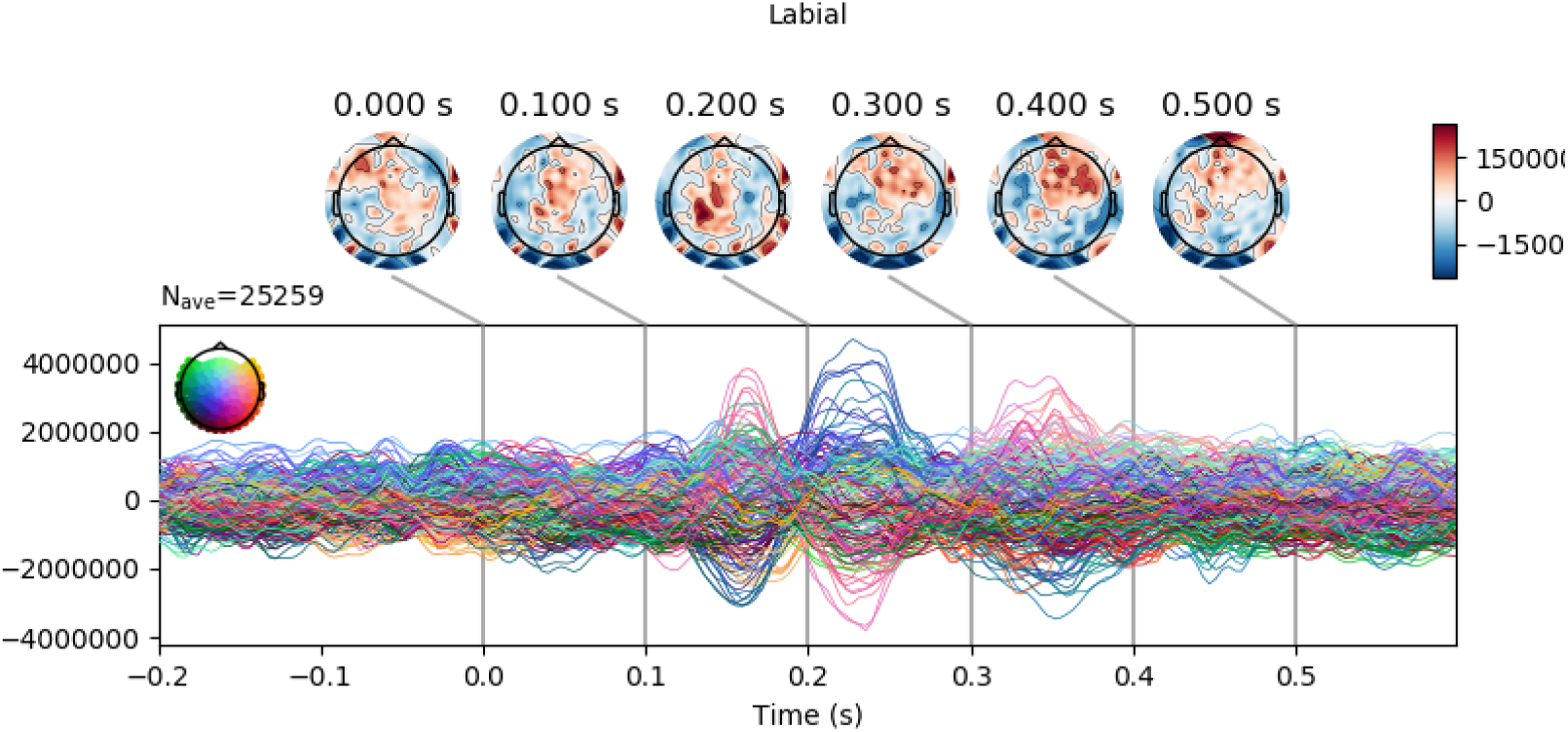
Coefficients for Labial.

**Figure 30:**
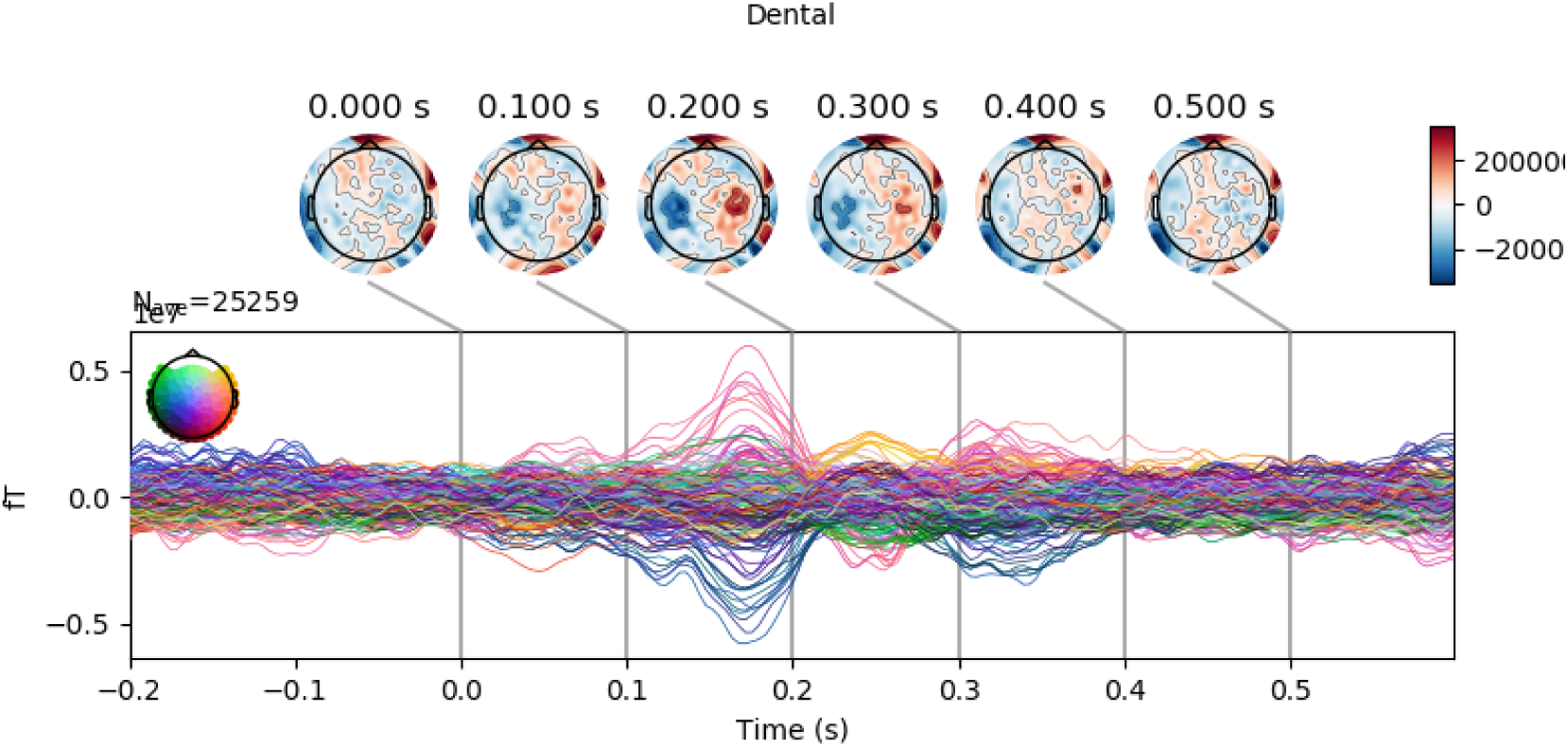
Coefficients for Dental.

**Figure 31:**
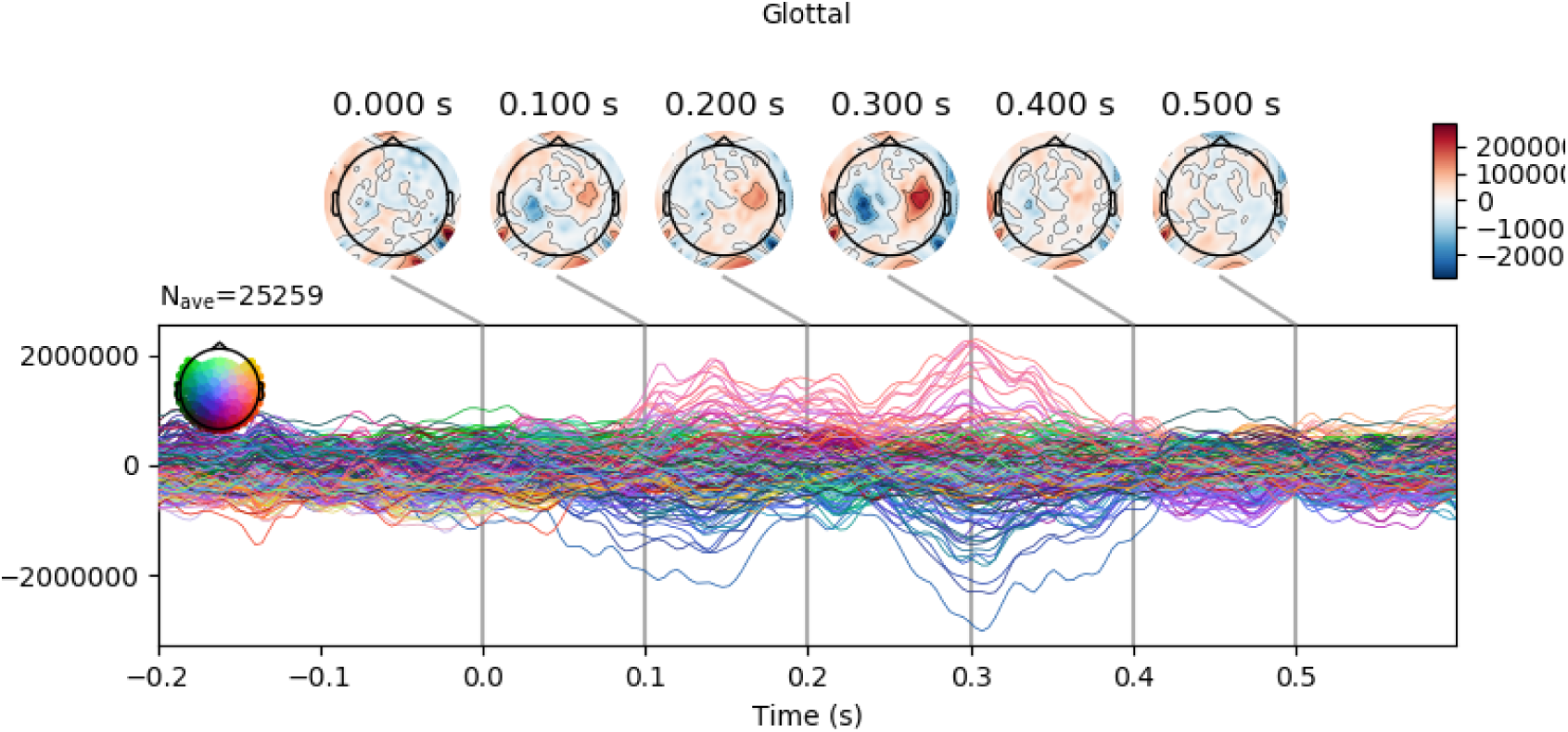
Coefficients for Glottal.

#### 5.10.5. Place of articulation: Vowels

**Figure 32:**
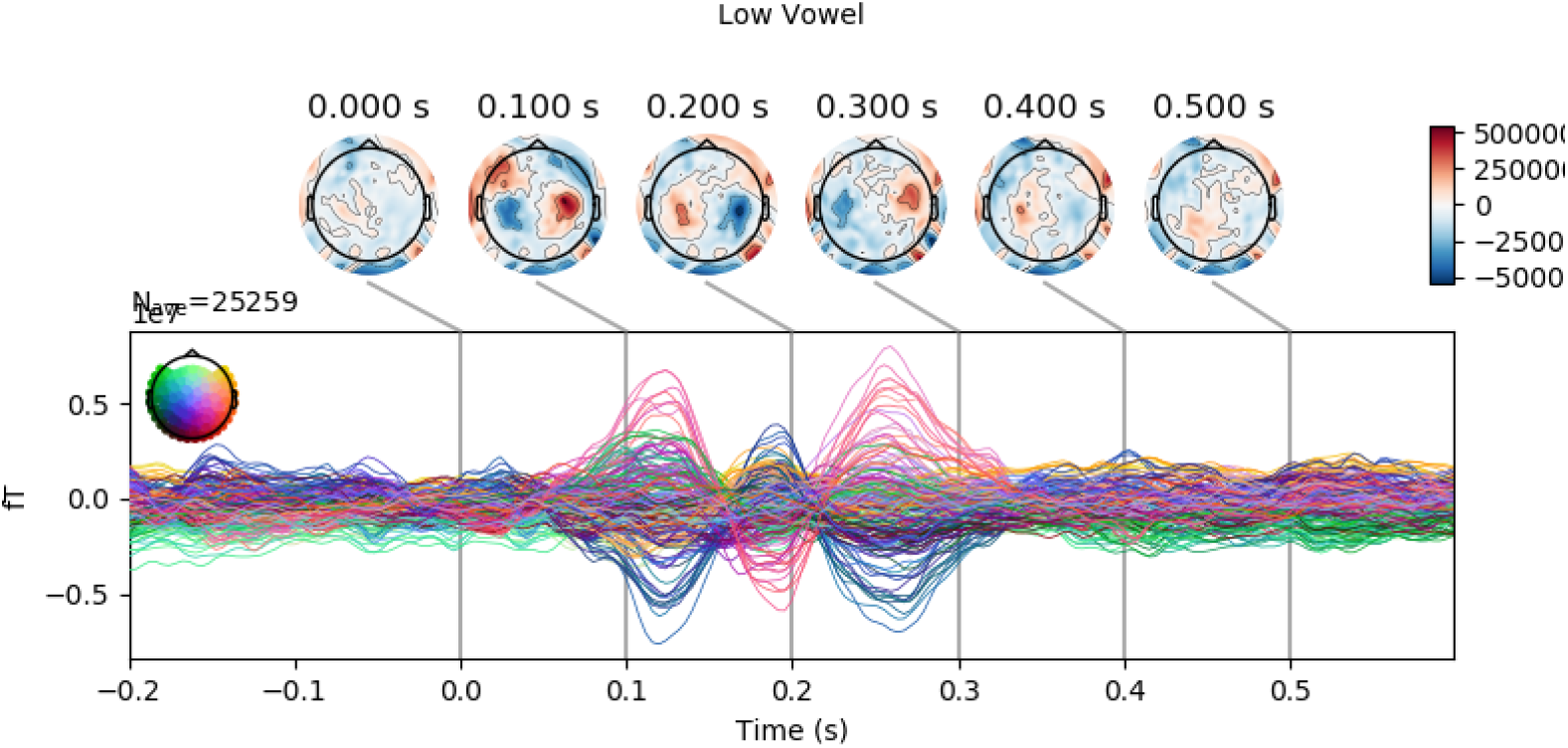
Coefficients for Low Vowel.

**Figure 33:**
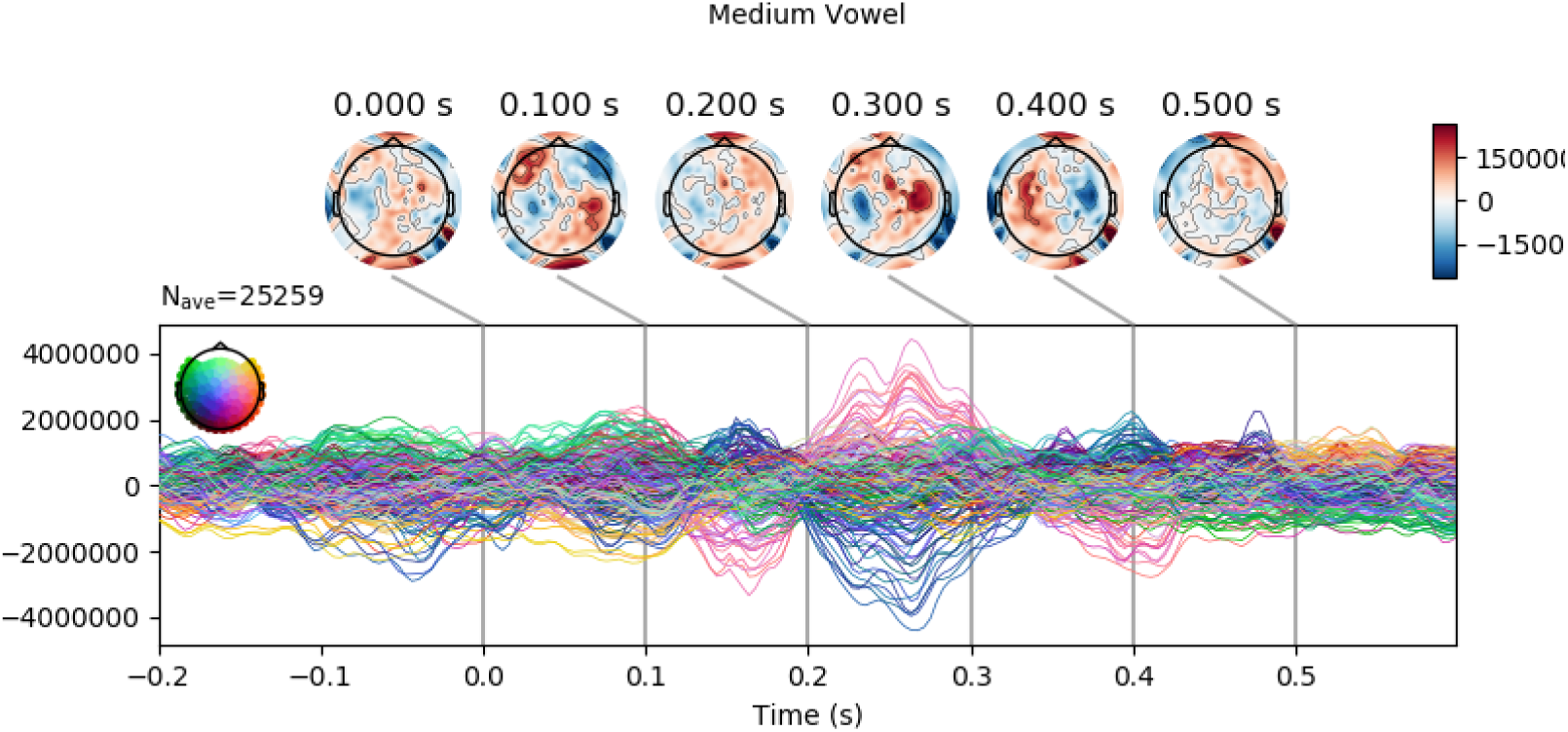
Coefficients for Mid Vowel.

**Figure 34:**
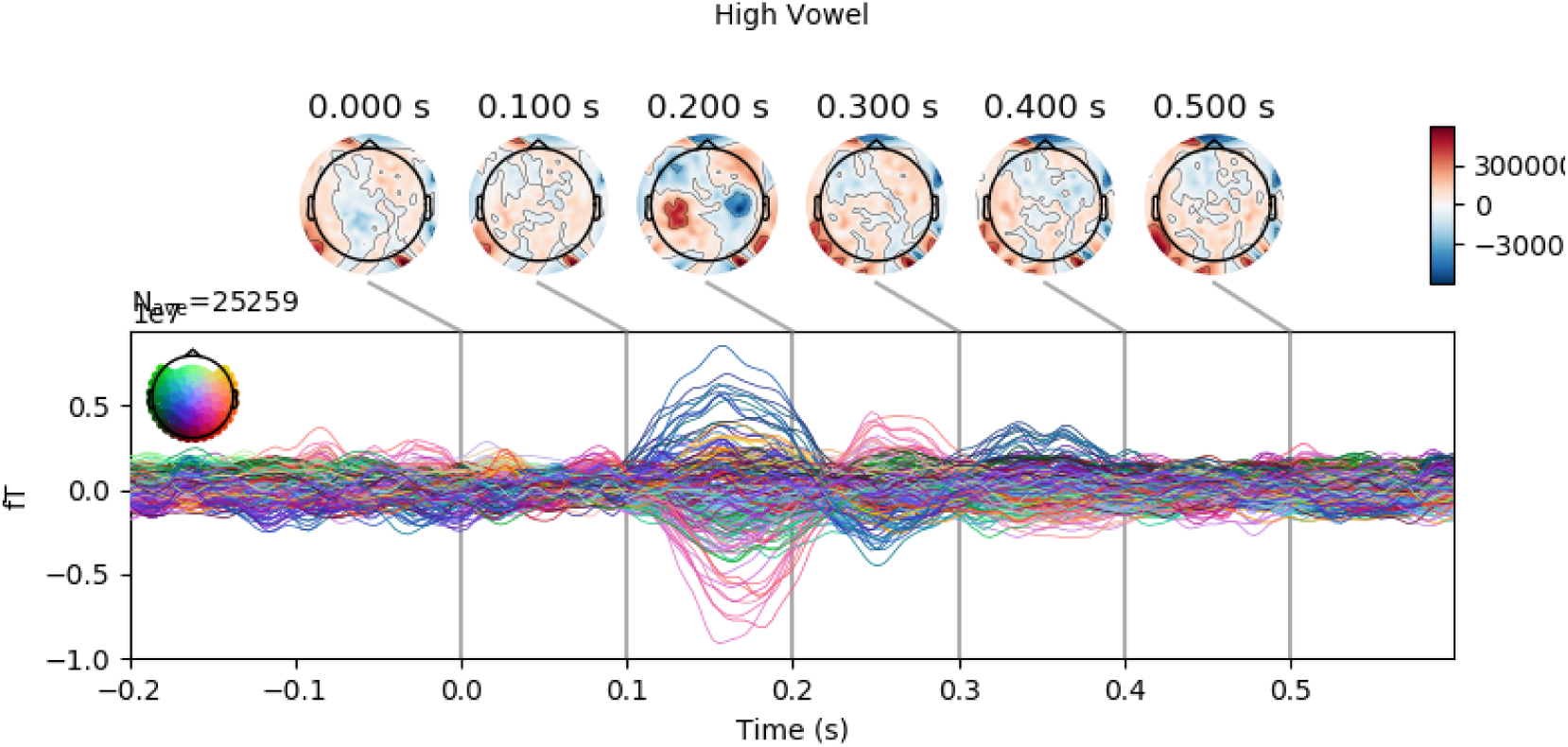
Coefficients for High Vowel.

**Figure 35:**
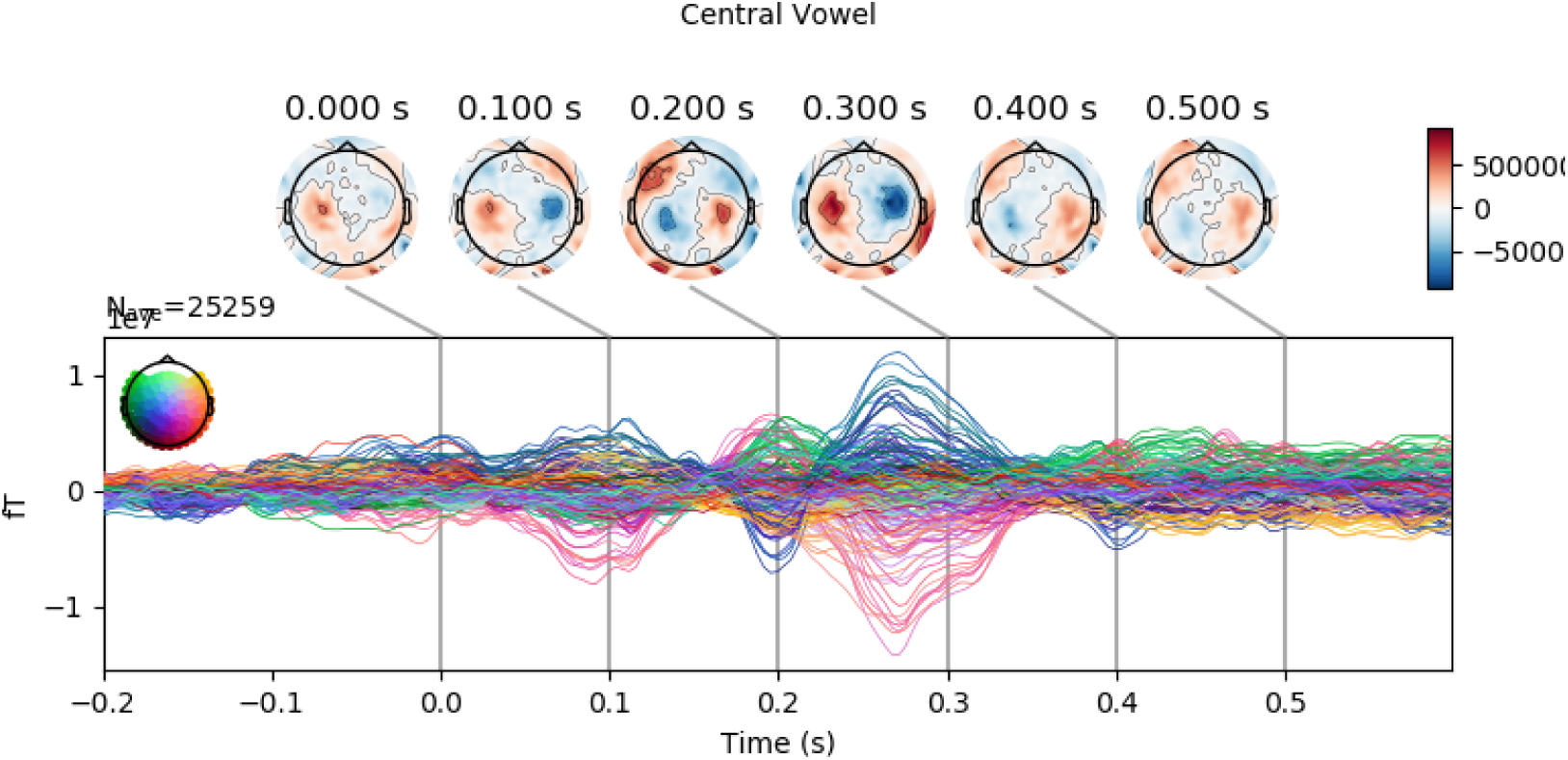
Coefficients for Central Vowel.

#### 5.10.6. Boundary Features

**Figure 36:**
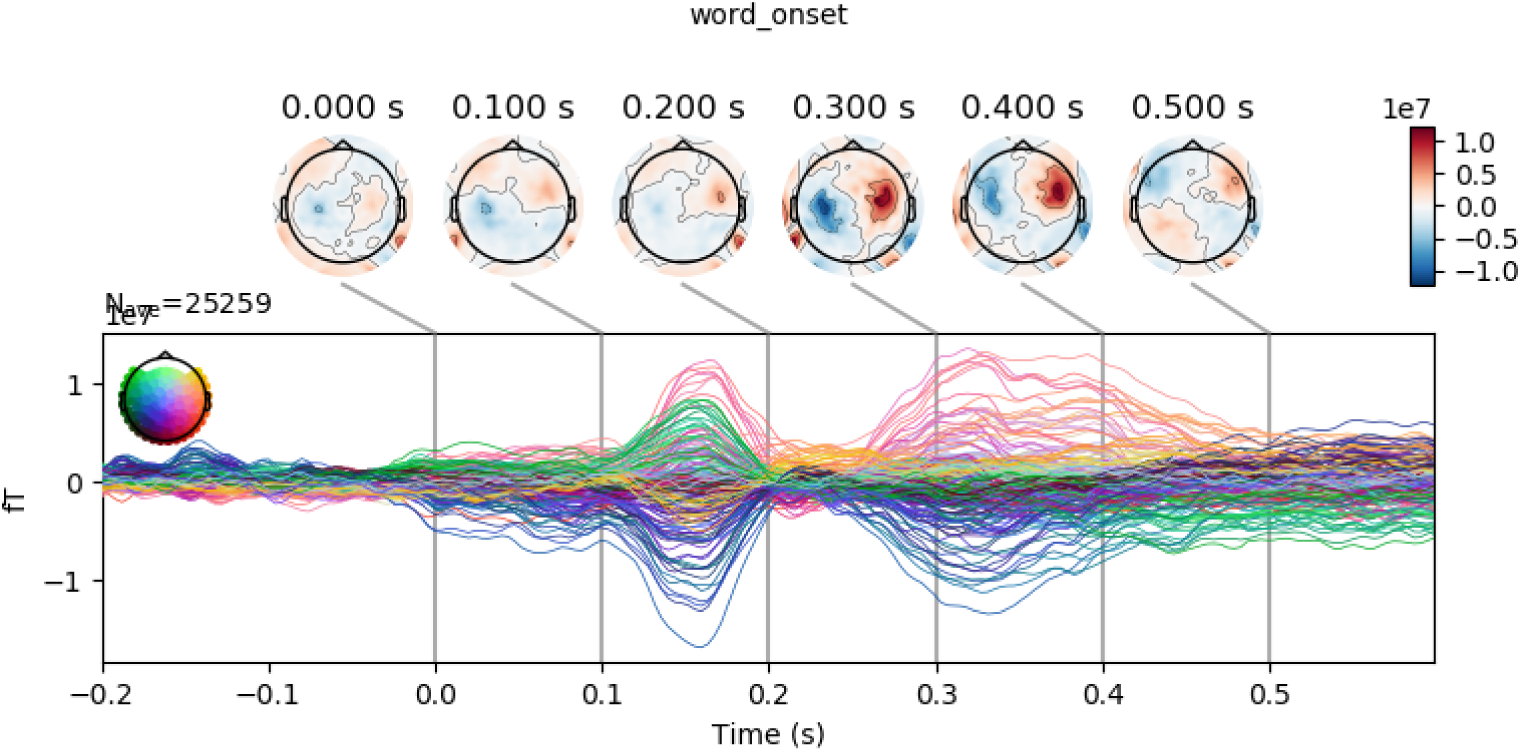
Coefficients for Word Onset.

**Figure 37:**
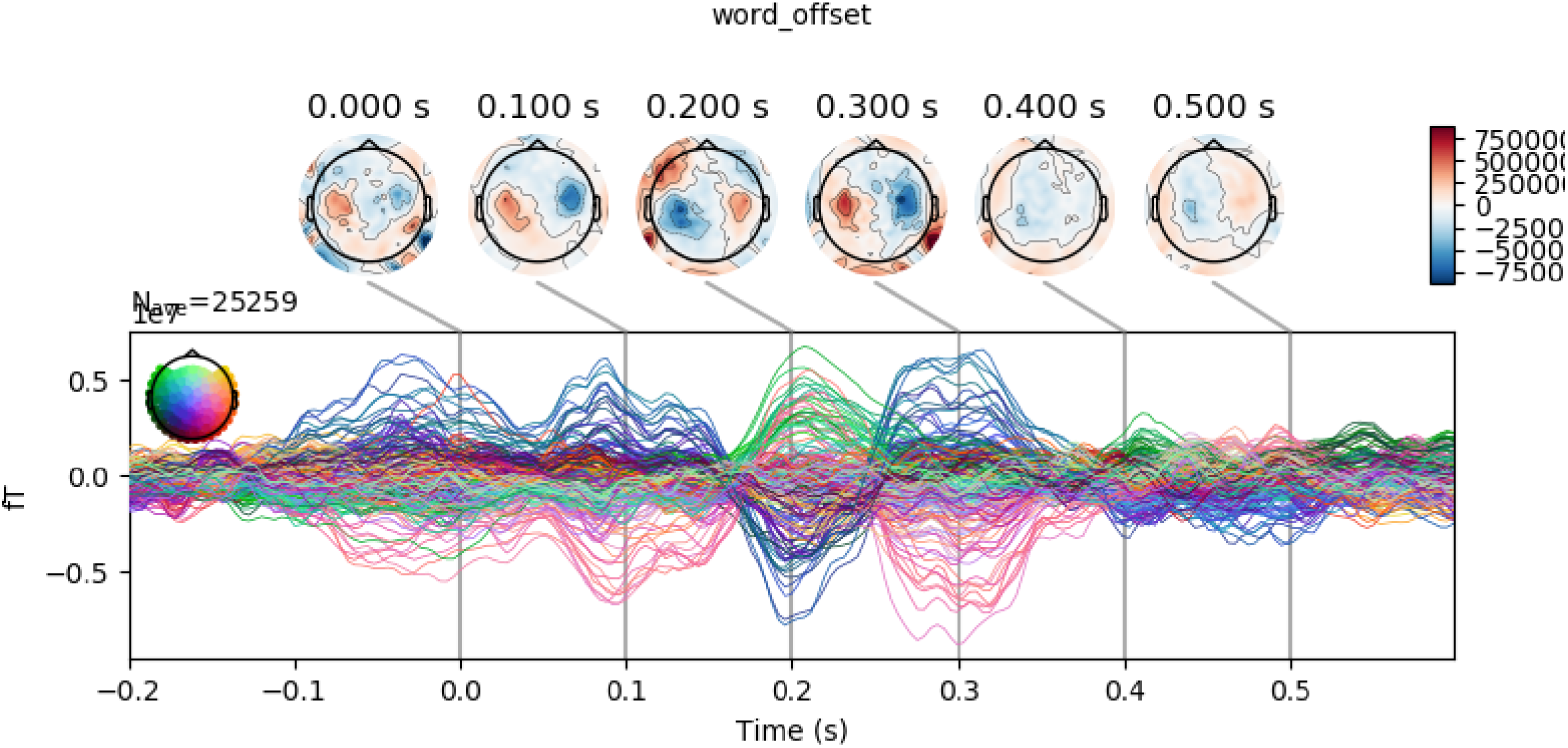
Coefficients for Word Offset.

**Figure 38:**
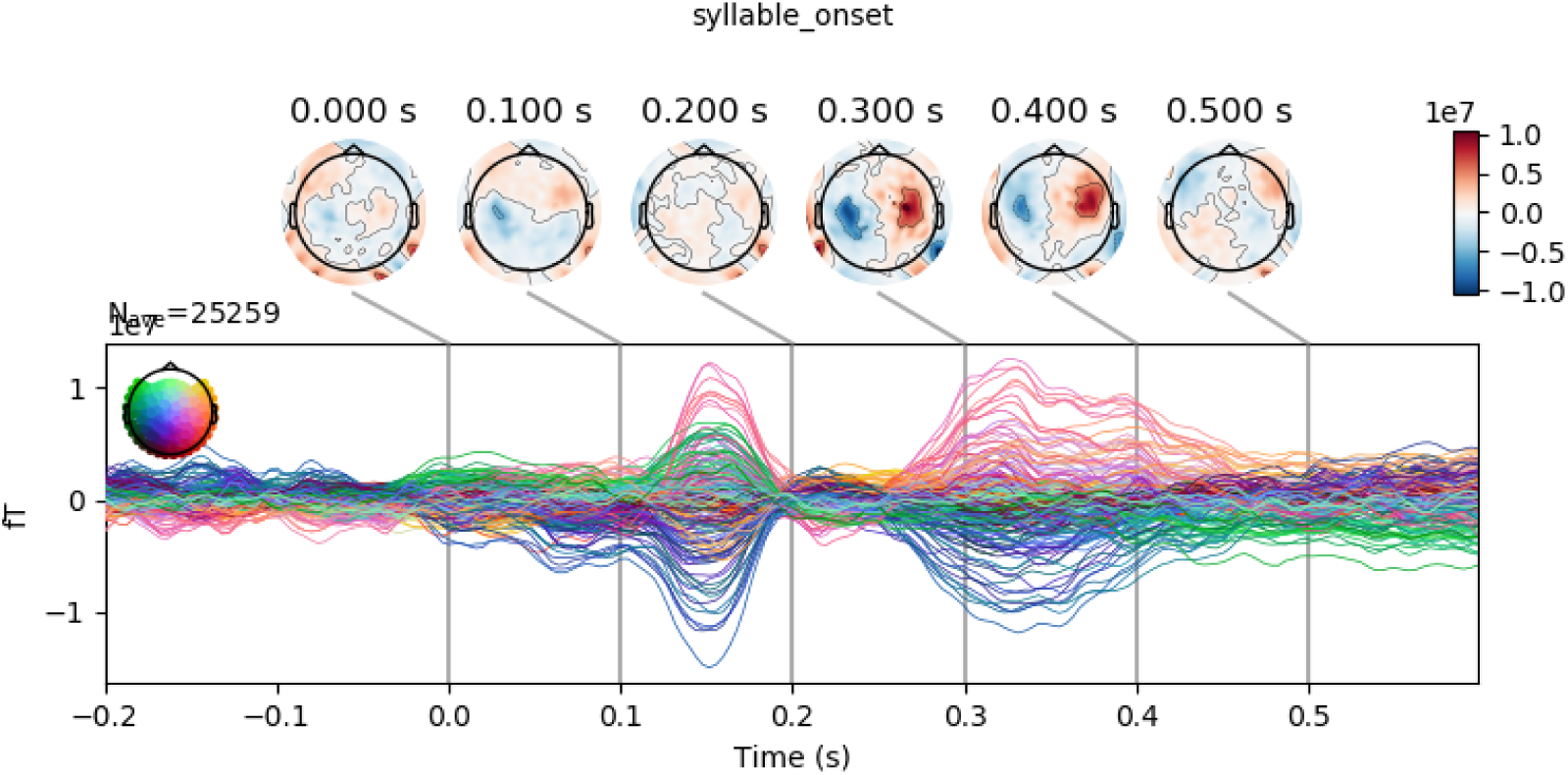
Coefficients for Syllable Onset.

**Figure 39:**
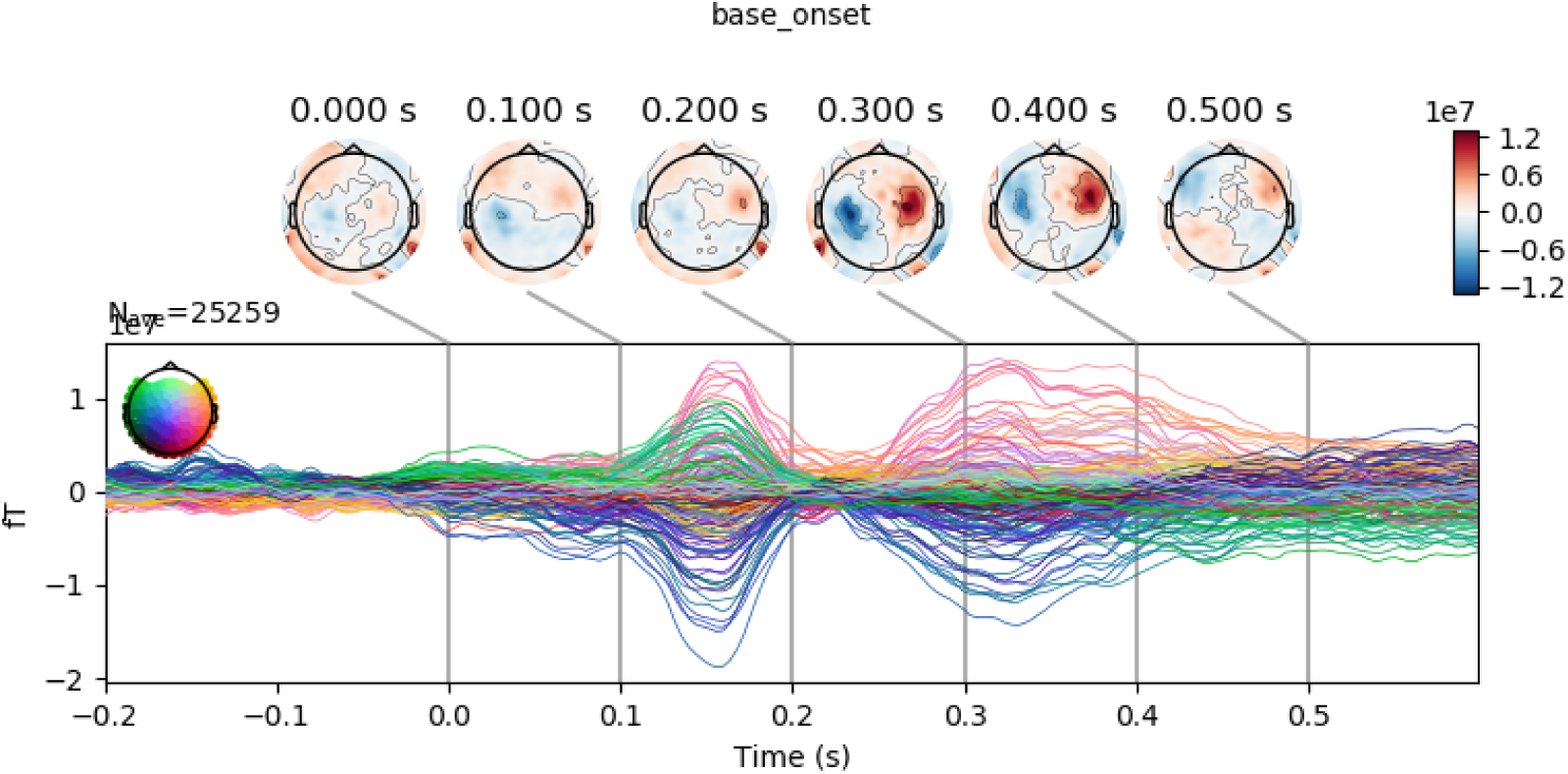
Coefficients for Root Morpheme onset.

**Figure 40:**
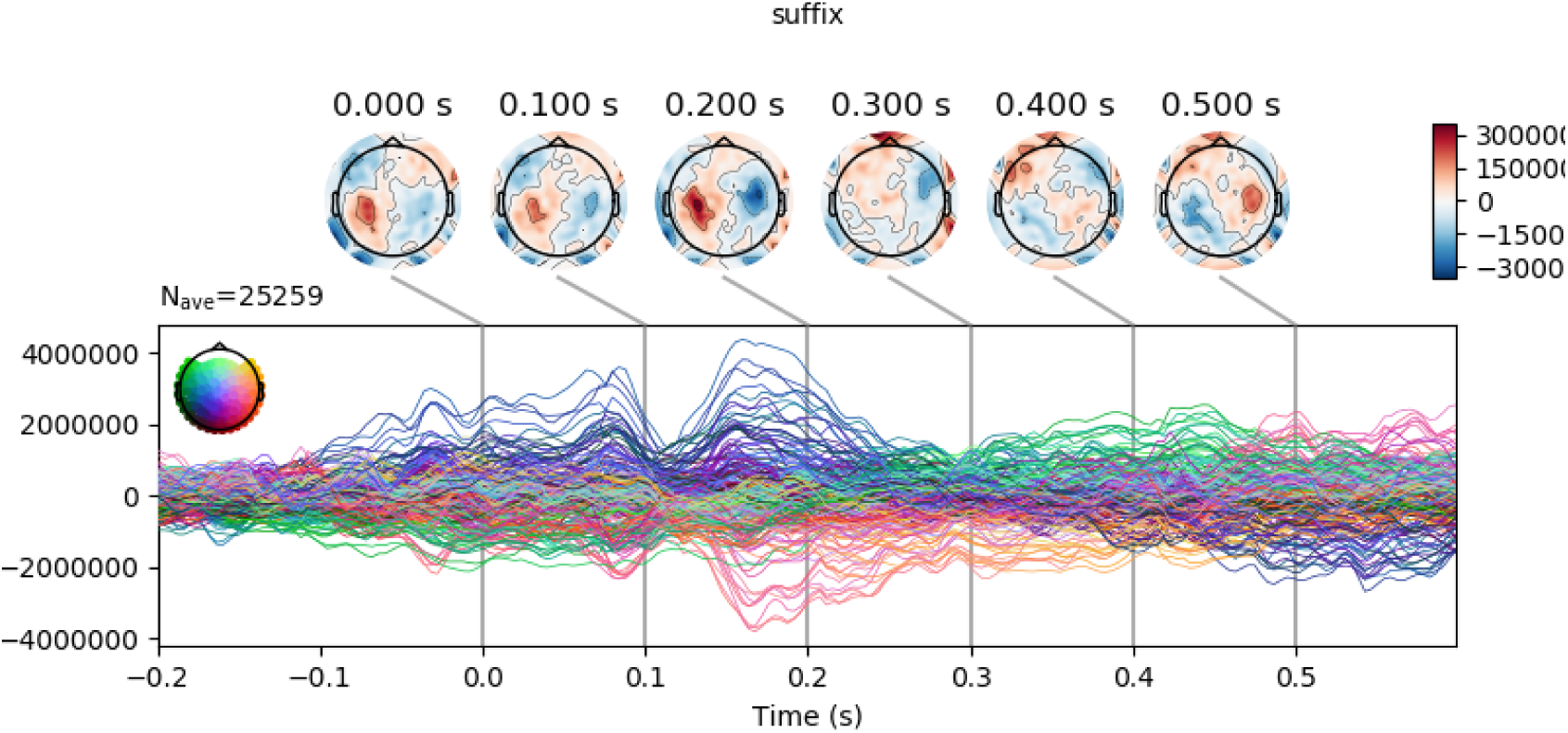
Coefficients for Suffix onset.

**Figure 41:**
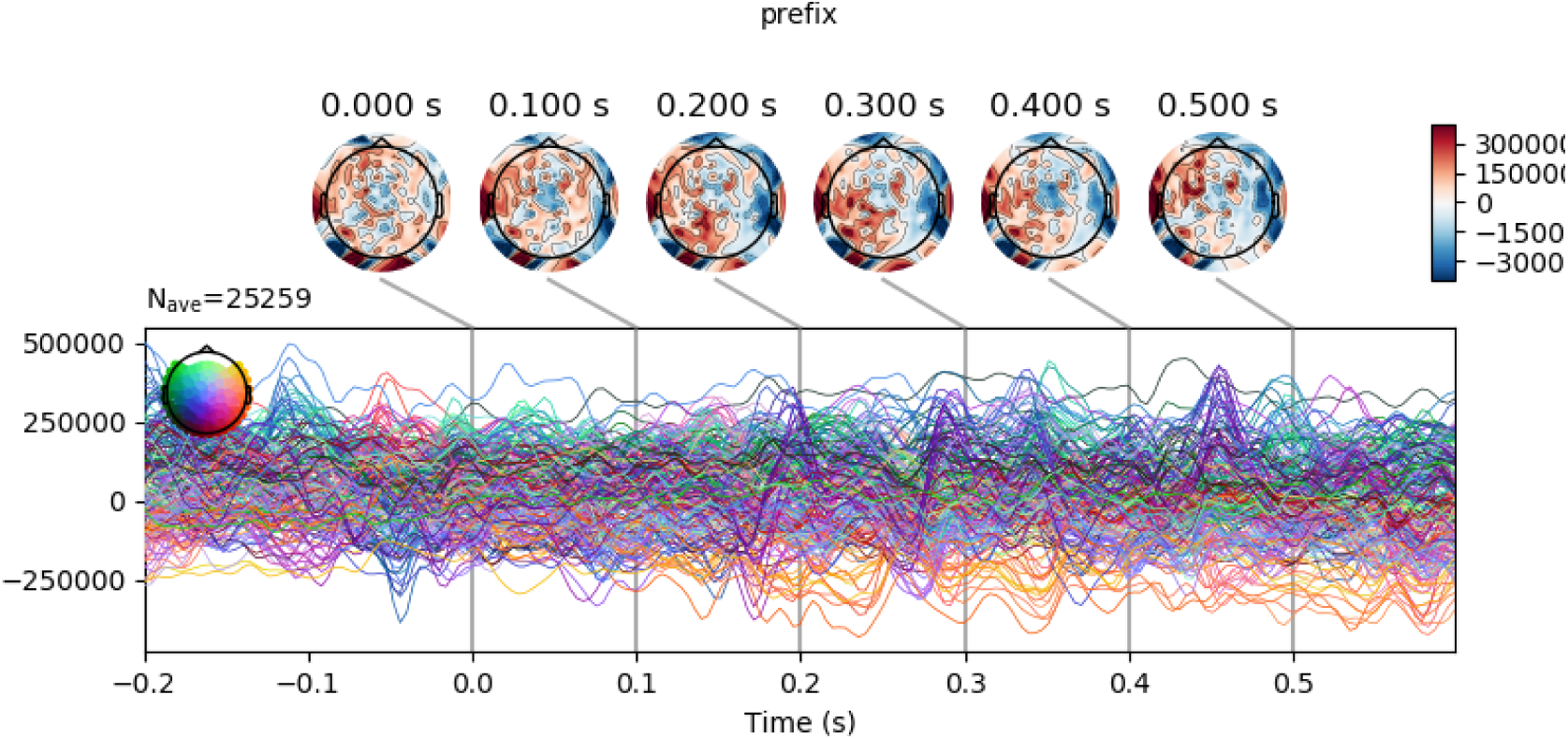
Coefficients for Prefix onset.

#### 5.10.7. Position Features

**Figure 42:**
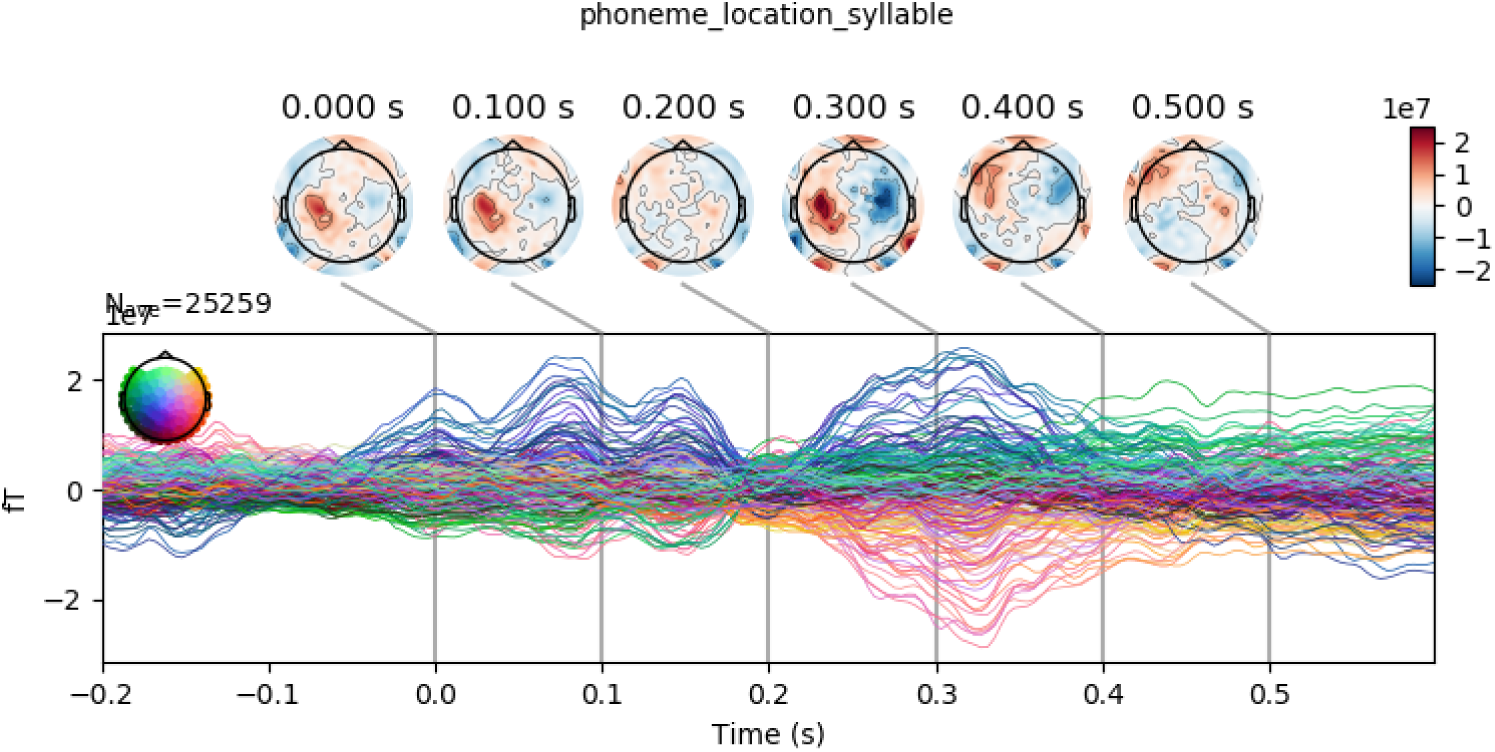
Coefficients for Phoneme location in syllable.

**Figure 43:**
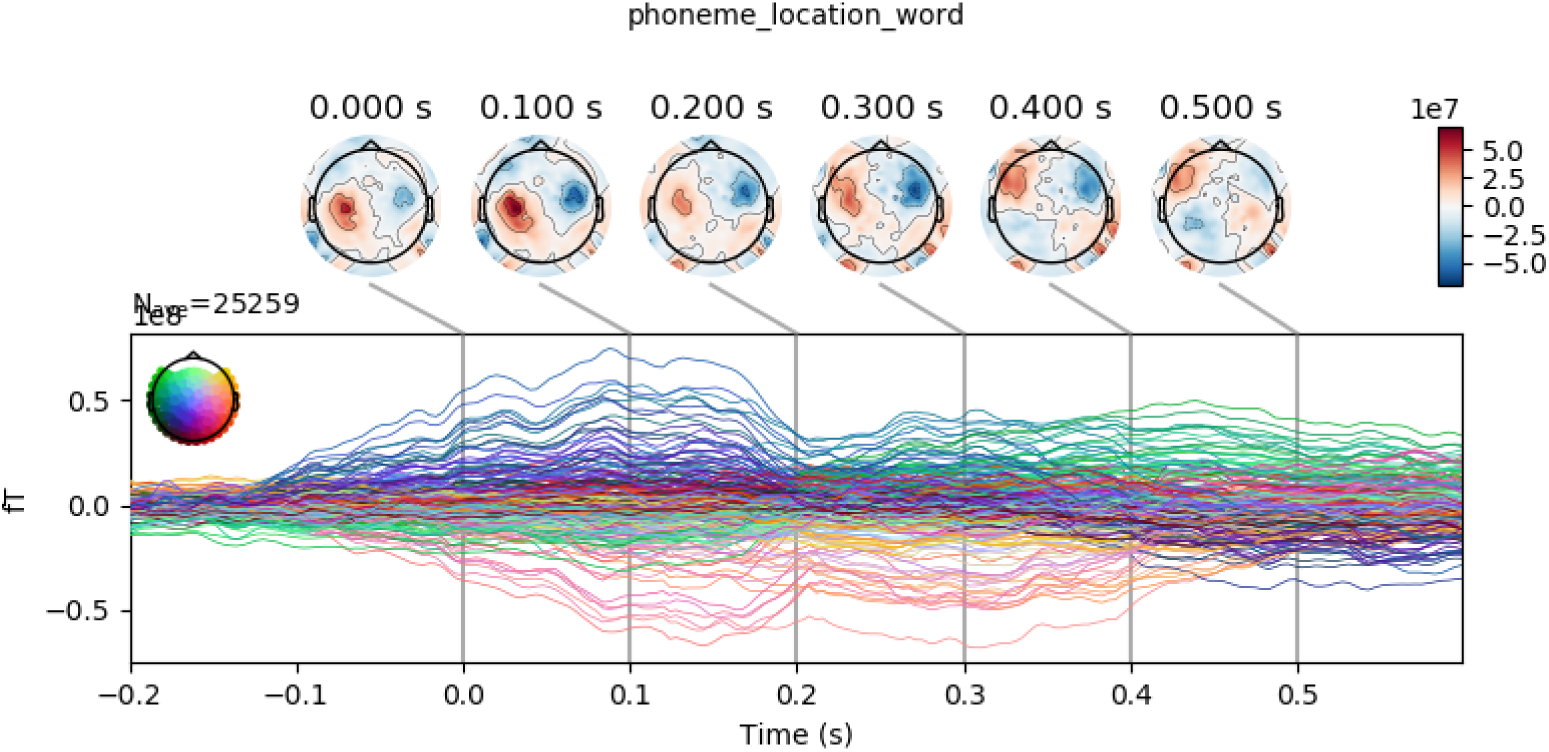
Coefficients for Phoneme location in word.

**Figure 44:**
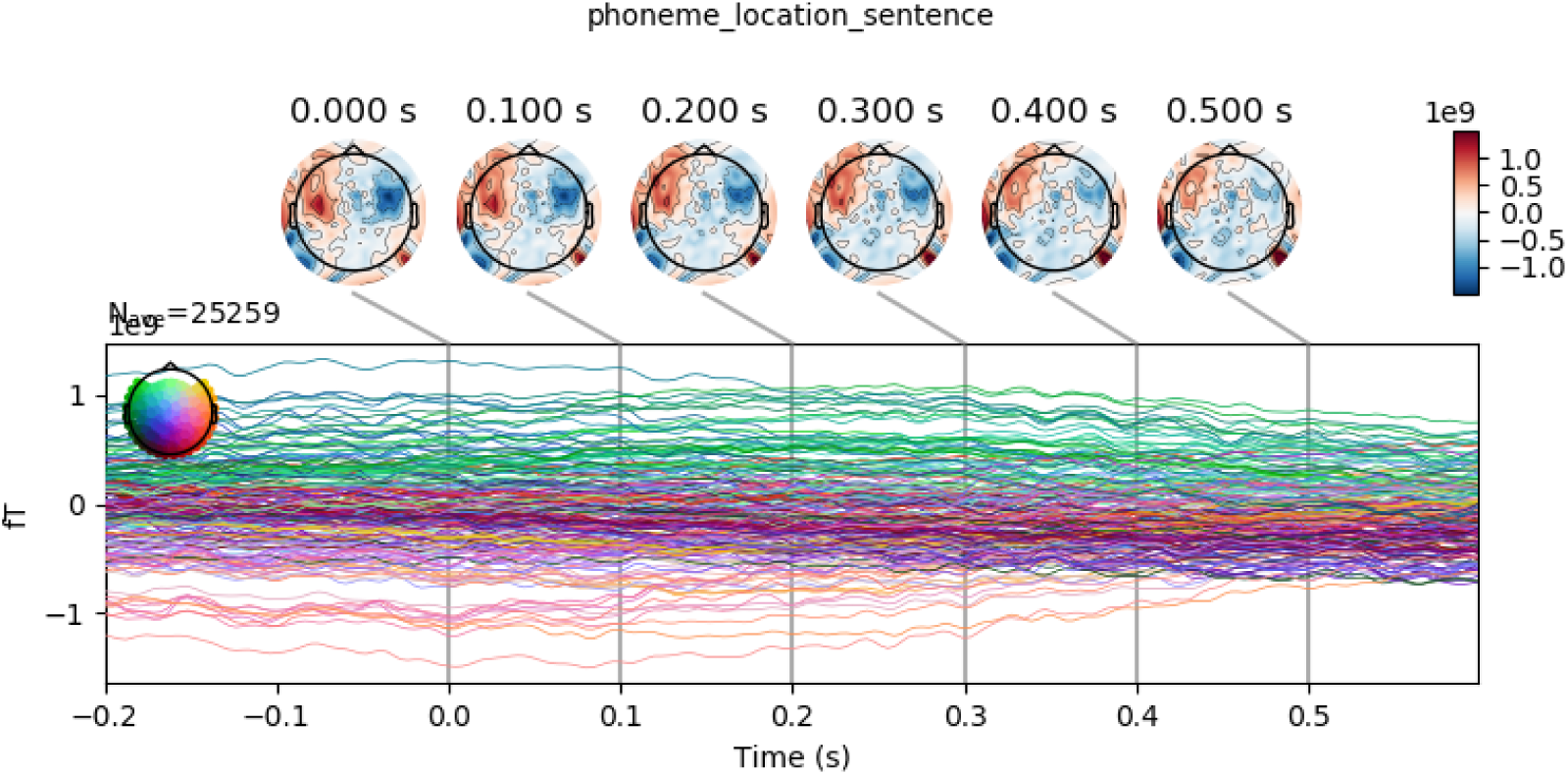
Coefficients for Phoneme location in sentence.

**Figure 45:**
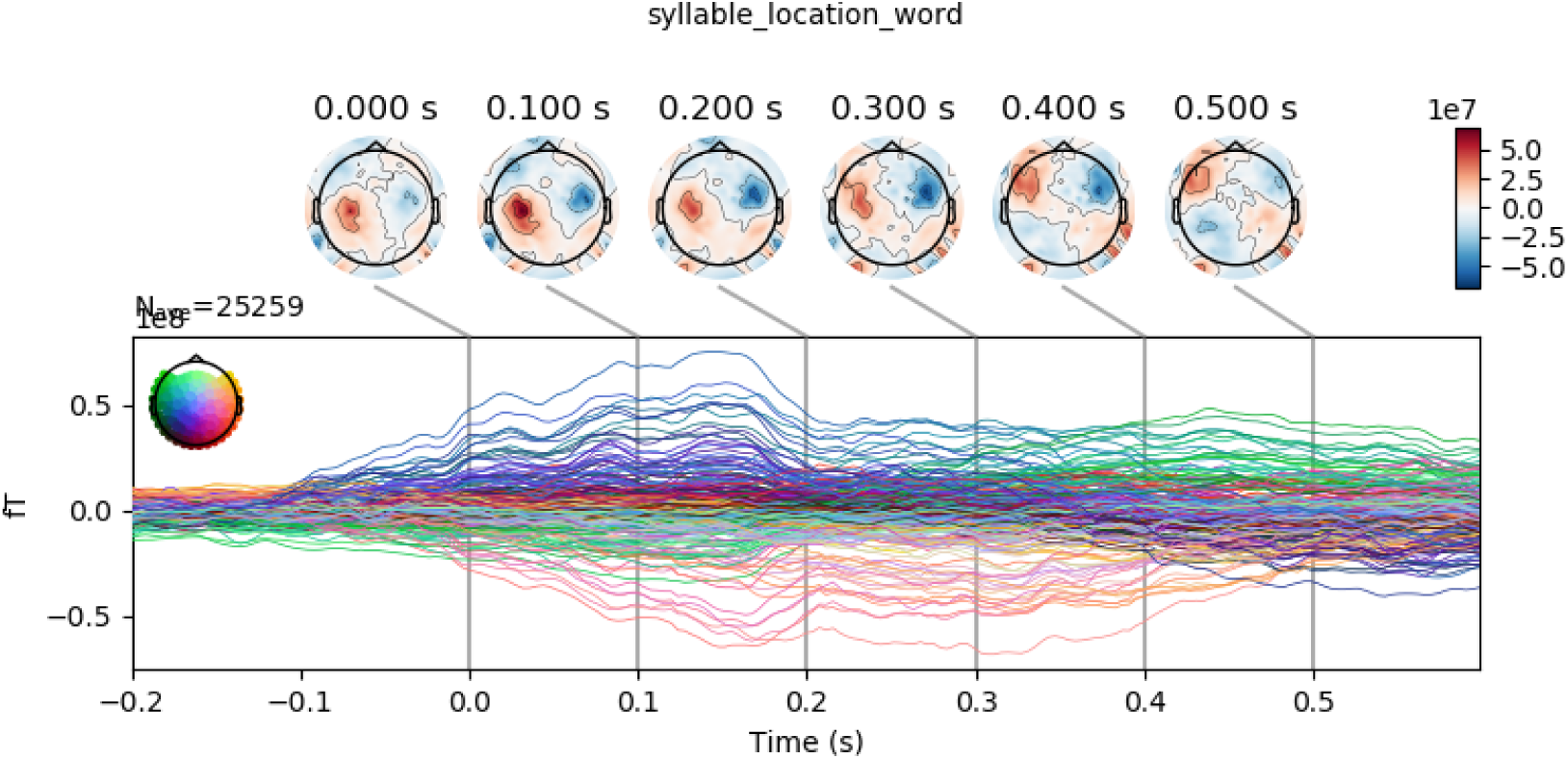
Coefficients for Syllable location in word.

#### 5.10.8. Lexical stress

**Figure 46:**
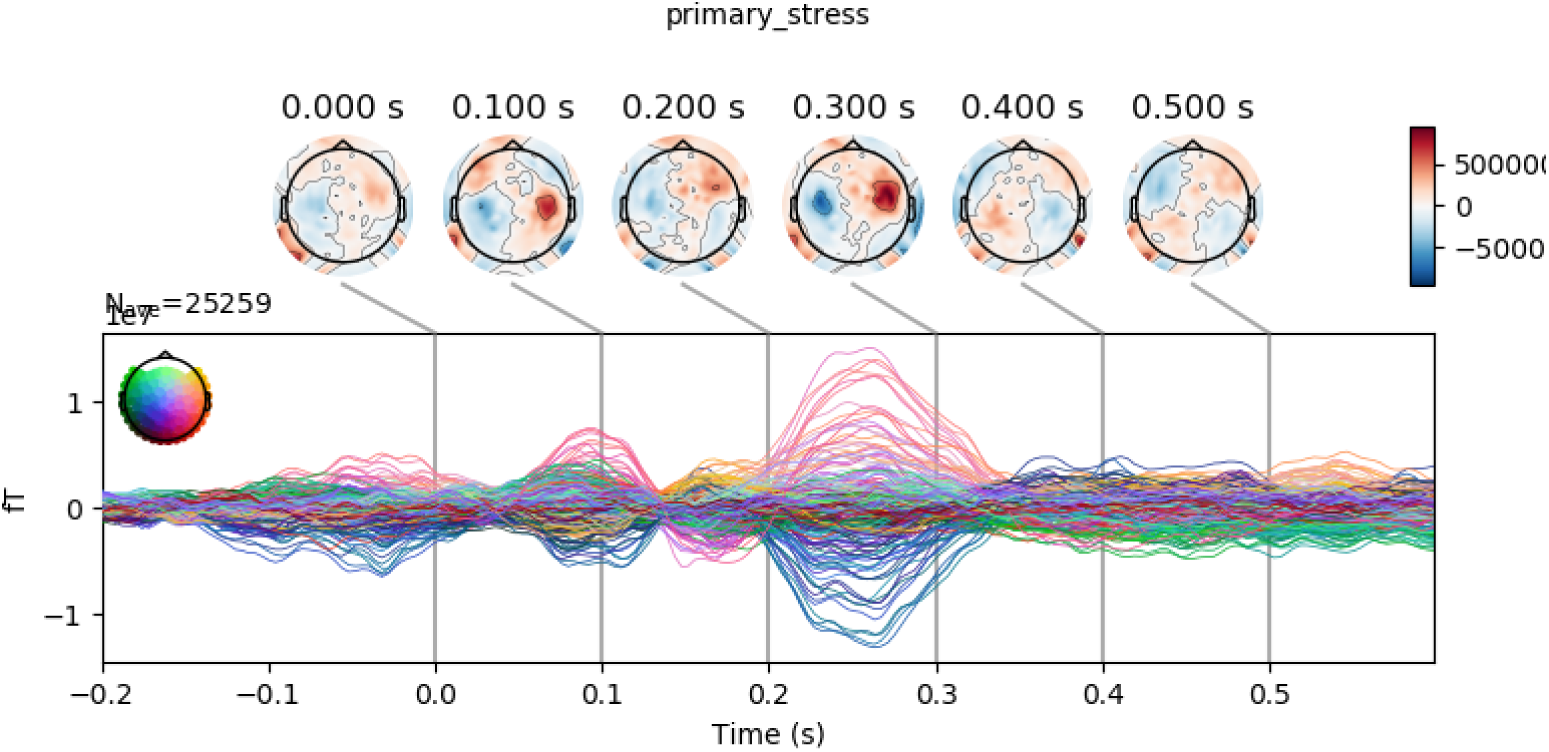
Coefficients for Primary Stress on syllable.

**Figure 47:**
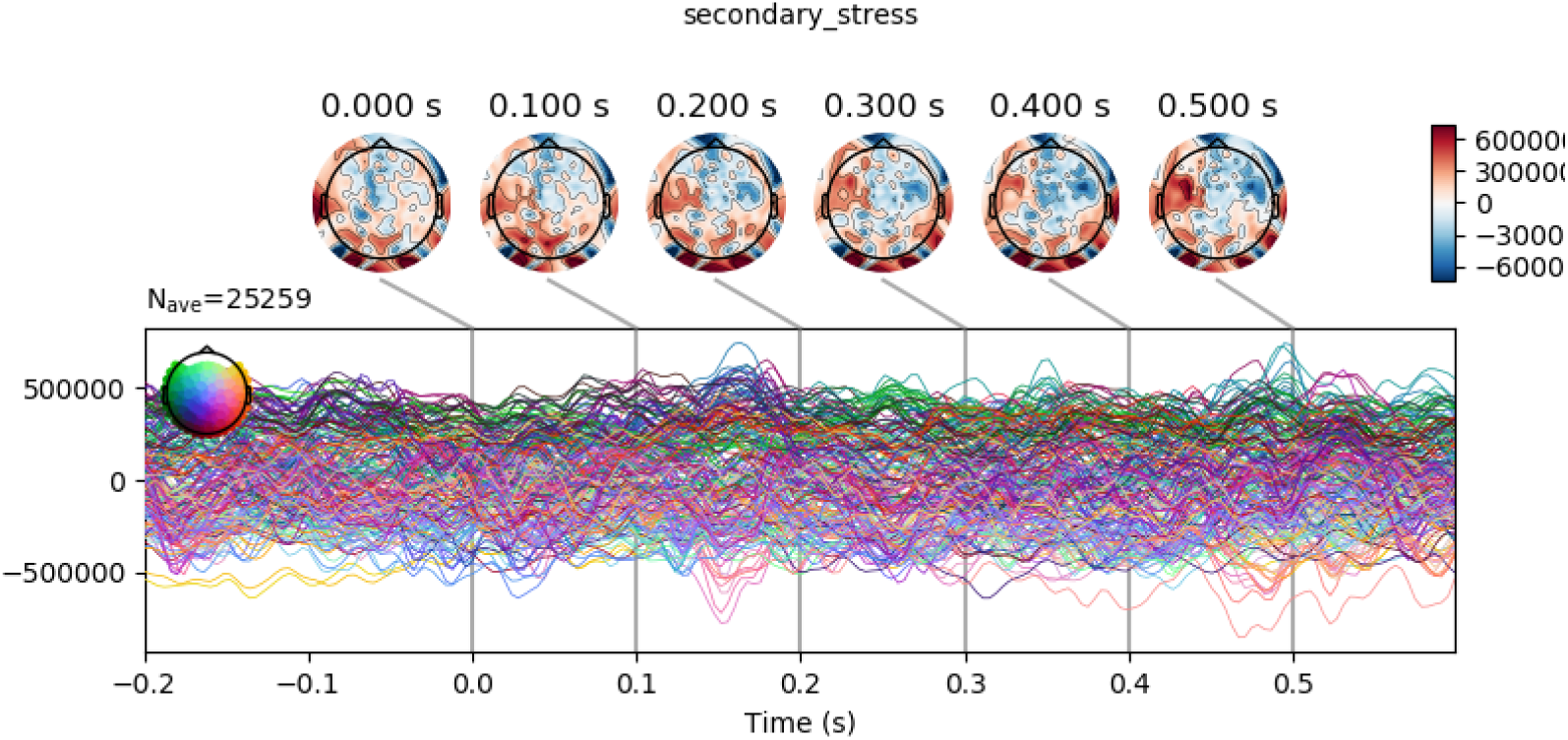
Coefficients for Secondary Stress on syllable.

#### 5.10.9. Statistical Features

**Figure 48:**
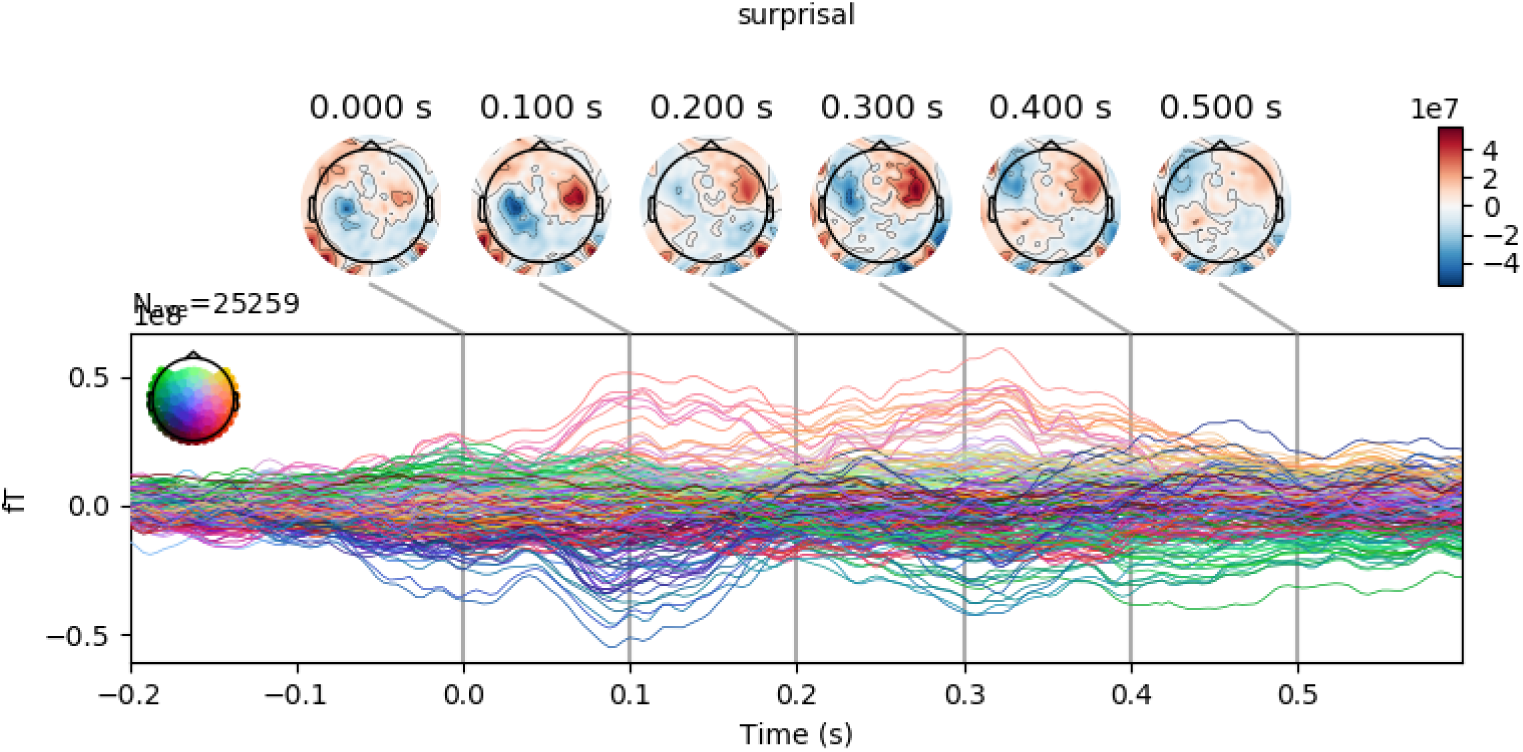
Coefficients for Surprisal.

**Figure 49:**
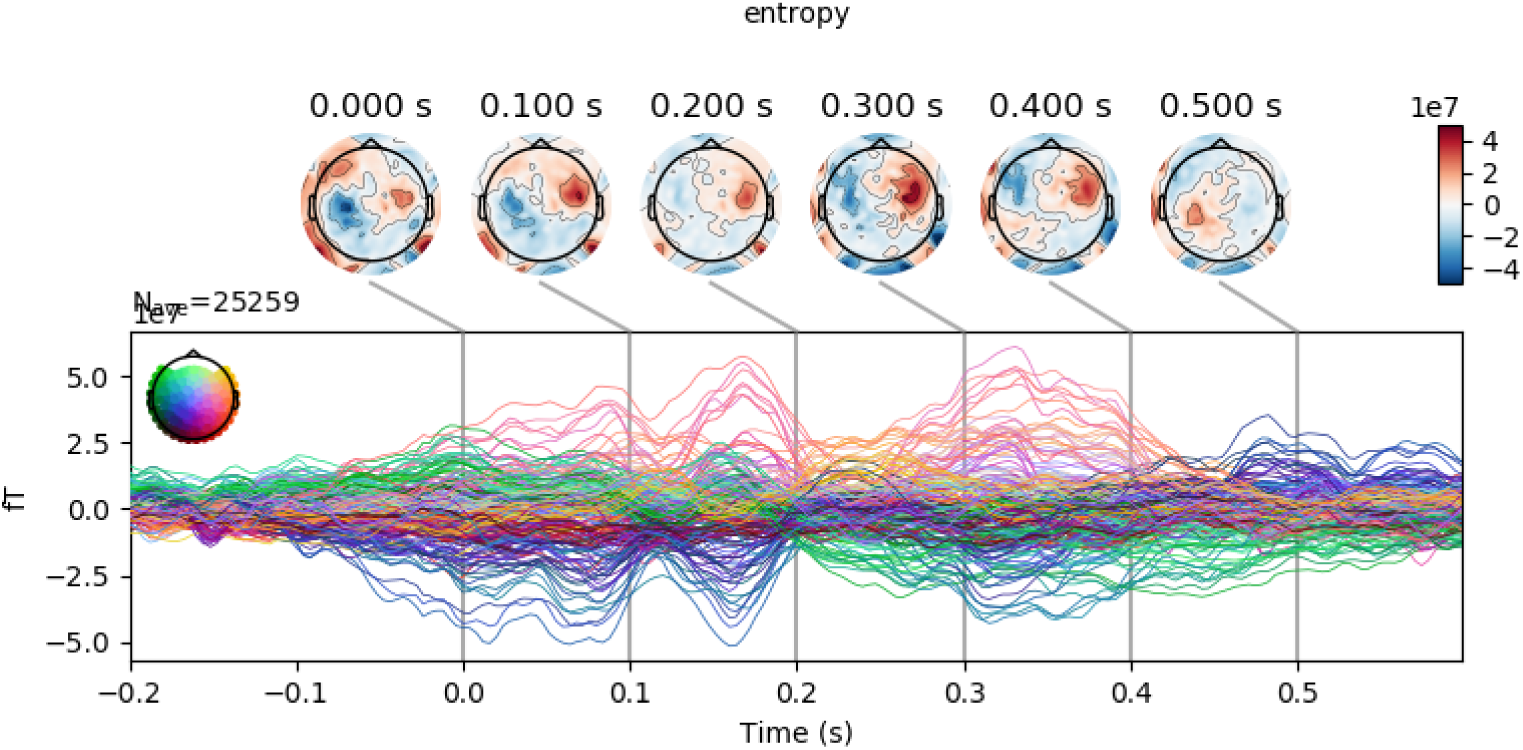
Coefficients for Entropy.

**Figure 50:**
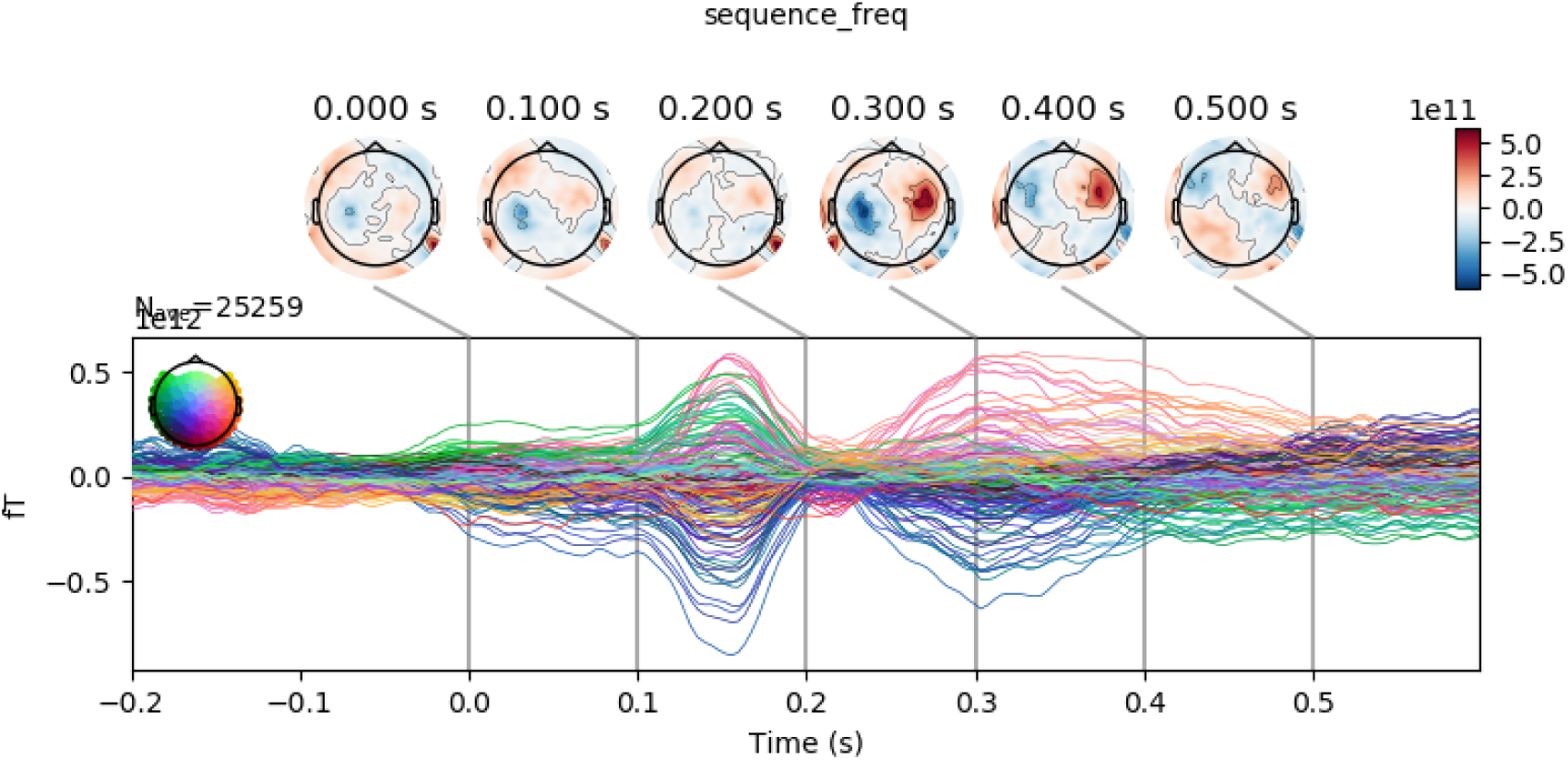
Coefficients for Sequence Frequency.

## Notes

### Competing Interest Statement

The authors have declared no competing interest.

## References

[1] Pisoni, D. B. & Luce, P. A. Acoustic-phonetic representations in word recognition. Cognition 25, 21–52 (1987).

[2] Wöstmann, M., Fiedler, L. & Obleser, J. Tracking the signal, cracking the code: Speech and speech comprehension in non-invasive human electrophysiology. Language, Cognition and Neuroscience 32, 855–869 (2017).

[3] Benzeghiba, M. et al. Automatic speech recognition and speech variability: A review. Speech communication 49, 763–786 (2007).

[4] Baevski, A., Zhou, H., Mohamed, A. & Auli, M. wav2vec 2.0: A framework for self-supervised learning of speech representations. arXiv preprint 2006.11477 (2020).

[5] Millet, J. & King, J.-R. Inductive biases, pretraining and fine-tuning jointly account for brain responses to speech. arXiv preprint 2103.01032 (2021).

[6] Marslen-Wilson, W. D. & Welsh, A. Processing interactions and lexical access during word recognition in continuous speech. Cognitive psychology 10, 29–63 (1978).

[7] McClelland, J. L. & Elman, J. L. The trace model of speech perception. Cognitive psychology 18, 1–86 (1986).

[8] Norris, D. Shortlist: A connectionist model of continuous speech recognition. Cognition 52, 189–234 (1994).

[9] Mesgarani, N., Cheung, C., Johnson, K. & Chang, E. F. Phonetic feature encoding in human superior temporal gyrus. Science 343, 1006–1010 (2014).

[10] Chang, E. F. et al. Categorical speech representation in human superior temporal gyrus. Nature neuroscience 13, 1428 (2010).

[11] Khalighinejad, B., da Silva, G. C. & Mesgarani, N. Dynamic encoding of acoustic features in neural responses to continuous speech. Journal of Neuroscience 37, 2176–2185 (2017).

[12] Yi, H. G., Leonard, M. K. & Chang, E. F. The encoding of speech sounds in the superior temporal gyrus. Neuron 102, 1096–1110 (2019).

[13] Gwilliams, L. & Marantz, A. Non-linear processing of a linear speech stream: The influence of morphological structure on the recognition of spoken arabic words. Brain and language 147, 1–13 (2015).

[14] Leonard, M. K., Baud, M. O., Sjerps, M. J. & Chang, E. F. Perceptual restoration of masked speech in human cortex. Nature communications 7, 1–9 (2016).

[15] Brodbeck, C., Hong, L. E. & Simon, J. Z. Rapid transformation from auditory to linguistic representations of continuous speech. Current Biology 28, 3976–3983 (2018).

[16] Gwilliams, L., Linzen, T., Poeppel, D. & Marantz, A. In spoken word recognition, the future predicts the past. Journal of Neuroscience 38, 7585–7599 (2018).

[17] Kell, A. J. & McDermott, J. H. Deep neural network models of sensory systems: windows onto the role of task constraints. Current opinion in neurobiology 55, 121–132 (2019).

[18] Picton, T. W., Woods, D. L., Baribeau-Braun, J. & Healey, T. M. Evoked potential audiometry. J Otolaryngol 6, 90–119 (1977).

[19] Näätänen, R. & Picton, T. The n1 wave of the human electric and magnetic response to sound: a review and an analysis of the component structure. Psychophysiology 24, 375–425 (1987).

[20] Crosse, M. J., Di Liberto, G. M., Bednar, A. & Lalor, E. C. The multivariate temporal response function (mtrf) toolbox: a matlab toolbox for relating neural signals to continuous stimuli. Frontiers in human neuroscience 10, 604 (2016).

[21] King, J.-R., Charton, F., Lopez-Paz, D. & Oquab, M. Back-to-back regression: Disentangling the influence of correlated factors from multivariate observations. NeuroImage 220, 117028 (2020).

[22] Robles, L. & Ruggero, M. A. Mechanics of the mammalian cochlea. Physiological reviews 81, 1305–1352 (2001).

[23] De-Wit, L., Alexander, D., Ekroll, V. & Wagemans, J. Is neuroimaging measuring information in the brain? Psychonomic bulletin & review 23, 1415–1428 (2016).

[24] King, J. & Dehaene, S. Characterizing the dynamics of mental representations: the temporal generalization method. Trends in cognitive sciences 18, 203–210 (2014).

[25] King, J. & Wyart, V. The human brain encodes a chronicle of visual events at each instant of time thanks to the multiplexing of traveling waves. Journal of Neuroscience (2021).

[26] Gwilliams, L. & Davis, M. Extracting language content from speech sounds: An information theoretic approach (2020).

[27] Daube, C., Ince, R. A. & Gross, J. Simple acoustic features can explain phoneme-based predictions of cortical responses to speech. Current Biology 29, 1924–1937 (2019).

[28] Wickelgren, W. A. Short-term memory for phonemically similar lists. The American Journal of Psychology 78, 567–574 (1965).

[29] Glasspool, D. W. & Houghton, G. Serial order and consonant–vowel structure in a graphemic output buffer model. Brain and language 94, 304–330 (2005).

[30] Fischer-Baum, S. A common representation of serial position in language and memory. In Psychology of Learning and Motivation, vol. 68, 31–54 (Elsevier, 2018).

[31] Ohala, J. J., Browman, C. P. & Goldstein, L. M. Towards an articulatory phonology. Phonology 3, 219–252 (1986).

[32] Browman, C. P. & Goldstein, L. Articulatory gestures as phonological units. Phonology 6, 201–251 (1989).

[33] MacDonald, C. J., Lepage, K. Q., Eden, U. T. & Eichenbaum, H. Hippocampal “time cells” bridge the gap in memory for discontiguous events. Neuron 71, 737–749 (2011).

[34] Sohoglu, E., Peelle, J. E., Carlyon, R. P. & Davis, M. H. Predictive top-down integration of prior knowledge during speech perception. Journal of Neuroscience 32, 8443–8453 (2012).

[35] Bendixen, A., Scharinger, M., Strauss, A. & Obleser, J. Prediction in the service of comprehension: modulated early brain responses to omitted speech segments. Cortex 53, 9–26 (2014).

[36] Halle, M. & Stevens, K. Speech recognition: A model and a program for research. IRE transactions on information theory 8, 155–159 (1962).

[37] Gagnepain, P., Henson, R. N. & Davis, M. H. Temporal predictive codes for spoken words in auditory cortex. Current Biology 22, 615–621 (2012).

[38] Gwilliams, L., Poeppel, D., Marantz, A. & Linzen, T. Phonological (un) certainty weights lexical activation. arXiv preprint 1711.06729 (2017).

[39] Di Liberto, G. M., Wong, D., Melnik, G. A. & de Cheveigné, A. Low-frequency cortical responses to natural speech reflect probabilistic phonotactics. Neuroimage 196, 237–247 (2019).

[40] Gold, J. I. & Shadlen, M. N. The neural basis of decision making. Annual review of neuroscience 30 (2007).

[41] Ide, N. & Macleod, C. The american national corpus: A standardized resource of american english. In Proceedings of corpus linguistics, vol. 3, 1–7 (Lancaster University Centre for Computer Corpus Research on Language …, 2001).

[42] Yuan, J. & Liberman, M. Speaker identification on the scotus corpus. Journal of the Acoustical Society of America 123, 3878 (2008).

[43] Gramfort, A. et al. Mne software for processing meg and eeg data. Neuroimage 86, 446–460 (2014).

[44] King, S. & Taylor, P. Detection of phonological features in continuous speech using neural networks (2000).

[45] Balota, D. A. et al. The english lexicon project. Behavior research methods 39, 445–459 (2007).

[46] King, J.-R. et al. Encoding and decoding neuronal dynamics: Methodological framework to uncover the algorithms of cognition (2018).

[47] King, J.-R., Charton, F., Lopez-Paz, D. & Oquab, M. Discriminating the influence of correlated factors from multivariate observations: the back-to-back regression. bioRxiv (2020).

[48] Pedregosa, F. et al. Scikit-learn: Machine learning in python. Journal of machine learning research 12, 2825–2830 (2011).

